# A unified internal model theory to resolve the paradox of active versus passive self-motion sensation

**DOI:** 10.1101/132936

**Authors:** Jean Laurens, Dora Angelaki

## Abstract

Brainstem and cerebellar neurons implement an internal model to accurately estimate self-motion during externally-generated (‘passive’) movements. However, these neurons show reduced responses during self-generated (‘active’) movements, indicating that the brain computes the predicted sensory consequences of motor commands in order to cancel sensory signals. Remarkably, the computational processes underlying sensory prediction during active motion and their relationship to internal model computations established during passive movements remain unknown. Here we construct a Kalman filter that incorporates motor commands into a previously-established model of optimal passive self-motion estimation. We find that the simulated sensory error and feedback signals match experimentally measured neuronal response during active and passive head and trunk rotations and translations. We conclude that a single internal model of head motion can process motor commands and sensory afferent signals optimally, and we describe how previously identified neural responses in the brainstem and cerebellum may represent distinct nodes in these computations.

## Introduction

For many decades, research on vestibular function has used passive motion stimuli generated by rotating chairs, motion platforms or centrifuges to characterize the responses of the vestibular motion sensors in the inner ear and the subsequent stages of neuronal processing. This research has revealed elegant computations where the brain uses an internal model to overcome the dynamic limitations and ambiguities of the vestibular sensors (Mayne 1974; Oman 1982; Borah et al. 1988; Merfeld 1995; Zupan and Merfeld, 2002; Laurens 2006; Laurens and Droulez 2007, 2008; Laurens and Angelaki 2011; Karmali and Merfeld 2012; Lim et al. 2017). Neuronal correlates of these computations have been identified in brainstem and cerebellum (Angelaki et al. 2004; Shaikh et al. 2005; Yakusheva et al. 2007, 2008, 2013, Laurens et al. 2013a,b).

In the past decade, a few research groups have also studied how brainstem and cerebellar neurons modulate during active, self-generated head movements. Strikingly, several types of neurons, well-known for responding to vestibular stimuli during passive movement, lose or reduce their sensitivity during self-generated movement (Gdowski et al. 2000; Gdowski and McCrea 1999; Marlinski and McCrea 2009; McCrea et al. 1999; McCrea and Luan 2003; Roy and Cullen 2001; 2004; Brooks and Cullen 2009, 2013, 2014; Brooks et al. 2015; Carriot et al. 2014). In contrast, vestibular afferents respond indiscriminately for active and passive stimuli (Cullen and Minor 2002; Sadeghi et al. 2007; Jamali et al. 2009). These properties resemble sensory prediction errors in other sensorimotor functions such as fish electrosensation (Requarth and Sawtell 2011; Kennedy et al. 2014) and motor control (Tseng et al. 2007, Shadmer et al. 2010). Yet, a consistent quantitative take-home message has been lacking. Initial experiments implicated proprioceptive switches (Roy and Cullen, 2001; 2004). More recently, elegant experiments by Brooks, Cullen and colleagues (Brooks et al., 2015; Brooks and Cullen, 2015; Cullen and Brooks, 2015) have provided strong evidence that the brain predicts how self-generated motion activates the vestibular organs, and subtracts these predictions from afferent signals to generate sensory errors. The preferential response to passive motion, or to the passive components of combined active and passive motion, is then easily interpreted as sensory prediction error signals. However, the computational processes underlying this ‘prediction’ remain unclear.

Confronting the findings of studies utilizing passive and active motion stimuli leads to a paradox, where the brain appears to implement an elaborate internal model to process vestibular sensory signals, but suppresses the input of this internal model during self-generated motion. Thus, a highly influential interpretation is that the elaborate internal model characterized with passive stimuli would only be useful in situations that involve unexpected (passive) movements, such as recovering from loss of balance, but would remain unused during most normal activities. Here we propose an alternative - that the same internal model described previously for passive head movements also processes efference copies of motor commands during active head movements. Critically, we also demonstrate that accurate self-motion estimation would be severely compromised if signals were not processed by an identical internal model during actively-generated movements.

The essence of the theory developed previously for passive movements is that the brain uses an internal representation of the laws of physics and sensory dynamics (which can be elegantly modeled as forward internal models of the sensors) to process vestibular signals. In contrast, although it is understood that transforming head motor commands into sensory predictions is likely to also involve internal models, no explicit mathematical implementation has ever been proposed. Are the two internal models distinct? This is doubtful, because the same vestibular and cerebellar neurons appear involved in both (Angelaki and Cullen, 2008). If implemented by overlapping neural populations, how are the two internal models related to each other? A survey of the many studies by Cullen and colleagues even questions the origin of the sensory signals canceling vestibular afferent activity, as early studies emphasized a critical role of neck proprioception (Roy and Cullen, 2001; 2004). More recent studies generally implicate the generation of sensory prediction errors, without ever specifying whether the implicated forward internal models involve vestibular or proprioceptive cues (Brooks et al., 2015). We will show here that active and passive motion should be processed by the same forward internal models of all sensors involved in head motion sensation: canals, otolith and neck proprioceptors.

To demonstrate the unified framework governing self-motion sensation during both active and passive head movements, we use the framework of the Kalman filter (Fig. 1; Kalman 1960), which represents the simplest and most commonly used mathematical technique to implement statistically optimal dynamic estimation and explicitly computes sensory prediction errors. Here we build a quantitative Kalman filter model that integrates motor commands and vestibular signals during active and passive rotations, tilts and translations. We show how the same internal model may process both active and passive motion stimuli and provide quantitative simulations that reproduce a wide range of behavioral and neuronal responses. These simulations also generate testable predictions, in particular which passive stimuli should induce sensory errors and which should not, that may motivate future studies and guide interpretation of experimental findings. Finally, we summarize these internal model computations into a schematic diagram, and we discuss how various populations of brainstem and cerebellar neurons may encode the underlying sensory error or feedback signals.

**Figure 1:**
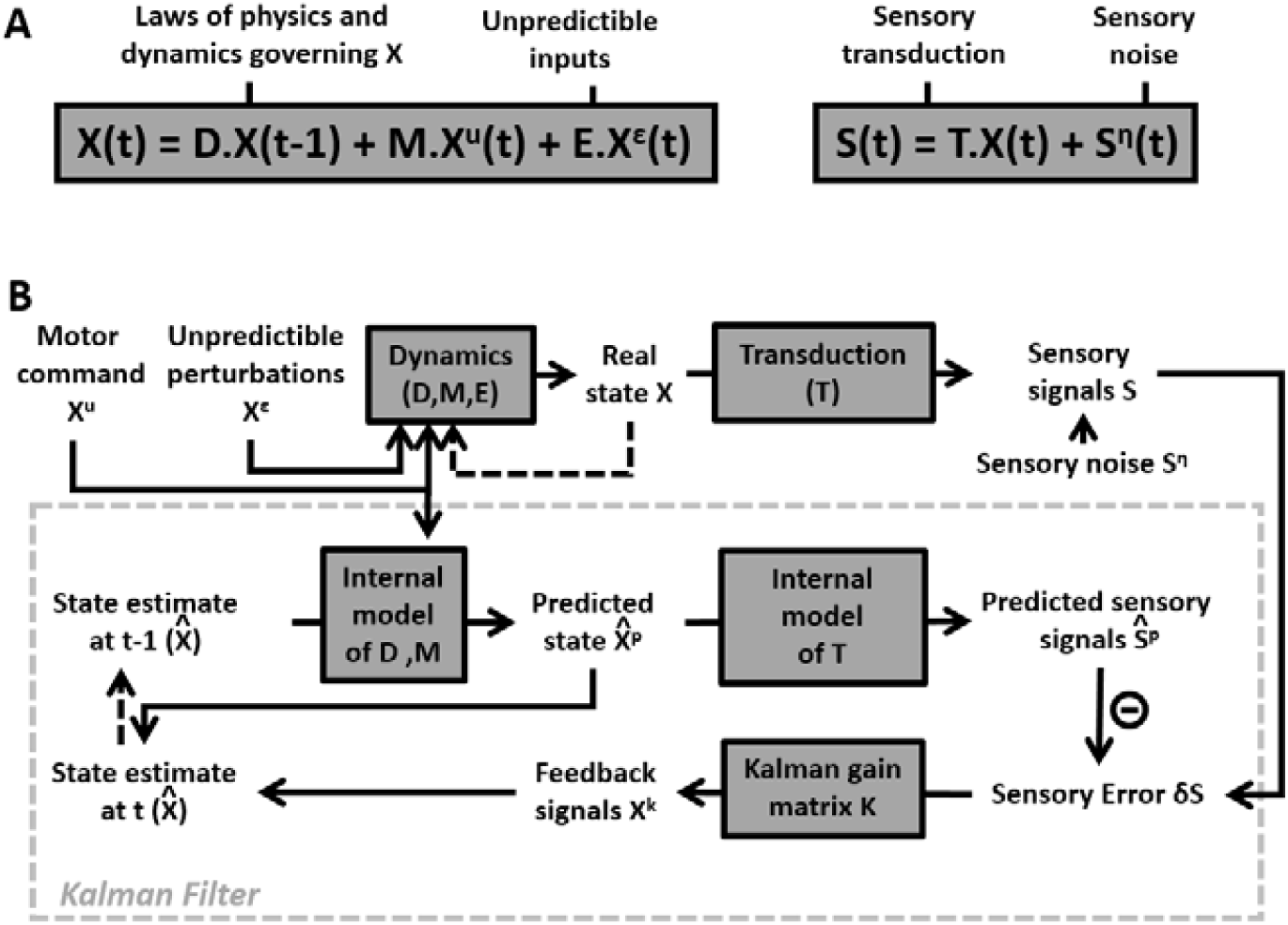
Generic structure of a Kalman Filter. (A) Equations of the Kalman filter algorithm describing how motor commands and sensory signals are processed for optimal state estimation. Motion variables (*X*) are computed as a function of motor commands *X*^*u*^ and unpredictable perturbations *X*^*ε*^. Matrices *D*, *M*, *E* and *T* encode the system’s dynamics. Sensory inputs *S* are computed as a function of *X* and sensory noise, *S*^*η*^. (B) Schematic implementing the equations of the Kalman filter algorithm. An internal estimate of motion variables (internal states, 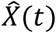) is computed dynamically as a function of 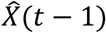, motor commands *X*^*u*^(*t*) and sensory signals *S*(*t*). The dashed arrows indicate that the estimate at time *t* is passed to the next time step, where it becomes the estimate at (*t* − 1). Sensory errors *δS* are transformed into feedback *X*^*k*^ = *K*. *δS*, where *K* is a matrix of feedback gains, whose rank is determined by the dimensionality of both the state variable *X* and the sensory signals *S*. The box defined by dashed gray lines illustrates the Kalman filter computations.

## Results

### Overview of Kalman filter model of head motion

A Kalman filter (Kalman 1960) is based on a forward model of a dynamical system, defined by a set of state variables *X* that are driven by their own dynamics, motor commands *X*^*u*^ and internal or external perturbations *X*^*ε*^. The evolution of *X* is modeled as *X*(*t*) = *D*. *X*(*t* − 1) + *M*. *X*^*u*^(*t*) + *E*. *X*^*ε*^ (Fig. 1A, left) where matrices *D*, *M* and *E* reflect the system’s dynamics and response to motor inputs and perturbations, respectively. A set of sensors, grouped in a variable *S*, measures state variables or combinations thereof (encoded in a matrix *T*) that can be modeled as *S*(*t*) = *T*. *X*(*t*) + *S*^*η*^(*t*), where *S*^*η*^ is sensory noise (Fig. 1A, right). Note that *S* may provide ambiguous or incomplete information, since some sensors may measure a mixture of state variables, and some variables may not be measured at all.

The Kalman filter, schematically represented in Fig. 1B, uses the available information to track an optimal internal estimate of the state variable 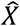. At each time *t*, the Kalman filter computes a preliminary estimate (also called a prediction) 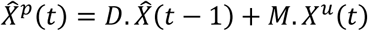 and a corresponding predicted sensory signal 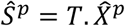. In general, the resulting state estimate 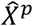 and the predicted sensory prediction 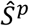 may differ from the real values *X* and *S* because: (1) *X*^*ε*^ ≠ 0, but the brain cannot predict the perturbation *X*^*ε*^(*t*), (2) the brain doesn’t know the value of the sensory noise *S*^*η*^(*t*) and (3) the previous estimate 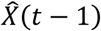 used to compute 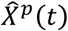 could be incorrect. These errors are reduced using sensory information, as follows (Fig. 1B): First, the prediction Ŝ^*p*^ and the sensory input *S* are compared to compute a sensory error *δS*. Second, sensory errors are transformed into a feedback *X*^*k*^ = *K*. *δS* where *K* is a matrix of feedback gains, whose dimensionality depends on both the state variable *X* and the sensory inputs. Thus, an improved estimate at time *t* is 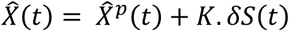. The feedback gain matrix *K* determines how sensory errors improve the final estimate, and is computed based on *D*, *E*, *T* and on the variances of *X*^*ε*^ and *S*^*η*^ (see Suppl. Methods, ‘Kalman filter algorithm’ for details).

Fig. 2 applies this framework to the problem of estimating self-motion (rotation, tilt, and translation) using vestibular sensors, with two types of motor commands: angular velocity (*Ω*^*u*^) and translational acceleration (*A*^*u*^), with corresponding unpredicted inputs, *Ω*^*ε*^ and *A*^*ε*^ (Fig. 2A) that represent passive motion or motor error (see Discussion: ‘Role of the vestibular system during active motion: fundamental, ecological and clinical implications’). The sensory signals (*S*) we consider initially encompass the semicircular canals (rotation sensors that generate a sensory signal *V*) and the otoliths organs (linear acceleration sensors that generate a sensory signal *F*) – proprioception is also added in subsequent sections. Each of these sensors has distinct properties, which can be accounted for by the internal model of the sensors (T in Fig. 1). The semicircular canals exhibit high-pass dynamic properties, which are modeled by another state variable *C*(*t*) (see Suppl. Methods, ‘Model of head motion and vestibular sensors’). The otolith sensors exhibit negligible dynamics, but are fundamentally ambiguous: they sense gravitational as well as linear acceleration – a fundamental ambiguity resulting from Einstein’s equivalence principle [Einstein 1907; modeled here as 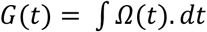 and *F*(*t*) = *G*(*t*) + *A*(*t*); note that *G* and *A* are expressed in comparable units; see Methods; “Simulation parameters”]. Thus, in total, the state variable *X* has 4-degrees of freedom (Fig. 2A): angular velocity *Ω* and linear acceleration *A* (which are the input/output variables directly controlled), as well as *C* (a hidden variable that must be included to model the dynamics of the semicircular canals) and tilt position *G* (another hidden variable that depends on rotations *Ω*, necessary to model the sensory ambiguity of the otolith organs).

**Figure 2:**
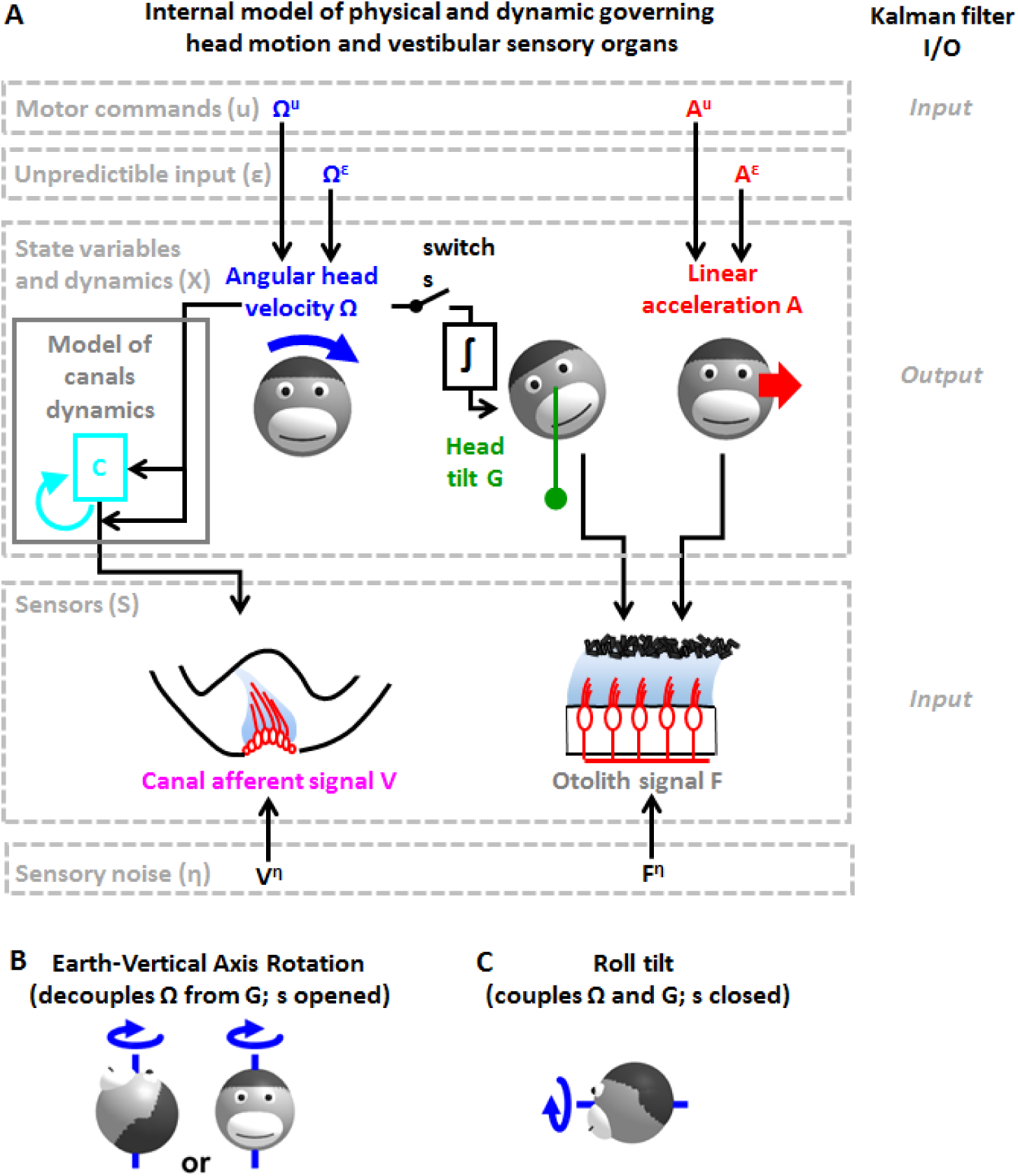
Application of the Kalman filter algorithm into optimal self-motion estimation using an internal model with four state variables and two vestibular sensors. (A) Schematic diagram of the model. Inputs to the model include motor commands, unexpected perturbations, as well as sensory signals. Motor commands during active movements, that is angular velocity (*Ω*^*u*^) and translational acceleration (*A*^*u*^), are known by the brain. Unpredicted internal or external factors such as external (passive) motion are modeled as variables *Ω*^*ε*^ and *A*^*ε*^. The state variable has 4 degrees of freedom: angular velocity *Ω*, tilt position *G*, linear acceleration *A* and a hidden variable *C* used to model the dynamics of the semicircular canals (see Methods). Two sensory signals are considered: semicircular canals (rotation sensors that generate a signal *V*) and the otoliths organs (linear acceleration sensors that generate a signal *F*). Sensory noise *V*^*η*^ and *F*^*η*^ is illustrated here, but omitted from all simulations for simplicity. (B, C) illustration of rotations around earth-vertical (B) and earth-horizontal (C) axes.

The Kalman filter computes optimal estimates 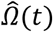, 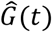, 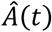 and 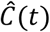 based on motor commands and sensory signals. Note that we don’t introduce any tilt motor command, as tilt is assumed to be controlled only indirectly though rotation commands (*Ω*^*u*^). For simplicity, we restrict self-motion to a single axis of rotation (e.g., roll) and a single axis of translation (inter-aural). The model can simulate either rotations in the absence of head tilt (e.g., rotations around an earth-vertical axis: EVAR, Fig. 2B) or tilt (Fig. 2C, where tilt is the integral of rotation velocity, 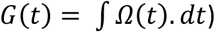 using a switch (but see Suppl. Methods, ‘Three-dimensional Kalman filter’ for a 3D model). Sensory errors are used to correct internal motion estimates using the Kalman gain matrix, such that the Kalman filter as a whole performs optimal estimation. In theory, the Kalman filter includes a total of 8 feedback signals, corresponding to the combination of 2 sensory (canal and otolith) errors and 4 internal states 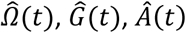 and 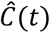]. From those 8 feedback signals, two are always negligible; see Suppl. Methods, ‘Kalman feedback gains’).

We will show how this model performs optimal estimation of self-motion using motor commands and vestibular sensory signals in a series of increasingly complex simulations. We start with a very short (200ms) EVAR stimulus, where canal dynamics is negligible (Fig. 3), followed by a longer EVAR that highlights the role of an internal model of the canals (Fig. 4). Next, we consider the more complex tilt and translation movements that require all 4 state variables to demonstrate how canal and otolith errors interact to disambiguate otolith signals (Fig. 5, 6). Finally, we extend our model to simulate independent movement of the head and trunk by incorporating neck proprioceptive sensory signals (Fig. 7). For each motion paradigm, identical active and passive motion simulations will be shown side by side in order to demonstrate how the internal model integrates sensory information and motor commands. We show that the Kalman feedback plays a preeminent role, which explains why lots of brain machinery is devoted to its implementation (see Discussion). For convenience, all mathematical notations are summarized in Table 1. For Kalman feedback gain nomenclature and numerical values, see Table 2.

**Figure 3:**
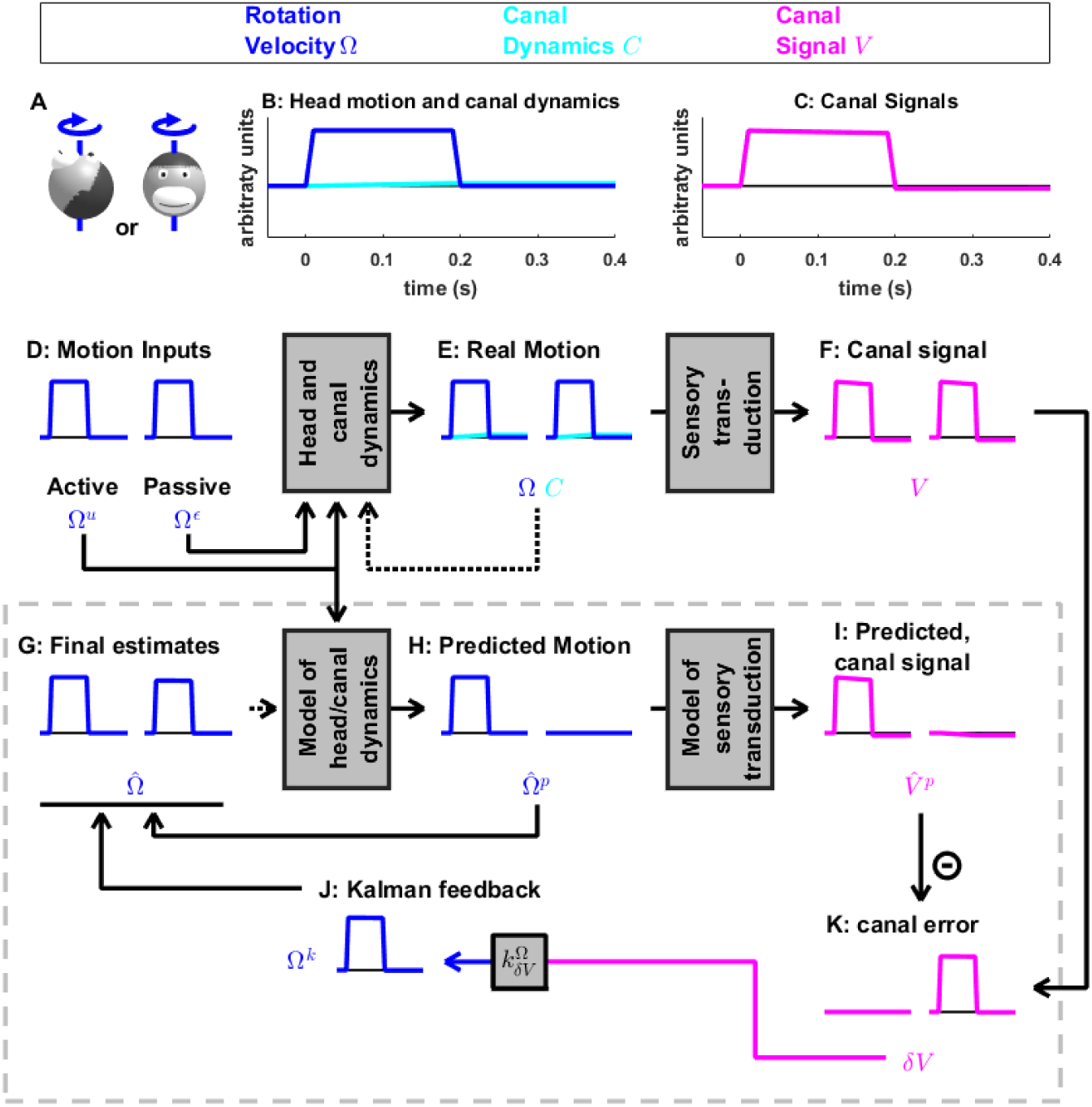
Short duration rotation around an earth-vertical axis (as in Fig. 2B). (A) Illustration of the stimulus lasting 200 ms. (B,C) Time course of motion variables and sensory (canal) signals. (D-K) Simulated variables during active (left panels) and passive motion (right panels). Only the angular velocity state variable *Ω* is shown (tilt position *G* and linear acceleration *A* are not considered in this simulation, and the hidden variable *C* is equal to zero). Continuous arrows represent the flow of information during one time step, and broken arrows the transfer of information from one time step to the next. (J) Kalman feedback. For clarity, the Kalman feedback is shown during passive motion only (it is always zero during active movements in the absence of any perturbation and noise). The box defined by dashed gray lines illustrates the Kalman filter computations. For the rest of mathematical notations, see Table 1.

**Figure 4:**
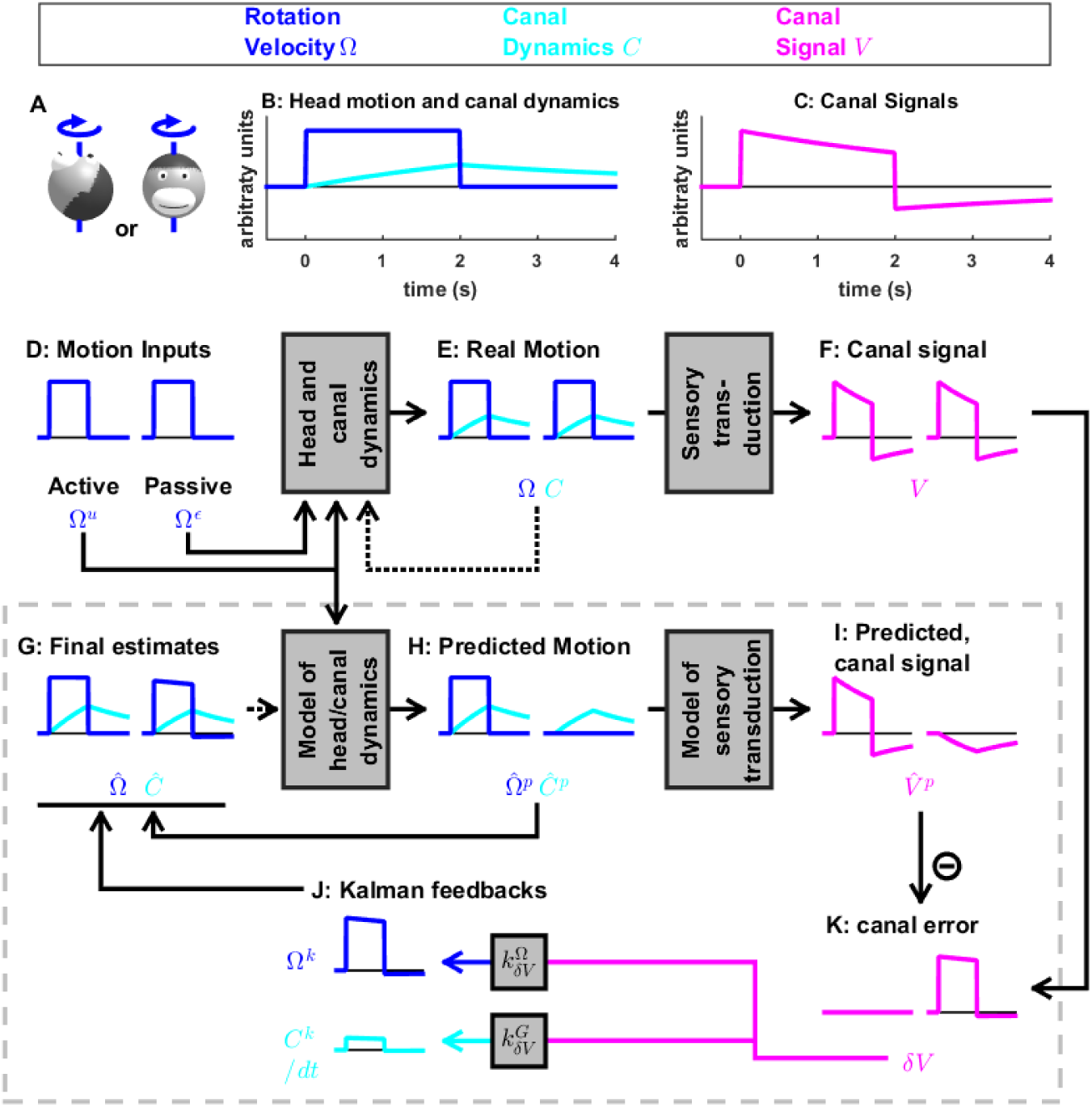
Medium-duration rotation around an earth-vertical axis, demonstrating the role of the internal model of canal dynamics. (A) Illustration of the stimulus lasting 2 s. (B,C) Time course of motion variables and sensory (canal) signals. (D-K) Simulated variables during active (left panels) and passive motion (right panels). Two state variables are shown: the angular velocity *Ω*(blue) and canal dynamics *C* (cyan). Continuous arrows represent the flow of information during one time step, and broken arrows the transfer of information from one time step to the next. (J) Kalman feedback. For clarity, the Kalman feedback (reflecting feedback from the canal error signal to the two state variables) is shown during passive motion only (it is always zero during active movements in the absence of any perturbation and noise). All simulations use a canal time constant of 4s. Note that, because of the integration, the illustrated feedback *C*^*k*^ is scaled by a factor 1/*δt*; see Suppl. Methods, ‘Kalman feedback gains’. The box defined by dashed gray lines illustrates the Kalman filter computations. For the rest of mathematical notations, see Table 1.

**Table 1:** List of motion variables and mathematical notations.

|  |  |  |
| --- | --- | --- |
| 1052 | *Motion variables* |  |
| 1053 |  |  |
| 1054 | $\Omega$ | Head rotation velocity (in space) |
| 1055 | $G$ | Head Tilt |
| 1056 | $A$ | Linear Acceleration |
| 1057 | $C$ | Canals dynamics |
| 1058 | $\Omega_{TS}$ | Trunk in space rotation velocity (variant of the model) |
| 1059 | $\Omega_{HT}$ | Head on trunk rotation velocity (variant of the model) |
| 1060 | $N$ | Neck position (variant of the model) |
| 1061 | $X$ | Matrix containing all motion variables in a model |
| 1062 |  |  |
| 1063 | *Sensory variables* |  |
| 1064 |  |  |
| 1065 | $V$ | Semicircular canal signal |
| 1066 | $F$ | Otolith signal |
| 1067 | $P$ | Neck proprioceptive signal |
| 1068 | $Vis$ | Visual rotation signal |
| 1069 | $S$ | Matrix containing all sensory variables in a model |
| 1070 |  |  |
| 1071 | *Accent and superscripts (motion variables)* |  |
| 1072 |  |  |
| 1073 | $X$ | Real value of a variable |
| 1074 | $\hat{X}$ | Final estimate |
| 1075 | $\hat{X}^p$ | Predicted (or preliminary) estimate |
| 1076 | $X^u$ | Motor command affecting the variable |
| 1077 | $X^\varepsilon$ | Perturbation or motor error affecting the variable (standard deviation $\sigma_X$) |
| 1078 | $\sigma_X$ | Standard deviation of $X^\varepsilon$ |
| 1079 | $X^k$ | Kalman feedback on the variable |
| 1080 |  |  |
| 1081 | *Accent and superscripts (sensory variables)* |  |
| 1082 |  |  |
| 1083 | $S$ | Real value of a variable |
| 1084 | $\hat{S}^p$ | Predicted value |
| 1085 | $S^\eta$ | Sensory noise |
| 1086 | $\sigma_S$ | Standard deviation of $S^\eta$ |
| 1087 | $\delta S$ | Sensory error |
| 1088 | $k_{\delta S}^X$ | Kalman gain of the feedback from $S$ to a motion variable $X$ |
| 1089 |  |  |
| 1090 | *Other* |  |
| 1091 |  |  |
| 1092 | $\delta t$ | Time step used in the simulations |
| 1093 | $M'$ | Transposed of a matrix $M$ |
| 1094 | $\tau_c$ | Time constant of the semicircular canals |

**Table 2:**
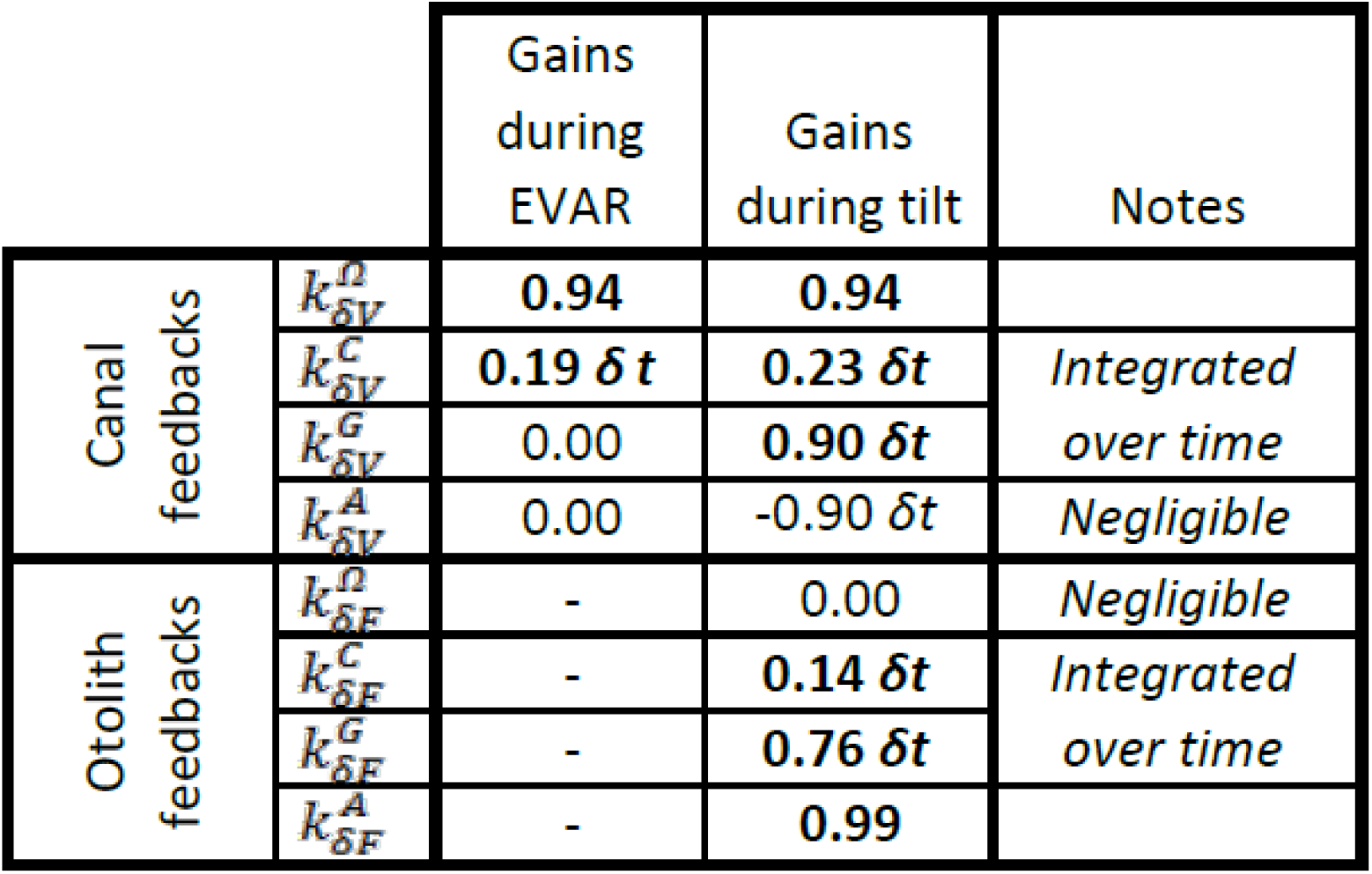
Kalman feedback gains during EVAR and tilt/translation. Some feedback gains are constant independently of *δt* while some other scale with *δt* (see Suppl. Methods, ‘Feedback gains’ for explanations). Gains that have negligible impact on the motion estimates are indicated in normal fonts, others with profound influence are indicated in bold. The feedback gains transform error signals into feedback signals.

### Passive motion induces sensory errors

Previous studies (McCrea and Luan 2003; Roy and Cullen 2001; 2004; Brooks and Cullen 2014) have proposed that vestibular and rostral fastigial nuclei neurons that respond specifically to passive rotations encode sensory errors resulting from the discrepancy between vestibular signals and the expected consequences of motor commands. Here, we demonstrate that the Kalman filter predicts sensory discrepancies consistent with these results.

In Fig. 3, we simulate rotations around an earth-vertical axis (Fig. 3A) with a short duration (0.2s, Fig. 3B), chosen to minimize canal dynamics (*C* ≈ 0, Fig. 3B, cyan) such that the canal response matches the velocity stimulus (*V* ≈ *Ω*, compare magenta curve in Fig. 3C with blue curve in Fig. 3B). We simulate active motion (Fig. 3D-K, left panels), where *Ω* = *Ω*^*u*^ (Fig. 3D) and *Ω*^*ε*^ = 0 (not shown), as well as passive motion (Fig. 3D-K, right panels), where *Ω* = *Ω*^*ε*^ (Fig. 3D) and *Ω*^*u*^ = 0 (not shown). The rotation velocity stimulus (*Ω*, Fig. 3E, blue) and canal activation (*V*, Fig. 3F, magenta) are identical in both active and passive stimulus conditions. As expected, the final velocity estimate 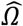 (output of the filter, Fig. 3G, blue) is equal to the stimulus *Ω* (Fig. 3E, blue) during both passive and active conditions. Thus, this first simulation is meant to emphasize differences in the flow of information *within* the Kalman filter, rather than differences in performance between passive and active motion (which is identical).

The fundamental difference between active and passive motion resides in the prediction of head motion (Fig. 3H) and sensory canal signals (Fig. 3I). During active motion, the motor command *Ω*^*u*^ (Fig. 3D) is converted into a predicted rotation 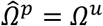 (Fig. 3H) by the internal model, and in turn in a predicted canal signal 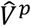 (Fig. 3I). Of course, in this case, we have purposely chosen the rotation stimulus to be so short (0.2s), such that canal afferents reliably encode the rotation stimulus (*V* ≈ *Ω*; compare Fig. 3F and 3E, left panels) and the internal model of canals dynamics has a negligible contribution; i.e., 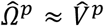 (compare Fig. 3I and 3H, left panels). Because the canal sensory error is null, i.e. 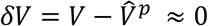 (Fig. 3K, left panel), the Kalman feedback pathway remains silent (not shown) and the net motion estimate is unchanged compared to the prediction, i.e. 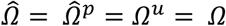. In conclusion, during active rotation (and in the absence of perturbations, motor or sensory noise), motion estimates are generated entirely based on an accurate predictive process, in turn leading to an accurate prediction of canal afferent signals. In the absence of sensory mismatch, these estimates don’t require any further adjustment.

In contrast, during passive motion the predicted rotation is null (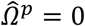, Fig. 3H, right panel), and therefore the predicted canal signal is also null (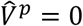, Fig. 3I, right panel). Therefore, canal signals during passive motion generate a sensory error 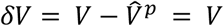(Fig. 3K, right panel). This sensory error is converted into a feedback signal 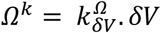 (Fig. 3J) with a Kalman gain 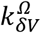 (feedback from canal error *δV* to angular velocity estimate Ω) that is close to 1 (Table 2; note that this value represents an optimum and is computed by the Kalman filter algorithm). The final motion estimate is generated by this feedback, i.e. 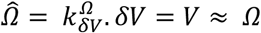.

These results illustrate the fundamental rules of how active and passive motion signals are processed by the Kalman filter (and, as hypothesized, the brain). During active movements, motion estimates are generated by a *predictive mechanism*, where motor commands are fed into an internal model of head motion. During passive movement, motion estimates are formed based on *feedback signals* that are themselves driven by sensory canal signals. In both cases, specific nodes in the network are silent (e.g., predicted canal signal during passive motion, Fig. 3I; canal error signal during active motion, Fig. 3K), but the same network operates in unison under *all* stimulus conditions. Thus, depending on whether the neuron recorded by a microelectrode in the brain carries *predicted*, *actual* or *error* sensory signals, differences in neural response modulation is expected between active and passive head motion. For example, if a cell encodes canal error exclusively, it will show maximal modulation during passive rotation, and no modulation at all during active head rotation. If a cell encodes mixtures of canal sensory error and actual canal sensory signals (e.g., through a direct canal afferent input), then there will be non-zero, but attenuated, modulation during active, compared to passive, head rotation. Indeed, a range of response attenuation has been reported in the vestibular nuclei (see Discussion).

We emphasize that in Fig. 3 we chose a *very* short-duration (0.2s) motion profile, where semicircular canal dynamics is negligible and the sensor can accurately follow the rotation velocity stimulus. We now consider more realistic rotation durations, and demonstrate how predictive and feedback mechanisms interact for accurate self-motion estimation. Specifically, canal afferent signals attenuate (because of their dynamics) during longer duration rotations – and this attenuation is already sizable for rotations lasting 1s or longer. We next demonstrate that the internal model of canal dynamics must be engaged for accurate rotation estimation, even during purely actively-generated head movements.

### Internal model of canals

We now simulate a longer head rotation, lasting 2s (Fig. 4A,B, blue). The difference between the actual head velocity *Ω* and the average canal signal *V* is modeled as an internal state variable *C*, which follows low-pass dynamics (see Suppl. Methods, ‘Model of head motion and vestibular sensors’). At the end of the 2s rotation, the value of *C* reaches its peak at ∼40% of the rotation velocity (Fig. 4B, cyan), modeled to match precisely the afferent canal signal *V*, which decreases by a corresponding amount (Fig. 4C). Note that *C* persists when the rotation stops, matching the canal aftereffect (*V* = −*C* < 0 after t>2s). Next we demonstrate how the Kalman filter uses the internal variable *C* to compensate for canal dynamics.

During active motion, the motor command *Ω*^*u*^ (Fig. 4D) is converted into an accurate prediction of head velocity 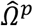(Fig. 4H, blue). Furthermore, *Ω*^*u*^ is also fed through the internal model of the canals to predict 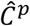 (Fig. 4H, cyan). By combining the predicted internal state variables 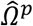 and 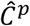, the Kalman filter computes a canal prediction 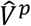 that follows the same dynamics as *V* (compare Fig. 4F and I, left panels). Therefore, as in Fig. 3, the resulting sensory mismatch is 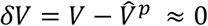 and the final estimates (Fig. 4G) are identical to the predicted estimates (Fig. 4H). Thus, the Kalman filter maintains an accurate rotation estimate by feeding motor commands though an internal model of the canal dynamics. Note, however, that because in this case *V*≠*Ω* (compare magenta curve in Fig. 4F and blue curve in Fig. 4E, left panels), 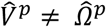 (compare magenta curve in Fig. 4I and blue curve in Fig. 4H, left panels). Thus, the sensory mismatch can only be null under the assumption that motor commands have been processed through the internal model of the canals. But before we elaborate on this conclusion, let’s first consider passive stimulus processing.

During passive motion, the motor command *Ω*^*u*^ is equal to zero. First, note that the final estimate 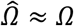 is accurate (Fig. 4G), as in Fig. 3G, although canal afferent signals don’t encode *Ω* accurately. Second, note that the internal estimate of canal dynamics 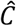 (Fig. 4G) and the corresponding prediction (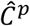; Fig. 4H) are both accurate (compare with Fig. 4E). This occurs because the canal error *δV* (Fig. 4K) is converted into a second feedback, *C*^*k*^, (Fig. 4J, cyan), which updates the internal estimate 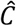 (see Suppl. Methods, ‘Velocity Storage’). Finally, in contrast to Fig. 3, the canal sensory error *δV* (Fig. 4K) does not follow the same dynamics as *V* (Fig. 4C,F), but is (as it should) equal to Ω (Fig. 4B). This happens because, though a series of steps (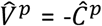 in Fig. 4I and 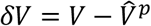 in Fig. 4K) 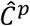, is added to the vestibular signal *V* to compute *δV* ≈ *D*. This leads to the final estimate 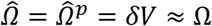 (Fig. 4G). Model simulations during even longer duration rotations and visual-vestibular interactions are illustrated in Fig. 4 Suppl. 1. Thus, the internal model of canal dynamics improves the rotation estimate during passive motion. Remarkably, this is important not only during very long duration rotations (as is often erroneously presumed), but also during short stimuli lasting 1-2s, as illustrated with the simulations in Fig. 4.

We now return to the actively-generated head rotations to ask the important question: What would happen if the brain didn’t use an internal model of canal dynamics? We simulated motion estimation where canal dynamics was removed from the internal model used by the Kalman filter (Fig. 4 Suppl 2). During both active and passive motion, the net estimate 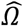 is inaccurate as it parallels *V*, exhibiting a decrease over time and an aftereffect. In particular, during active motion, the motor commands provide accurate signals 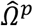, but the internal model of the canals fails to convert them into a correct prediction 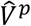, resulting in a sensory mismatch. This mismatch is converted into a feedback signal *Ω*^*k*^ that degrades the accurate prediction 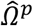 such that the final estimate 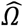 is inaccurate. These simulations highlight the role of the internal model of canal dynamics, which continuously integrates rotation information in order to anticipate canal afferent activity during *both* active and passive movements. Without this sensory internal model, active movements would result in sensory mismatch, and the brain could either transform this mismatch into sensory feedback, resulting in inaccurate motion estimates. Note that the need for the internal model is important even during natural short-duration and high-velocity head rotations (Fig. 4 Suppl. 3). Thus, even though particular nodes (neurons) in the circuit (e.g. vestibular and rostral fastigial nuclei cells presumably reflecting either *δV* or *Ω*^*k*^ in Fig. 3, 4; see Discussion) are attenuated or silent during active head rotations, efference copies of motor commands must *always* be processed though the internal model of the canals – motor commands *cannot* directly drive appropriate sensory prediction errors. This intuition has remained largely unappreciated by studies comparing how central neurons modulate during active and passive rotations – a misunderstanding that has led to the seemingly strawman dichotomy belittling all important insights gained by decades of studies using passive motion stimuli (see Discussion).

### Active versus passive tilt

Next, we study the interactions between rotation, tilt and translation perception. We first simulate a short duration (0.2s) roll tilt (Fig. 5A; with a positive tilt velocity *Ω*, Fig. 5B, blue). Tilt position (*G*, Fig. 5B, green) ramps during the rotation and then remains constant. As in Fig. 3, canal dynamics *C* is negligible (*V* ≈ *Ω*; Fig. 5F, magenta) and the final rotation estimate 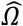 is accurate (Fig. 5G, blue). Also similar to Fig. 3, 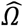 is carried by the predicted head velocity node during active motion 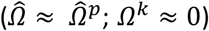 and by the Kalman feedback node during passive motion (*Ω* ≈ *Ω*^*k*^; 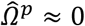). That is, the final rotation estimate, which is accurate during both active and passive movements, is carried by different nodes (thus, likely different cell types; see Discussion) within the neural network.

**Figure 5:**
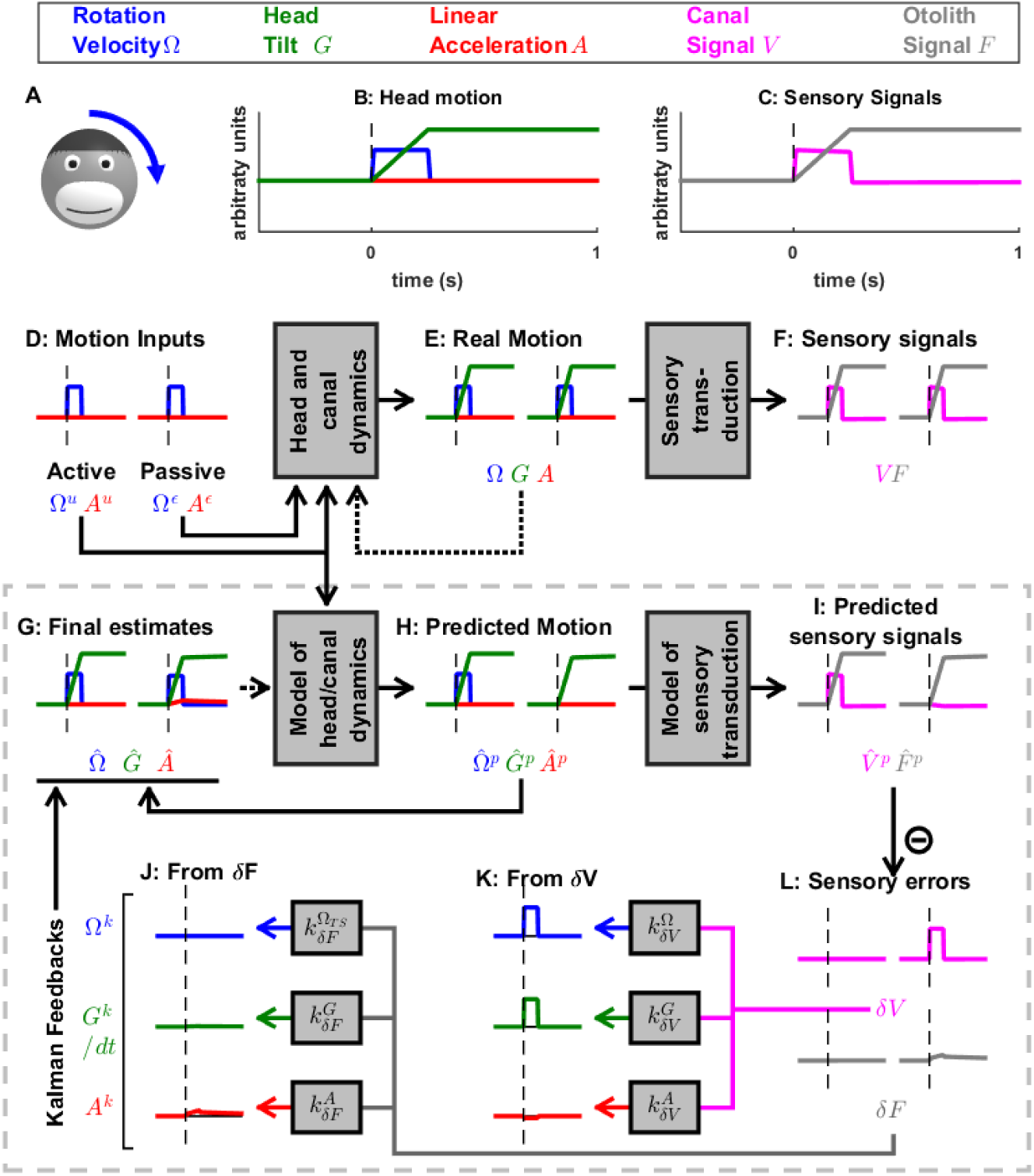
Simulation of short duration head tilt. (A) Illustration of the stimulus lasting 0.2 s. (B,C) Time course of motion variables and sensory (canal and otolith) signals. (D-L) Simulated variables during active (left panels) and passive motion (right panels). Three state variables are shown: the angular velocity *Ω* (blue), tilt position *G*, and linear acceleration *A*. Continuous arrows represent the flow of information during one time step, and broken arrows the transfer of information from one time step to the next. (J, K) Kalman feedback (shown during passive motion only). Two error signals (*δV*: canal error; *δF*: otolith error) are transformed into feedbacks to state variables *Ω*^*k*^: blue, *G*^*k*^: green, *A*^*k*^: red (variable *C*^*k*^ is not shown, but see Fig. 5, Suppl. 1 for simulations of a 2s tilt). Feedback originating from *δF* is shown in (J) and from *δV* in (K). The feedbacks to *G*^*k*^ are scaled by a factor 1/*δt* (see Suppl. Methods, ‘Kalman feedback gains’). Note that in this simulation we consider an active (*Ω*^*u*^) or passive (*Ω*^*ε*^) rotation velocity as input. The tilt itself is a consequence of the rotation, and not an independent input. The box defined by dashed gray lines illustrates the Kalman filter computations. For the rest of mathematical notations, see Table 1.

When rotations change orientation relative to gravity, another internal state (tilt position *G*, not included in the simulations of Fig. 3 and 4) and another sensor (otolith organs; *F* = *G* since *A* = 0; Fig. 5F, black) are engaged. During actively-generated tilt movements, the rotation motor command (*Ω*^*u*^) is temporally integrated by the internal model of head motion (see *Eq*. 3*c* of Suppl. Methods, ‘Kalman filter algorithm developed’), generating an accurate prediction of head tilt 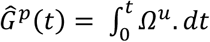 (Fig. 5H, left panel, green). This results in a correct prediction of the otolith signal 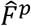 (Fig. 5I, grey) and therefore, as in previous simulations of active movement, the sensory mismatch for both the canal and otolith signals (Fig. 5L, magenta and gray, respectively) and feedback signals (not shown) are null; and the final estimates, driven exclusively by the prediction, are accurate; 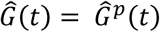 and 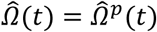.

During passive tilt, the canal error, *δV*, is converted into Kalman feedback that updates 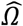 (Fig. 5K, blue) and 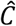 (not shown here; but see Fig. 5 Suppl. 1 for 2s tilt simulations), as well as the two other state variables (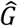 and 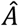). Specifically, the feedback from *δV* to 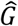 updates the predicted tilt 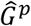 and is temporally integrated by the Kalman filter (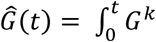 see Suppl. Methods, ‘Passive Tilt’; Fig. 5K, green). The feedback signal from *δV* to 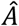 has a minimal impact, as illustrated in Fig. 5K, red (see also Suppl. Methods, ‘Kalman feedback gains’ and Table 2).

Because *δV* efficiently updates the tilt estimate 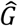 the otolith error *δF* is close to zero during passive tilt (Fig. 5L, gray; see Suppl. Methods, ‘Passive Tilt’) and therefore all feedback signals originating from *δF* (Fig. 5J) play a minimal role (see Suppl. Methods, ‘Passive Tilt’) during pure tilt (this is the case even for longer duration stimuli; Fig. 5 Suppl. 1). This simulation highlights that, although tilt is sensed by the otoliths, passive tilt doesn’t induce any sizeable otolith error. Thus, unlike neurons tuned to canal error, the model predicts that those cells tuned to otolith error will not modulate during either passive or actively-generated head tilt. Therefore, cells tuned to otolith error would respond primarily during translation, and not during tilt, thus they would be identified ‘translation-selective’. Furthermore, the model predicts that those neurons tuned to passive tilt (e.g., Purkinje cells in the caudal cerebellar vermis; Laurens et al., 2013b) likely reflect canal error (Fig. 5L, magenta). Thus, the model predicts that tilt-selective Purkinje cells should encode tilt velocity, and not tilt position, a prediction that remains to be tested experimentally (see Discussion).

### Otolith errors are interpreted as translation and tilt with distinct dynamics

Next, we simulate a brief translation (Fig. 6). During active translation, we observe, as in previous simulations of active movements, that the predicted head motion matches the sensory (otolith in this case: *F* = *A*) signals (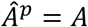 and 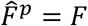). Therefore, as in previous simulations of active motion, the sensory prediction error is zero (Fig. 6L) and the final estimate is equal to, and driven by, the prediction (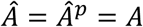 Fig. 6G, red).

**Figure 6:**
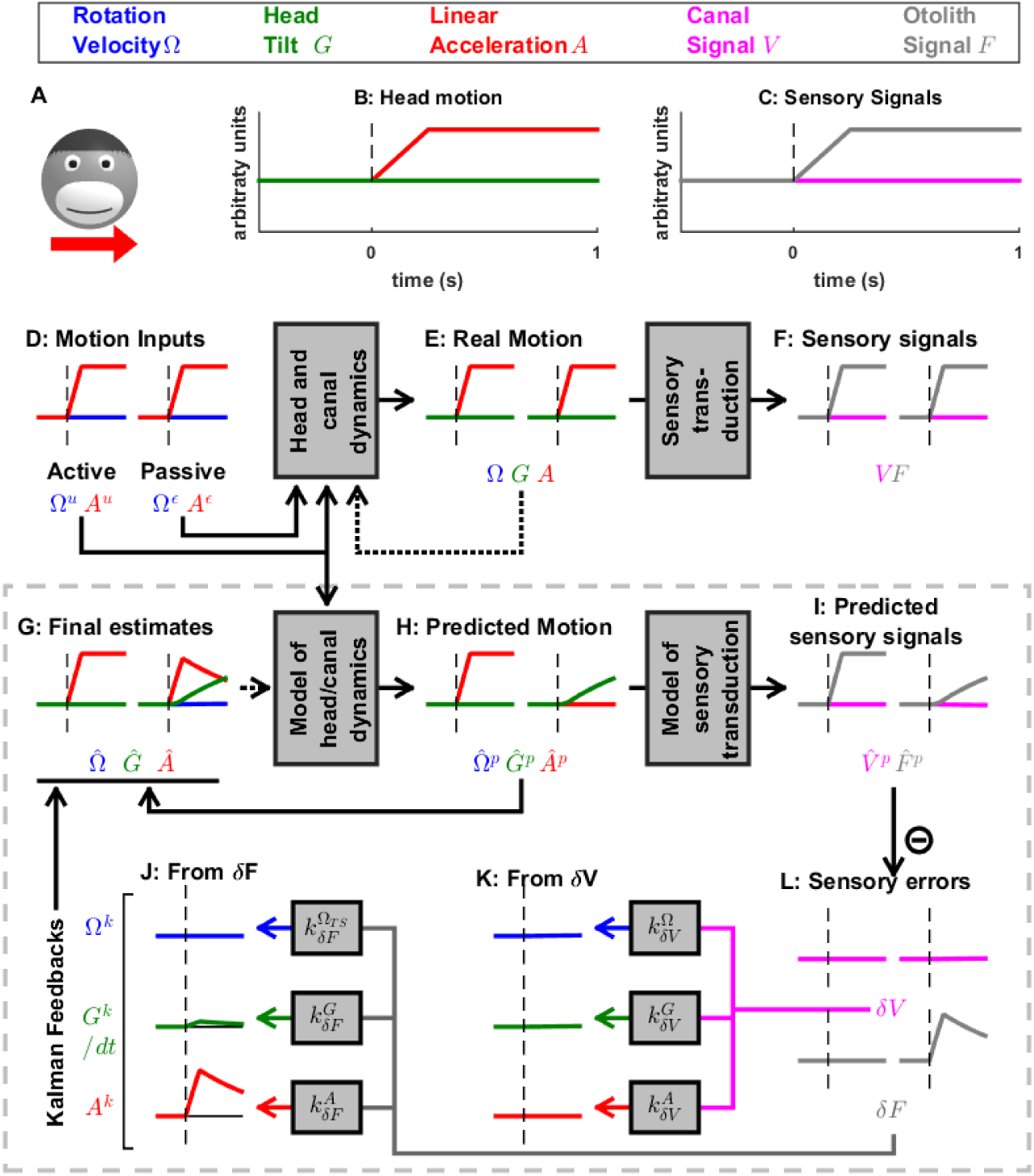
Simulation of short duration translation. Same legend as Fig. 5. Note that *F* is identical in Fig. 5 and 6: in terms of sensory inputs, these simulation differ only in the canal signal.

During passive translation, the predicted acceleration is null (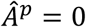, Fig 6H, red), similar as during passive rotation in Fig. 3, 4). However, a sizeable tilt signal (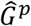 and 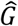, Fig. 6G,H, green), develops over time. This (erroneous) tilt estimate can be explained as follows: soon after translation onset (vertical dashed lines in Fig. 6B-J), 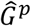 is close to zero. The corresponding predicted otolith signal is also close to zero 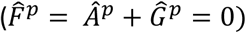, leading to an otolith error *δF* ≈ *A* (Fig. 6L, right, gray). Through the Kalman feedback gain matrix, this otolith error, *δF*, is converted into: (1) an acceleration feedback *A*^*k*^ (Fig. 6J, red) with gain 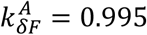 (the close to unity feedback gain indicates that otolith errors are interpreted as acceleration: 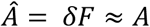; note however that the otolith error *δF* vanishes over time, as explained next); and (2) a tilt feedback *G*^*k*^ (Fig. 6J, green), with 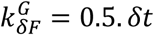. This tilt feedback, although too weak to have any immediate effect, is integrated over time (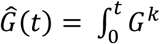; see Fig. 5 and Suppl. Methods, ‘Somatogravic effect’), generating the rising tilt estimate 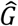 (Fig. 6G, green) and 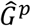 (Fig. 6H, green).

The fact that the Kalman gain feedback from the otolith error to the 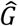 internal state generates the somatogravic effect is illustrated in Fig. 6 Suppl. 1, where a longer acceleration (20s) is simulated. At the level of final estimates (perception), these simulations predict the occurrence of tilt illusions during sustained translation (somatogravic illusion; Graybiel 1952; Paige and Seidman 1999). Further simulations show how activation of the semicircular canals without a corresponding activation of the otoliths (e.g., during combination of tilt and translation; Angelaki et al., 2004; Yakusheva et al., 2007) leads to an otolith error (Fig. 6 Suppl. 2) and how signals from the otoliths (that sense indirectly whether or not the head rotates relative to gravity) can also influence the rotation estimate 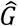 at low frequencies (Fig. 6 Suppl. 3; this property has been extensively evaluated by Laurens and Angelaki, 2011). These simulations demonstrate that the Kalman filter model efficiently simulates all previous properties of both perception and neural responses during passive tilt and translation stimuli (see Discussion).

The model analyzed so far has considered only vestibular sensors. Nevertheless, active head rotations often also activate neck proprioceptors, when there is an independent rotation of the head relative to the trunk. Indeed, a number of studies (Brooks and Cullen 2009; 2013; Brooks et al. 2015) have identified neurons in the rostral fastigial nuclei that encode the rotation velocity of the trunk. These neurons receive convergent signals from the semicircular canals and neck muscle proprioception and, accordingly, are named ‘bimodal neurons’, to contrast with ‘unimodal neurons’, which encode passive head velocity. Because the bimodal neurons don’t respond to active head and trunk movements, they likely encode feedback signals related to trunk velocity. We developed a variant of the Kalman filter to model both unimodal and bimodal neuron types (Fig. 7; see also Suppl. Methods and Fig. 7 Suppl. 1-3).

**Figure 7:**
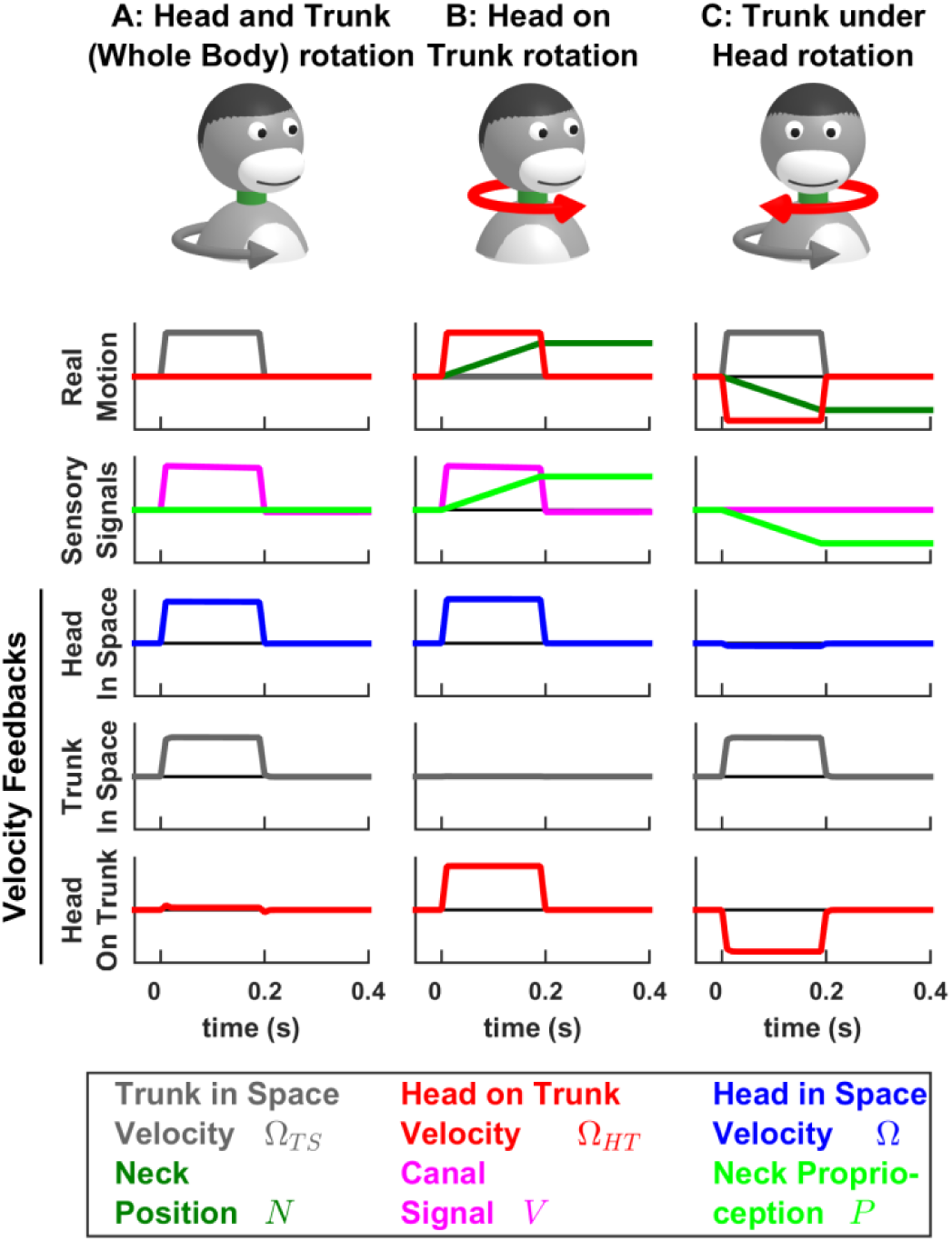
Simulations of passive trunk and head movements. We use a variant of the Kalman filter model (see Suppl. Methods) that tracks the velocity of both head and trunk (trunk in space: gray; head in space: blue; head on trunk: red) based on semicircular canal and neck proprioception signals. The real motion (first line), sensory signals (second line) and velocity feedback signals (third to fifth lines) are shown during (A) passive whole head and trunk rotation, (B) passive head on trunk rotation, and (C) passive trunk under head rotation. See Fig. 7 Suppl. 1-3 for other variables and simulations of active motion.

### Neck proprioceptors and encoding of trunk versus head velocity

The model tracks the velocity of the trunk in space *Ω*_*TS*_ and the velocity of the head on the trunk *Ω*_*HT*_ as well as neck position 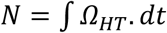. Sensory inputs are provided by the canals (that sense the total head velocity, *Ω* = *Ω*_*TS*_ + *Ω*_*HT*_), and proprioceptive signals from the neck musculature, which are assumed to encode neck position (*P*).

In line with previous simulations, we find that, during active motion, the predicted sensory signals are accurate. Consequently, the Kalman feedback pathways are silent (Fig. 7 Suppl. 1-3; active motion is not shown in Fig. 7). In contrast, passive motion induces sensory errors and Kalman feedbacks. The velocity feedback signals (elaborated in Fig. 7 Suppl. 1-3) have been re-plotted in Fig. 7, where we illustrate head in space (blue), trunk in space (gray), and head on trunk (red) velocity (neck position feedback signals are only shown in Fig. 7 Suppl. 1-3).

During passive whole head and trunk rotation, where the trunk rotates in space (Fig. 7A, Real motion: *Ω*_*TS*_ > 0, grey) and the head moves together with the trunk (head on trunk velocity *Ω*_*HT*_ = 0, red, head in space *Ω* > 0, blue), we find that the resulting feedbacks accurately encode these rotation components (Fig. 7A, Velocity Feedbacks; see also Fig. 7 Suppl. 1). During head on trunk rotation (Fig. 7B, Fig. 7 Suppl. 2), the Kalman feedbacks accurately encode the head on trunk (red) or in space (blue) rotation, and the absence of trunk in space rotation (gray). Finally, during trunk under head rotation that simulates a rotation of the trunk while the head remains fixed in space, resulting in a neck counter-rotation, the various motion components are accurately encoded by Kalman feedbacks (Fig. 7C, Fig. 7 Suppl. 3). We propose that unimodal and bimodal neurons reported in (Brooks and Cullen 2009, 2013) encode feedback signals about the velocity of the head in space (*Ω*^*k*^, Fig. 7, blue) and of the trunk in space (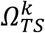, Fig. 7, gray), respectively. Furthermore, in line with experimental findings (Brooks and Cullen 2013), these feedback pathways are silent during self-generated motion.

The Kalman filter makes further predictions that are entirely consistent with experimental results. First, it predicts that proprioceptive error signals during passive neck rotation encode velocity (Fig. 7 Suppl. 3L; see Suppl. Methods, ‘Feedback signals during neck movement’). Thus, the Kalman filter explains the striking result that the proprioceptive responses of bimodal neurons encode trunk *velocity* (Brooks and Cullen 2009; 2013), even if neck proprioceptors encode neck position. Note that neck proprioceptors likely encode a mixture of neck position and velocity at high frequencies (Mergner et al. 1991). Additional simulations (not shown) where neck proprioceptive signals are assumed to encode mixtures of position and velocity yield similar results as those shown here. We used a model where neck proprioceptors encode position for simplicity, and in order to demonstrate that Kalman feedbacks encode trunk velocity even when proprioceptive signals encode position.

Second, the model predicts another important property of bimodal neurons: their response gains to both vestibular (during sinusoidal motion of the head and trunk together) and proprioceptive (during sinusoidal motion of the trunk when the head is stationary) stimulation vary identically if a constant rotation of the head relative to the trunk is added, as an offset, to the sinusoidal motion (Brooks and Cullen 2009). We propose that this offset head rotation extends or contracts individual neck muscles and affects the signal to noise ratio of neck proprioceptors. Indeed, simulations shown in Fig. 7 Suppl. 4 reproduce the effect of head rotation offset on bimodal neurons. In agreement with experimental findings, we also find that simulated unimodal neurons are not affected by these offsets (Fig. 7 Suppl. 4).

Finally, the model also predicts the dynamics of trunk and head rotation perception during long-duration rotations (Fig. 7 Suppl. 5), which has been established by behavioral studies (Mergner et al. 1991).

### Interactions between active and passive motion

The theoretical framework of the Kalman filter asserts that the brain cancels predictable sensory inputs, leading to an attenuation of central vestibular responses, which presumably encode sensory errors or feedback gains, during active motion. An alternative explanation for the attenuation of central responses could be that the brain simply suppresses vestibular sensory inflow. Experimental evidence in favor of the Kalman filter framework comes from recordings performed when passive motion is applied concomitantly to an active movement (Brooks and Cullen 2013, 2014; Carriot et al. 2013). Indeed, neurons that respond during passive but not active motion have been found to encode the passive component of combined passive and active motion, as expected based on the Kalman framework. We present corresponding simulation results in Fig. 8.

**Figure 8:**
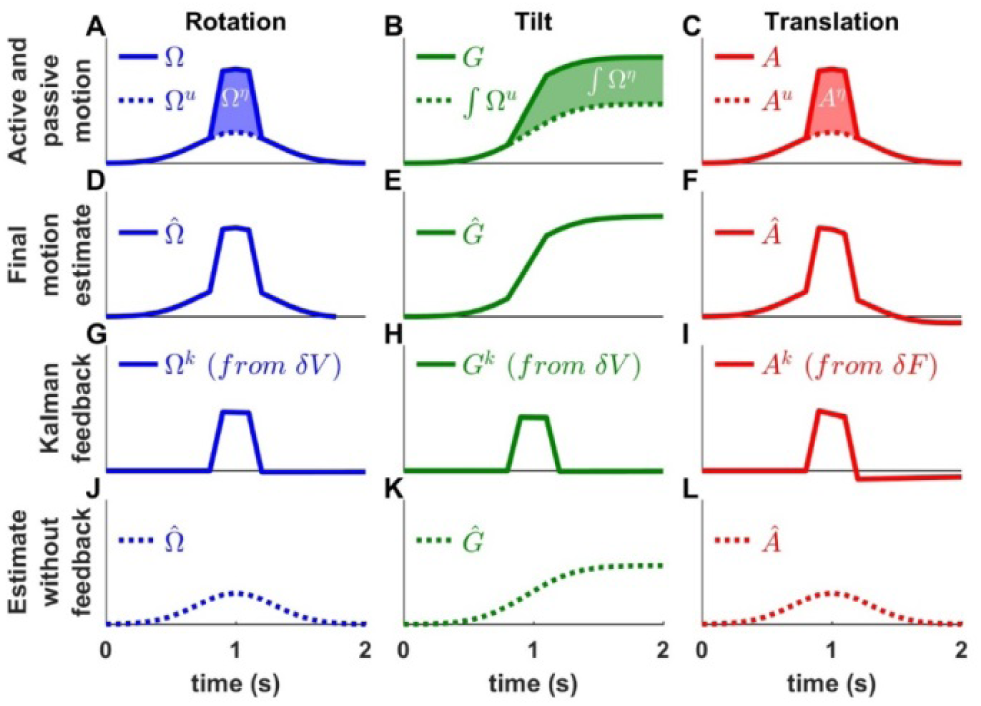
Interaction of active and passive motion. Active movements (Gaussian profiles) and passive movements (short trapezoidal profiles) are superimposed. (A) Active (*Ω*^*u*^) and passive (*Ω*^*ɛ*^) rotations. (B) Head tilt resulting from active and passive rotations (the corresponding tilt components are *∫ Ω*^*u*^. *dt* and *∫ Ω*^*ɛ*^. *dt*). (C) Active (*A*^*u*^) and passive (*A*^*ɛ*^) translations. (D-F) Final motion estimates (equal to the total motion). (G-I) The Kalman feedbacks correspond to the passive motion component. (J-K) Final estimates computed by inactivating all Kalman feedback pathways. These simulations represent the motion estimates that would be produced if the brain suppressed sensory inflow during active motion.

We simulate a rotation movement (Fig. 8A), where an active rotation (*Ω*^*u*^, Gaussian velocity profile) is combined to a passive rotation (*Ω*^*£*^, trapezoidal profile), a tilt movement (Fig. 8B; using similar velocity inputs, *Ω*^*u*^ and *Ω*^*£*^, where the resulting active and passive tilt components are 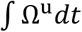 and 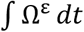), and a translation movement (Fig. 8C). We find that, in all simulations, the final motion estimate (Fig. 8D-F; 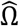, 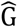 and 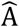, respectively) matches the combined active and passive motion (*Ω*, G and A, respectively). In contrast, the Kalman feedbacks (Fig. 8G-I) specifically encode the passive motion components. Specifically, the rotation feedback (*Ω*^*k*^, Fig. 8G) is identical to the passive rotation *Ω*^*ε*^ (Fig. 8A). As in Fig. 5, the tilt feedback (*G*^*k*^, Fig. 8H) encodes tilt velocity, also equal to *Ω*^*ε*^ (Fig. 8A). Finally, the linear acceleration feedback (*A*^*k*^, Fig. 8I) follows the passive acceleration component, although it decreases slightly with time because of the somatogravic effect. Thus, Kalman filter simulations confirm that neurons that encode sensory mismatch or Kalman feedback should selectively follow the passive component of combined passive and active motion.

What would happen if the brain simply discarded vestibular sensory (or feedback) signals during active motion? We repeat these simulations after removing the vestibular sensory input signals from the Kalman filter. We find that the net motion estimates encode only the active movement components (Fig. 8J-L; 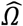, 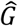 and 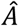) – thus, not accurately estimating the true movement. Furthermore, as a result of the sensory signals being discarded, all sensory errors and Kalman feedbacks are null. These simulations indicate that suppressing vestibular signals during active motion would prevent the brain from detecting passive motion occurring during active movement (see Discussion, “Role of the vestibular system during active motion: ecological, clinical and fundamental implications.”), in contradiction with experimental results.

## Discussion

We have extended a well-established model where the brain processes vestibular information optimally using internal models (Laurens 2006; Laurens and Droulez, 2007; 2008; Laurens et al. 2010, 2011a; Laurens and Angelaki, 2011; Laurens et al. 2013a,b) by incorporating efference copies of motor commands during active motion. Previous studies (Gdowski et al. 2000; Gdowski and McCrea 1999; Marlinski and McCrea 2009; McCrea et al. 1999; McCrea and Luan 2003; Roy and Cullen 2001; 2004; Brooks and Cullen 2009, 2013, 2014; Brooks et al. 2015; Carriot et al. 2014) have suggested that the brain cancels vestibular reafference during active motion, but an actual quantitative model has never been presented. We have tested the hypothesis that this postulated cancellation mechanism uses exactly the same sensory internal model computations already discovered using passive motion stimuli (Mayne 1974; Oman 1982; Borah et al. 1988; Merfeld 1995; Zupan and Merfeld, 2002; Laurens 2006; Laurens and Droulez 2007, 2008; Laurens and Angelaki 2011; Karmali and Merfeld 2012; Lim et al. 2017). Presented simulations confirm the hypothesis that the *same* internal model (consisting of forward internal models of the canals, otoliths and neck proprioceptors) can reproduce behavioral and neuronal responses to *both* active and passive motion. The formalism of the Kalman filter allows predictions of internal variables during both active and passive motion, with a strong focus on sensory error and feedback signals, which we hypothesize are realized in the response patterns of central vestibular neurons.

Perhaps most importantly, this work resolves an apparent paradox between active and passive movements (Angelaki and Cullen, 2008), by placing them into a unified theoretical framework where a single internal model tracks head motion based on motor commands and sensory feedback signals. We have shown here that internal model computations, which have been extensively studied through decades of passive motion experiments, are equally required to process active motion signals. These computational elements operate during both passive and active head movements, although particular cell types that encode sensory errors or feedback signals may not modulate during active movements because the corresponding sensory prediction error is negligible. This highlights the relevance and importance of passive motion stimuli, as critical experimental paradigms that can efficiently interrogate the network for establishing computational principles shared between active and passive movements, but which cannot easily be disentangled during active movements.

### Summary of the Kalman filter model

We have developed the first ever model that simulates self-motion estimates during both actively-generated and passive head movements. This model, summarized schematically in Fig. 9, transforms motor commands and Kalman filter feedback signals into internal estimates of head motion (rotation and translation) and predicted sensory signals. There are two important take-home messages: (1) Because of the physical properties of the two vestibular sense organs, the predicted motion generated from motor commands is not equal to predicted sensory signals (for example, the predicted rotation velocity is processed to account for canal dynamics in Fig. 4). Instead, the predicted rotation, tilt and translation signals generated by efference copies of motor commands must be processed by the corresponding forward models of the sensors in order to generate accurate sensory predictions. This important insight about the nature of these internal model computations has not been appreciated by the qualitative schematic diagrams of previous studies. (2) In an environment devoid of externally generated passive motion, motor errors and sensory noise, the resulting sensory predictions would always match sensory afferent signals accurately. In a realistic environment, however, unexpected head motion occurs due to both motor errors and external perturbations (see ‘Role of the vestibular system during active motion: ecological, clinical and fundamental implications’). Sensory vestibular signals are then used to correct internal motion estimates through the computation of sensory errors and their transformation into Kalman feedback signals. Given two sensory errors (*δV* originating from the semicircular canals and *δF* originating from the otoliths) and four internal state variables (rotation, internal canal dynamics, tilt and linear acceleration: 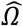, 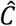, 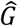, 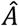), 8 feedback signals must be constructed. However, in practice, two of these signals have negligible influence for all movements (δV feedback to 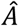 and *δF* feedback to 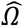; see Table 2 and Suppl. Methods, ‘Kalman Feedback Gains’), thus only 6 elements are summarized in Fig. 9.

**Figure 9:**
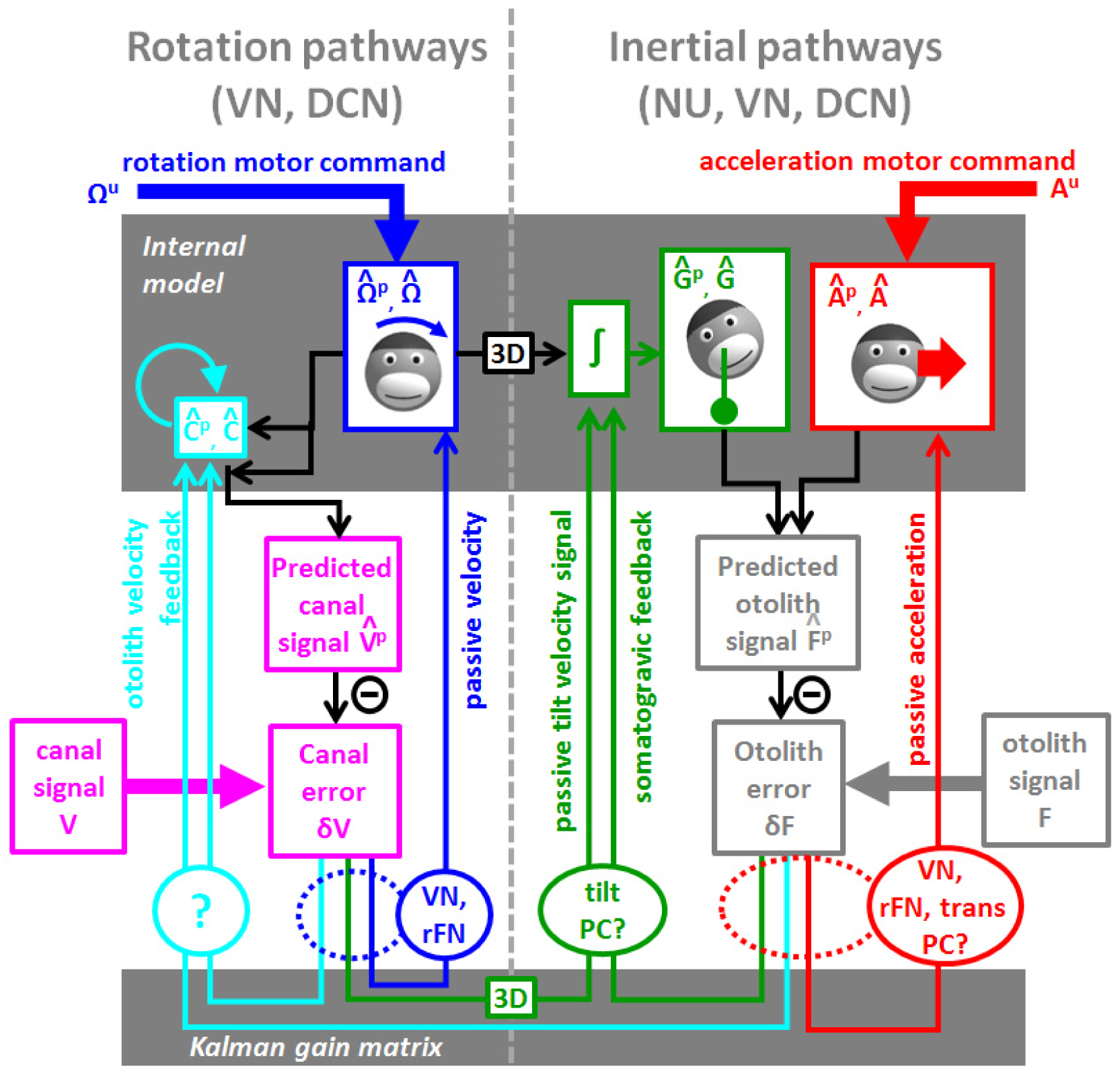
Schematic diagram of central vestibular computations. This diagram is organized to offer a synthetic view of the processing elements, as well as their putative neural correlates. An internal model (top gray box) predicts head motion based on motor commands and receives feedback signals. The internal model computes predicted canal and otolith signals that are compared to actual canal and otolith inputs. The resulting sensory errors are transformed by the Kalman gain matrix into a series of feedback ‘error’ signals. Left: canal error feedback signals; Right: otolith error feedback signals. Rotation signals are spatially transformed (“3D” boxes) into tilt velocity signals. Ovals indicate putative neuronal correlates of the feedback signals (VN: vestibular only vestibular nuclei neurons; rFN: rostral fastigial nuclei neurons, PC: Purkinje cells in the caudal vermis, DCN: deep cerebellar nuclei).

The non-negligible feedback signals originating from the canal error *δV* are as follows (Fig. 9, left):

- The feedback to the rotation estimate 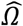 represents the traditional “direct” vestibular pathway (Raphan et al. 1979, Laurens and Angelaki 2011). It is responsible for rotation perception during high-frequency (unexpected) vestibular stimulation, and has a gain close to unity.
- The feedback to 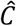 feeds into the internal model of the canals, thus allowing compensation for canals dynamics. This pathway corresponds to the “velocity storage” (Raphan et al. 1979, Laurens and Angelaki 2011). Importantly, the contribution of this signal is significant for movements larger than ∼1s, particularly during high velocity rotations.
- The feedback to tilt 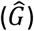 converts canal errors into a tilt velocity (*dG*/*dt*) signal, which is subsequently integrated by the internal model of head tilt.

The non-negligible feedback signals originating from the otolith error *δF* are as follows (Fig. 9, right):

- The feedback to linear acceleration 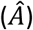 converts unexpected otolith activation into an acceleration signal, and is responsible for acceleration perception during passive translations (as well as experimentally generated otolith errors; Merfeld et al. 1999, Laurens et al. 2013a).
- The *δF* feedback to tilt 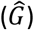 implements the somatogravic effect that acts to bias the internal estimate of gravity towards the net otolith signal so as to reduce the otolith error.
- The *δF* feedback to 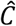 plays a similar role with the feedback to tilt 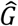, i.e. to reduce the otolith error; but acts indirectly by biasing the internal estimate of rotation in a direction which, after integration, drives the internal model of tilt so that it matches otolith signal (this feedback was called ‘velocity feedback’ in Laurens and Angelaki 2011). Behavioral studies (and model simulations) indicate that this phenomenon has low-frequency dynamics and results in the ability of otolith signals to estimate rotational velocity (Angelaki and Hess, 1996; Hess and Angelaki, 1993). Lesion studies have demonstrated that this feedback depends on an intact nodulus and ventral uvula, the vermal vestibulo-cerebellum (Angelaki and Hess, 1995a,b).

The model in Fig. 9 is entirely compatible with previous models based on optimal passive self-motion computations (Oman 1982; Borah et al. 1988; Merfeld 1995; Laurens 2006; Laurens and Droulez 2007, 2008; Laurens and Angelaki 2011; Karmali and Merfeld 2012; Lim et al. 2017; Zupan and Merfeld, 2002). The present model is, however, distinct in two very important aspects: First, it takes into account active motor commands and integrates these commands with the vestibular sensory signals. Second, because it is formulated as a Kalman filter, it makes specific predictions about the feedback error signals, which constitute the most important nodes in understanding the neural computations underlying head motion sensation. Indeed, as will be summarized next, the properties of most cell types in the vestibular and cerebellar nuclei, as well as the vestibulo-cerebellum, appear to represent either sensory error or feedback signals.

#### Vestibular and rostral fastigial neurons encode sensory error or feedback signals during rotation and translation

Multiple studies have reported that vestibular-only (erroneous word to describe ‘non-eye-movement-sensitive’) neurons in the vestibular nuclei (VN) encode selectively passive head rotation (McCrea and Luan 2003; Roy and Cullen 2001; 2004; Brooks and Cullen 2014) or passive translation (Carriot et al. 2013), but suppress this activity during active head movements. In addition, a group of rostral fastigial nuclei (unimodal rFN neurons; Brooks and Cullen 2013; Brooks et al. 2015) also selectively encodes passive (but not active) rotations. These rotation-responding VN/rFN neurons likely encode either the semicircular canal error *δV* itself or its Kalman feedback to the rotation estimate (blue in Fig. 9, dashed and solid ovals ‘VN, rFN’, respectively). The translation-responding neurons likely encode either the otolith error *δF* or its feedback to the linear acceleration estimate (Fig. 9, solid and dashed red lines ‘VN, trans PC’). Because error and feedback signals are proportional to each other in the experimental paradigms considered here, whether VN/rFN encode sensory errors or feedback signals cannot easily be distinguished using vestibular stimuli alone. Nevertheless, it is also important to emphasize that, while the large majority of VN and rFN neurons exhibit reduced responses during active head movements, this suppression is rarely complete (McRea et al. 1999; Roy and Cullen 2001; Brooks and Cullen 2013; Carriot et al. 2015). Thus, neuronal responses likely encode mixtures of error/feedback and sensory motion signals (e.g., such as those conveyed by direct afferent inputs).

During large amplitude passive rotations (Figure 4 Suppl. 3), the rotation estimate persists longer than the vestibular signal (Fig. 4, blue; a property called velocity storage). Because the internal estimate is equal to the canal error, this implies that vestibular nuclei neurons (that encode the canal error) should exhibit dynamics that are different from those of canal afferents, having incorporated velocity storage signals. This has indeed been demonstrated in VN neurons during optokinetic stimulation (Fig. 4 Suppl. 1; Waespe and Henn 1977) and rotation about tilted axes (Fig. 6 Suppl. 3; Reisine and Raphan 1992).

#### Thalamus-projecting VN neurons possibly encode final motion estimates

Based on the work summarized above, the final estimates of rotation (Fig. 4G) and translation (Fig. 6G), which are the desirable signals to drive perception, do not appear to be encoded by most VN/rFN cells. Thus, one may assume that they are reconstructed downstream, perhaps in thalamic (Marlinski and McCrea, 2008; Meng et al., 2007; Meng and et al., 2010; but see Dale and Cullen 2016) or cortical areas. Interestingly, more than half (57%) of the VN cells projecting to the thalamus respond similarly during passive and actively-generated head rotations (Marlinski and McCrea, 2009). The authors emphasized that VN neurons with attenuated responses during actively-generated movements constitute only a small fraction (14%) of those projecting to the thalamus. Thus, although abundant in the VN, these passive motion-selective neurons may carry sensory error/feedback signals to the cerebellum, spinal cord or even other VN neurons (e.g., those coding the final estimates; Marlinski and McCrea, 2009). VN neurons identified physiologically to project to the cervical spinal cord do not to modulate during active rotations, so they could encode either passive head rotation or active and passive trunk rotation (McCrea et al., 1999). Thus, no firm conclusions can be drawn.

Furthermore, the dynamics of the thalamus-projecting VN neurons with similar responses to passive and active stimuli were not measured (Marlinski and McCrea, 2009). Recall that the model predicts that final estimates of rotation differ from canal afferent signals only in their response dynamics (Fig. 4, compare panels F and G). It would make functional sense that these VN neurons projecting to the thalamus follow the final estimate dynamics (i.e., they are characterized by a prolonged time constant compared to canal afferents) – and future experiments should investigate this hypothesis. Therefore, we would like to emphasize how efficient passive vestibular stimuli, even though those that may seem unnatural, are for computational insights and for understanding the functional properties of the neural network.

#### Rostral fastigial neurons encoding passive trunk rotations

Another class of rFN neurons (and possibly VN neurons projecting to the thalamus; Marlinski and McCrea, 2009, or those projecting to the spinal cord; McCrea et al., 1999) specifically encodes passive trunk velocity in space, independently of head velocity (bimodal neurons; Brooks and Cullen 2009, 2013, Brooks et al. 2015). These neurons likely encode Kalman feedback signals about trunk velocity (Fig. 7, blue). Importantly, these neurons respond equivalently to passive whole trunk rotation when the trunk and the head rotate together (Fig. 7A) and to passive trunk rotation when the head is space-fixed (Fig. 7C). The first protocol activates the semicircular canals and induces a canal error *δV*, while the later activates neck proprioceptors and generates a proprioceptive error, *δP*. From a physiological point of view, this indicates that bimodal neurons respond to semicircular canals as well as neck proprioceptors (hence their name).

The Kalman filter also predicts that neck proprioceptive signals that encode neck position should be transformed into error signals that encode neck velocity. In line with model predictions, bimodal neurons encode velocity signals that originate from neck proprioception during passive sinusoidal (1Hz, Brooks and Cullen 2009) and transient (Gaussian velocity profile, Brooks and Cullen 2013) movements. Remarkably, although short-duration rotation of the trunk while the head is stationnary in space leads to a veridical perception of trunk rotation, long duration trunk rotation leads to an attenuation of the perceived trunk rotation and a growing illusion of head rotation in opposite direction (Mergner et al., 1991). These experimental findings are also predicted by the Kalman filter model (Fig. 7, Suppl. 5).

#### Purkinje cells in the vestibulo-cerebellum encode tilt and acceleration feedback

The simple spike modulation of two distinct types of Purkinje cells in the caudal cerebellar vermis (lobules IX-X, Uvula and Nodulus) encodes tilt (tilt-selective cells) and translation (translation-selective cells) during three-dimensional motion (Yakusheva et al., 2007; 2008; 2013; Laurens et al. 2013a,b). Therefore, it is possible that tilt- and translation selective cells encode tilt and acceleration feedbacks (Fig. 9, green and red lines, respectively). If so, we hypothesize that their responses are suppressed during active motion (Fig. 5 and 6). How Purkinje cells modulate during active motion is currently unknown. However, one study (Lee et al. 2015) performed when rats learned to balance on a swing indicates that Purkinje cell responses that encode trunk motion are reduced during predictable movements, consistent with the hypothesis that they encode sensory errors or Kalman feedback signals.

Model simulations have also revealed that passive tilt doesn’t induce any significant otolith error (Fig. 5J). In contrast, passive tilt elicits a significant canal error (Fig. 5K). Thus, we hypothesize that the tilt signal present in the responses of Purkinje cells originates from the canal error *δV* onto the tilt internal state variable. If it is indeed a canal, rather than an otolith, error, it should be proportional to tilt velocity instead of tilt position (or linear acceleration). Accordingly, we observed (Laurens et al. 2013b) that tilt-selective Purkinje cell responses were on average close to velocity (average phase lag of 36° during sinusoidal tilt at 0.5Hz). However, since sinusoidal stimuli are not suited for establishing dynamics (Laurens et al., 2017), further experiments are needed to confirm that tilt-selective Purkinje cells indeed encode tilt velocity.

Model simulations have also revealed that passive translation, unlike passive tilt, should include an otolith error. This otolith error feeds also into the tilt internal variable (Fig. 9, somatogravic feedback) and is responsible for the illusion of tilt during sustained passive linear acceleration (somatogravic effect; Graybiel 1952). Therefore, as summarized in Fig. 9 (green lines), both canal and otolith errors should feedback onto the tilt internal variable. The canal error should drive modulation during tilt, whereas the otolith error should drive modulation during translation. In support of these predictions, we have demonstrated that tilt-selective Purkinje cells also modulate during translation, with a gain and phase consistent with the simulated otolith-driven feedback (Laurens et al 2013b). Thus, both of these feedback error signals might be carried by caudal vermis Purkinje cells – and future experiments should address these predictions.

Note that semicircular canal errors must be spatially transformed in order to produce an appropriate tilt feedback. Indeed, converting a rotation into head tilt requires taking into account the angle between the rotation axis and earth-vertical. This transformation is represented by a bloc marked “3D” in Fig. 9 (see also (*eq*. 9) in Suppl. Methods, ‘Three-Dimensional Kalman filter’. Importantly, we have established (Laurens et al. 2013b) that tilt-selective Purkinje cells encode spatially transformed rotation signals, as predicted by theory. In fact, we have demonstrated that tilt-selective Purkinje cells do not simply modulate during vertical canal stimulation, but also carry the tilt signal during off-vertical axis yaw rotations (Laurens et al. 2013b).

In this respect, it is important to emphasize that truly tilt-selective neurons exclusively encode changes in orientation relative to gravity, rather than being generically activated by vertical canal inputs. Thus, it is critical that this distinction is experimentally made using three-dimensional motion (see Laurens et al. 2013b, Laurens and Angelaki 2015). Whereas 3D rotations have indeed been used to identify tilt-selective Purkinje cells in the vermis (Laurens et al. 2013b; Yakusheva et al. 2007), this is not true for other studies. For example, Laurens and Angelaki (2015) and Zhou et al. (2006) have reported tilt-modulated cells in the rFN and VN, respectively, but because these neurons were not tested in three dimensions, the signals carried by these neurons remain unclear.

### Further notes on tilt-selective Purkinje cells

As summarized above, the simple spike responses of tilt-selective Purkinje cells during passive motion have already revealed many details of the internal model computations. Thus, we have proposed that tilt-selective Purkinje cells encode the feedback signals about tilt, which includes scaled and processed (i.e. by a spatial transformation, green “3D” box in Fig. 9) versions of both canal and otolith sensory errors (Fig. 9, green oval, ‘tilt PC?’). However, there could be alternative implementations of the Kalman filter, where tilt-selective Purkinje cells may not encode only feedback signals. Next, we propose an alternative formulation regarding the signal they convey and the underlying organization of predictive and feedback pathways.

We note that motor commands *Ω*^*u*^ must be also be spatially processed (black “3D” box in Fig. 9) to contribute to the tilt prediction. One may question whether two distinct neuronal networks transform motor commands and canal errors independently (resulting in two “3D” boxes in Fig. 9). An alternative (Fig. 9 Suppl. 1) would be that the brain merges motor commands and canal error to produce a final rotation estimate prior to performing this transformation. From a mathematical point of view, this alternative would only require a re-arrangement of the Kalman filter equations, which would not alter any of the model’s conclusions. However, tilt-selective Purkinje cells, which encode a spatially transformed signal, would then carry a mixture of predictive and feedback signals and would therefore respond identically to active and passive tilt velocity. Therefore, the brain may perform a spatial transformation of predictive and feedback rotation signals independently; or may merge them before transforming them. Recordings from tilt-selective Purkinje cells during active movements will distinguish between these alternatives.

#### Summary of the neural implementation of sensory error and feedback signals

In summary, many of the response properties described by previous studies for vestibular nuclei and cerebellar neurons can be assigned a functional ‘location’ within the Kalman filter model. Interestingly, most of the central neurons fit well with the properties of sensory errors and/or feedback signals. That an extensive neural machinery has been devoted to feedback signals is not surprising, given their functional importance for self-motion estimation. For many of these signals, a distinction between sensory errors and feedbacks is not easily made. That is, rotation-selective VN and rFN neurons can encode either canal error (Fig. 9, bottom, dashed blue oval) or rotation feedback (Fig. 9, bottom, solid blue oval). Similarly, translation-selective VN, rFN and Purkinje cells can encode either otolith error (Fig. 9, bottom, dashed red oval) or translation feedback (Fig. 9, bottom, solid red oval). The only feedback that is easily distinguished based on currently available data is the tilt feedback (Fig. 9, green lines).

Although the blue, green and red feedback components of Fig. 9 can be assigned to specific cell groups, this is not the case with the cyan feedback components. First, note that, like the tilt variable (but unlike the rotation and translation variables, which receive significant feedback contributions from either only the canal or only the otolith errors, respectively), the canal internal model variable, receives non-negligible feedback contributions from both the canal and otolith sensory errors (Fig. 9, cyan lines). The canal feedback error changes the time constant of the rotation estimate (Fig. 4 and Fig. 4, Suppl. 1 and 3), whereas the otolith feedback error may suppress (post-rotatory tilt) or create a rotation estimate (Fig. 6, Suppl. 3). The neuronal implementation of the internal model of the canals 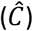, and of its associated feedback pathways, are currently unknown. However, lesion studies clearly indicate that the caudal cerebellar vermis, lobules X and IX might influence the canal internal model state variable (Angelaki and Hess 1995a,b; Wearne et al., 1998). In fact, it is possible that the simple-spike output of the translation-selective Purkinje cells also carries the otolith sensory error feedback to the canal internal model state variable (Fig. 9, bottom, cyan arrow passing though the dashed red ellipse). Similarly, the canal error feedback to the canal internal model state variable (Fig. 9, bottom, cyan arrow originating from the dashed blue ellipse) can originate from VN or rFN cells that selectively encode passive, not active, head rotation (Fig. 4J, note that the *C*^*k*^ feedback is but a scaled-down version of the *Ω*^*k*^ feedback).

Thus, although the feedback error signals to the canal internal model variable can be linked to known neural correlates, cells coding for the state variable 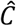 exclusively have not been identified. It is possible that the representation of the hidden variable 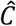 may be coded in a distributed fashion. After all, as already stated above, VN and rFN neurons have also been shown to carry mixed signals - they can respond to both rotation and translation, as well as they may carry both feedback/error and actual sensory signals. Thus, it is important to emphasize that these Kalman variables and error signals may be represented in a multiplexed way, where single neurons manifest mixed selectivity to more than just one internal state and/or feedback signals. This appears to be an organizational principle both in central vestibular areas (Laurens et al., 2017) and throughout the brain (Rigotti et al., 2013, Fusi et al., 2016). It has been proposed that mixed selectivity has an important computational advantage: high-dimensional representations with mixed selectivity allow a simple linear readout to generate a diverse array of potential responses (Fusi et al., 2016). In contrast, representations based on highly specialized neurons are low dimensional and may preclude a linear readout from generating several responses that depend on multiple task-relevant variables.

### Interruption of internal model computations during proprioceptive mismatch

In this treatment, we have considered primarily the importance of the internal model of the sensors to emphasize its necessity for both self-generated motor commands and unpredicted, external perturbations. It is important to also point out that self-generated movements also utilize an internal model of the motor plant which is modeled by a matrix M (as well as internal dynamics modeled by matrix D) in Fig. 1. In fact, it is this motor internal model that has been studied and shown to be altered in experiments where resistive torques are applied to the head (Brooks et al., 2015; Brooks and Cullen, 2015; Cullen and Brooks, 2015) or active movements are entirely blocked (Roy and Cullen, 2001; 2004; Carriot et al. 2013). Under these conditions, central neurons were shown to encode net head motion (i.e. active and passive indiscriminately) with a similar gain as during passive motion. This indicates that the brain ceases to subtract predicted sensory signals from afferent inputs whenever head motion is perturbed or, in other words, ceases to use an internal model of the motor plant when this model appears to be incorrect. This result may be reproduced by the Kalman filter by switching off the internal model of the motor plant (i.e. setting the matrix M to zero) when active movements are perturbed (Fig. 7 Suppl. 6C). Note that the mechanism that operates this switch is not included in the Kalman filter framework in Fig. 1.

Roy and Cullen (2004) have shown that this process is triggered by proprioceptive mismatch: Perturbing neck or body motion induces a discrepancy between the intended head position and proprioceptive feedback. Note that perturbing head motion also induces a vestibular mismatch since it causes the head velocity in space to differ from the motor plan. However, vestibular mismatch, which also occurs when passive rotations are superimposed to active movements, don’t interrupt internal model computations, as shown experimentally by (Brooks and Cullen 2013, 2014; Carriot et al. 2013) and illustrated in the model predictions of Fig. 8.

Therefore, proprioceptive mismatch is likely a specific indication that the internal model of the motor plant (matrix M in Fig. 1) is inaccurate. Instead of using an inaccurate model, the brain stops using it altogether until an accurate model is learned (Brooks et al., 2015; Brooks and Cullen, 2015; Cullen and Brooks, 2015). Note that the computation that detects proprioceptive mismatch may occur separately from the Kalman filter.

### Relation to previous three-dimensional models and Bayesian Inference

For simplicity, we have presented a linearized one-dimensional model in this study. Three-dimensional Kalman filter models can also be constructed (Borah et al. 1988; Lim et al. 2017), and we show in Suppl. Methods, ‘Three-dimensional Kalman filter’, how to generalize the model to three dimensions. However, to simplify the main framework and associated predictions, we presented the result of a simpler one-dimensional model.

The passive motion components of the model presented here is to a large extent identical to the Particle filter Bayesian model in (Laurens 2006, Laurens and Droulez 2007; 2008; Laurens and Angelaki 2011), which we have re-implemented as a Kalman filter, and into which we incorporated motor commands. One fundamental aspect of previous Bayesian models (Laurens 2006; Laurens and Droulez 2007; 2008) is the explicit use of two Bayesian priors that prevent sensory noise from accumulating over time. These priors encode the natural statistics of externally generated motion or motion resulting from motor errors and unexpected perturbations. Because, on average, rotation velocities and linear accelerations are close to zero, these Bayesian priors are responsible for the decrease of rotation estimates during sustained rotation (Fig. 4 Suppl. 2) and for the somatogravic effect (Fig. 6 Suppl. 2) (see Laurens and Angelaki 2011 for further explanations). The influence of the priors is higher when the statistical distributions of externally generated rotation (*Ω*^*ε*^) and acceleration (*A*^*ε*^) is narrower (Fig. 9 Suppl. 2), i.e. when their standard deviation is smaller. Stronger priors reduce the gain and time constant of rotation and acceleration estimates (Fig. 9 Suppl. 2). Importantly, the Kalman filter model predicts that the priors affect only the passive, but not the active, self-motion final estimates. Indeed, the rotation and acceleration estimates last indefinitely during simulated active motion (Fig. 4 Suppl. 2, Fig. 6 Suppl. 2). In this respect, the Kalman filter explains why the vestibulo-ocular reflex is reduced in figure ice skaters (Tanguy et al. 2008; Alpini et al. 2009): The range of head velocities experienced in these activities is wider than normal. In previous Bayesian models, we found that that widening the rotation prior should increase the time constant of vestibular responses, apparently in contradiction with these results. However, these models didn’t consider the difference between active and passive stimuli. The formalism of the Kalman filter reveals that Bayesian priors should reflect the distribution of passive motion or motor errors. In athletes that are highly trained to perform stereotypic movements, this distribution likely narrows, resulting in stronger priors and reduced vestibular responses.

### Role of the vestibular system during active motion: ecological, clinical and fundamental implications

Neuronal recordings (Brooks and Cullen 2013, 2014; Carriot et al. 2013) and the present modeling unambiguously demonstrate that central neurons respond to unexpected motion during active movement (a result that we reproduced in Fig. 8G-I). Beyond experimental manipulations, a number of processes may cause unpredictable motion to occur in natural environments. When walking on tree branches, boulders or soft grounds, the support surface may move under the feet, leading to unexpected trunk motion. A more dramatic example of unexpected trunk motion, that requires immediate correction, occurs when slipping or tripping. Complex locomotor activities involve a variety of correction mechanism among which spinal mechanisms and vestibular feedback play preeminent roles (Keshner et al. 1987; Black et al. 1988; Horstmann and Dietz 1988).

The contribution of the vestibular system for stabilizing posture is readily demonstrated by considering the impact of chronic bilateral vestibular deficits. While most patients retain an ability to walk on firm ground and even perform some sports (Crawford 1964; Herdman 1996), vestibular deficit leads to an increased incidence of falls (Herdman et al. 2000), difficulties in walking on uneven terrains and deficits in postural responses to perturbations (Keshner et al. 1987; Black et al. 1988; Riley 2010). This confirms that vestibular signals are important during active motion, especially in challenging environments. In this respect, the Kalman filter framework appears particularly well suited for understanding the effect of vestibular lesions.

Vestibular sensory errors also occur when the internal model of the motor apparatus is incorrect (Brooks and Cullen 2015) and these errors can lead to recalibration of this internal model. This suggests that vestibular error signals during self-generated motion may play two fundamental roles: (1) updating self-motion estimates and driving postural or motor corrections, and (2) providing teaching signals to internal models of motor control (Wolpert et al 1995) and therefore facilitating motor learning.

But perhaps most importantly, the model presented here should eliminate the misinterpretation that vestibular signals are ignored during self-generated motion – and that passive motion stimuli are old-fashioned and should no longer be used in experiments. Regarding the former conclusion, the presented simulations highlight the role of the internal models of canal dynamics and otolith ambiguity, which operate continuously to generate the *correct sensory prediction* during *both* active and passive movements. Without these internal models, the brain would be unable to correctly predict sensory canal and otolith signals and everyday active movements would lead to sensory mismatch (e.g., for rotations, see Fig. 4 Suppl. 2 and 3). Thus, even though particular nodes (neurons) in the circuit show attenuated or no modulation during active head rotations, vestibular processing remains the same - the internal model is both engaged and critically important for accurate self-motion estimation, even during actively-generated head movements. Regarding the latter conclusion, it is important to emphasize that passive motion stimuli have been, and continue to be, extremely valuable in revealing salient computations that would have been amiss if the brain’s intricate wisdom was interrogated only with self-generated movements.

### Conclusion

“A good model has a delightful way of building connections between phenomena that never would have occurred to one” (Robinson, 1977). Four decades later, this beautifully applies here, where the mere act of considering how the brain should process self-generated motion signals in terms of mathematical equations (instead of schematic diagrams) immediately revealed a striking similarity with models of passive motion processing and, by motivating this work, opened an avenue to resolve a standing paradox in the field.

The framework of the internal model hypothesis, and the series of quantitative models it has spawned, have explained and simulated behavioral and neuronal responses to a long list of passive motion paradigms, and with a spectacular degree of accuracy (Merfeld 1995; Merfeld et al. 1999; Angelaki et al. 2004; Laurens et al. 2010, 2011a; Laurens et al. 2013a,b; Lim et al. 2017). The present study offers the theoretical framework which will likely assist in understanding neuronal computations that are essential to active self-motion perception, balance and locomotor activity in everyday life.

## Methods

### Structure of a Kalman filter

In a Kalman filter (Kalman 1960), state variables *X* are driven by their own dynamics (matrix *D*), motor commands *X*^*u*^ and internal or external perturbations *X*^*ε*^ through the equation (Fig. 1A):

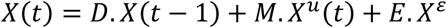

A set of sensors, grouped in a variable *S*, measures state variables or combinations thereof (encoded in a matrix *T*), and are modeled as:

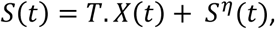

where *S*^*η*^ is Gaussian sensory noise. The model assumes that the brain has an exact knowledge of the forward model, i.e. of *D*, *M*, *E* and *T* as well as the variances of *X*^*ε*^ and *S*^*η*^. Furthermore, the brain knows the values of the motor inputs *X*^*u*^ and sensory signals *S*, but doesn’t have access to the actual values of *X*^*ε*^ and *S*^*η*^.

At each time *t*, the Kalman filter computes a preliminary estimate (also called a prediction) 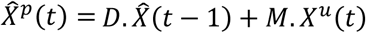 and a corresponding predicted sensory signal 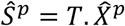 (Fig. 1B). This prediction 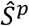 and the sensory input *S* are compared to compute a sensory error *δS*. Sensory errors are then transformed into a feedback *X*^*k*^ = *K*. *δS* where *K* is a matrix of feedback gains. Thus, an improved estimate at time *t* is 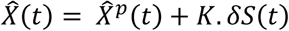. The value of the feedback gain matrix *K* determines how sensory errors (and therefore sensory signals) are used to compute the final estimate 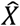, and is computed based on *D*, *E*, *T* and on the variances of *X*^*ε*^ and *S*^*η*^ (see Suppl. Methods, ‘Kalman filter algorithm’).

In the case of the self-motion model, the motor commands *Ω*^*u*^ and *A*^*u*^ are inputs to the Kalman filter (Fig. 2). Note that, while the motor system may actually control other variables (such as forces or accelerations), we consider that these variables are converted into *Ω*^*u*^ and *A*^*u*^. We demonstrate in Suppl. Methods, ‘Model of motor commands’ that altering these assumptions does not alter our conclusions. In addition to motor commands, a variety of internal or external factors such as external (passive) motion also affect *Ω* and *A*. The total rotation and acceleration components resulting from these factors are modeled as variables *Ω*^*ε*^ and *A*^*ε*^. Similar to (Laurens 2006, Laurens and Droulez 2007, 2008) we modeled the statistical distribution of these variables as Gaussians, with standard deviations *σ*_*Ω*_and *σ*_*A*_.

Excluding vision and proprioception, the brain senses head motion though the semicircular canals (that generate a signal *V*) and the otoliths organs (that generate a signal *F*). Thus, in initial simulations (Fig. 3-6), the variable *S* encompasses *V* and *F* (neck proprioceptors are added in Fig. 7).

The semicircular canals are rotation sensors that, due to their mechanical properties, exhibit high-pass filter properties. These dynamics may be neglected for rapid movements of small amplitude (such as Fig. 3) but can have significant impact during natural movements (Fig. 4 Suppl. 3). They are modeled using a hidden state variable *C*. The canals are also subject to sensory noise *V*^*η*^. Taken both the noise and the dynamics into account, the canals signal is modeled as *V* = *Ω*−*C* + *V*^*η*^.

The otolith organs are acceleration sensors. They exhibit negligible temporal dynamics in the range of motion considered here, but are fundamentally ambiguous: they sense gravitational as well as linear acceleration – a fundamental ambiguity resulting from Einstein’s equivalence principle (Einstein 1907). Gravitational acceleration along the inter-aural axis depends on head roll position, modeled here as 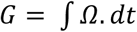. The otoliths encode the sum of *A* and *G* and is also affected by sensory noise *F*^*η*^, such that the net otolith signal is *F* = *A* + *G* + *F*^*η*^.

How sensory errors are used to correct motion estimates depends on the Kalman gain matrix, which is computed by the Kalman algorithm such that the Kalman filter as a whole performs optimal estimation. In theory, the Kalman filter includes a total of 8 feedback signals, corresponding to the combination of 2 sensory errors (canal and otolith errors) and 4 internal states (see Suppl. Methods, ‘Kalman feedback gains’).

It is important to emphasize that the Kalman filter model is closely related to previous models of vestibular information processing. Indeed, simulations of long-duration rotation and visuo-vestibular interactions (Fig. 4 Suppl. 2), as well as mathematical analysis (Laurens 2006), demonstrate that 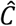 is equivalent to the “velocity storage” (Raphan et al. 1979, Laurens and Angelaki 2011). These low-frequency dynamics, as well as visuo-vestibular interactions, were previously simulated and interpreted in the light of optimal estimation theory; and accordingly are reproduced by the Kalman filter model.

The model presented here is to a large extent identical to the Particle filter Bayesian model in (Laurens 2006, Laurens and Droulez 2007; 2008; Laurens and Angelaki 2011). It should be emphasized that: (1) transforming the model into a Kalman filter didn’t alter the assumptions upon which the Particle filter was build; (2) introducing motor commands into the Kalman filter was a textbook process that did not require any additional assumptions or parameters; and (3) we used exactly the same parameter values as in Laurens (2006) and Laurens and Droulez (2008) (with the exception of *σ*_*F*_ whose impact, however, is negligible, and of the model of head on trunk rotation that required additional parameters; see next section).

Beyond the question of active and passive motion, these previous models and other Kalman filter models have been used to demonstrate how optimal inference performed by the Kalman filter improves the accuracy of self-motion estimates, in particular by preventing the accumulation of estimation errors (MacNeilage et al. 2008; Laurens and Angelaki 2011) and reducing sensory perception thresholds (Lim et al. 2017). We have also demonstrated that the Bayesian approach accounts for the effects of semicircular canal lesions (Laurens and Droulez, 2008).

### Simulation parameters

The parameters of the Kalman filter model are directly adapted from previous studies (Laurens 2006; Laurens and Droulez 2008). Tilt angles are expressed in radians, rotation velocities in rad/s, and accelerations in *g* (1 *g* = 9.81 m/s^2^). Note that a small tilt angle α (in radians) result in a gravitational acceleration sin(α)::α (in *g*). For this reason, tilt and linear acceleration variables are expressed in equivalent units, and may be added or subtracted. The standard deviations of the unpredictable rotations (*Ω*^*ε*^) and accelerations (*A*^*ε*^) are set to the standard deviations of the Bayesian a priori in Laurens 2006 and Laurens and Droulez 2008, i.e. *σ*_*Ω*_ = 0.7 rad/s (*Ω*^*ε*^) and *σ*_*A*_ =0.3 *g* (*A*^*ε*^). The standard deviation of the noise affecting the canals (*V*^*η*^) was set to *σ*_v_ =0.175 rad/s (as in Laurens 2006 and Laurens and Droulez 2008). The standard deviation of the otolith noise (*F*^*η*^) was set to *σ*_F_ =0.002 *g* (2 cm/s^2^). We verified that values ranging from 0 to 0.01 *g* had no effect on simulation results. The time constant of the canals was set to τ_c_=4s. Simulations used a time step of *δt* = 0.01s. We verified that changing the value of the time step without altering other parameters had no effect on the results.

We ran simulations using a variant of the model that included visual information encoding rotation velocity. The visual velocity signals were affected by sensory noise with a standard deviation *σ*_*Vis*_ = 0.12 rad/s, as in (Laurens and Droulez 2008).

Another variant modeled trunk in space velocity (*Ω*_*TS*_) and head on trunk velocity (*Ω*_*HT*_) independently. The standard deviations of unpredictable rotations were set to *σ*_*TS*_ = 0.7 rad/s (identical to *σ*_*Ω*_) and *σ*_*HT*_ = 3.5 rad/s. The standard deviation of sensory noise affecting neck afferents was set to *σ*_*P*_ = 0.0017 rad.

For simplicity, all simulations were run without adding the sensory noise *V*^*η*^ and *F*^*η*^. These noise-free simulations are representative of the results that would be obtained by averaging several simulation runs performed with sensory noise (e.g. as in Laurens and Droulez 2007). We chose to present noise-free results here in order to facilitate the comparison between simulations of active and passive motion.

**Table 3:**
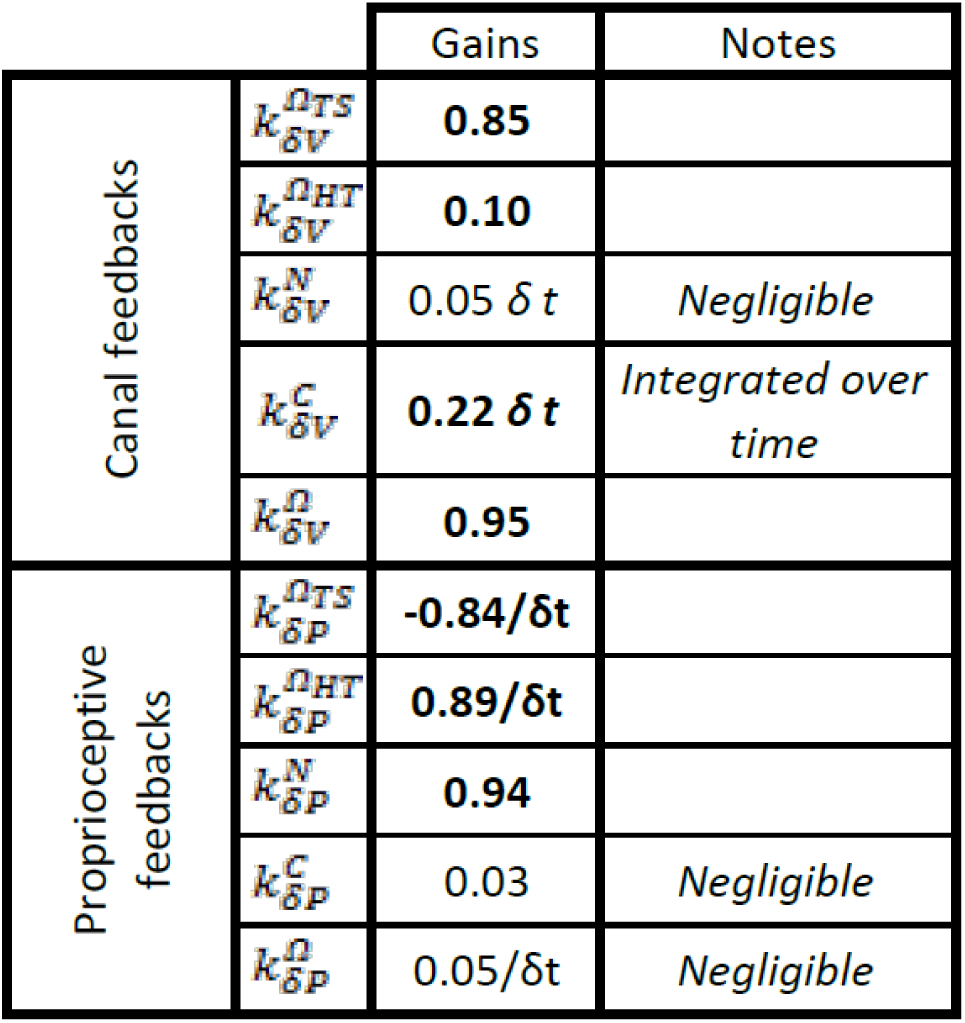
Kalman feedback gains during head and neck rotation. As in Table 2, some feedback gains are constant and independently of *δt*, while some others scale with or inversely to *δt* (see Suppl. Methods, “Feedback gains of the model of head and neck motion” for explanations). Gains that have negligible impact on the motion estimates are indicated in normal fonts, others with profound incluence are indicated in bold. The feedback gains 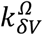 and 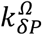 are computed as 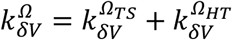 and 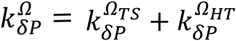.

**Figure 4 Supplement 1.**
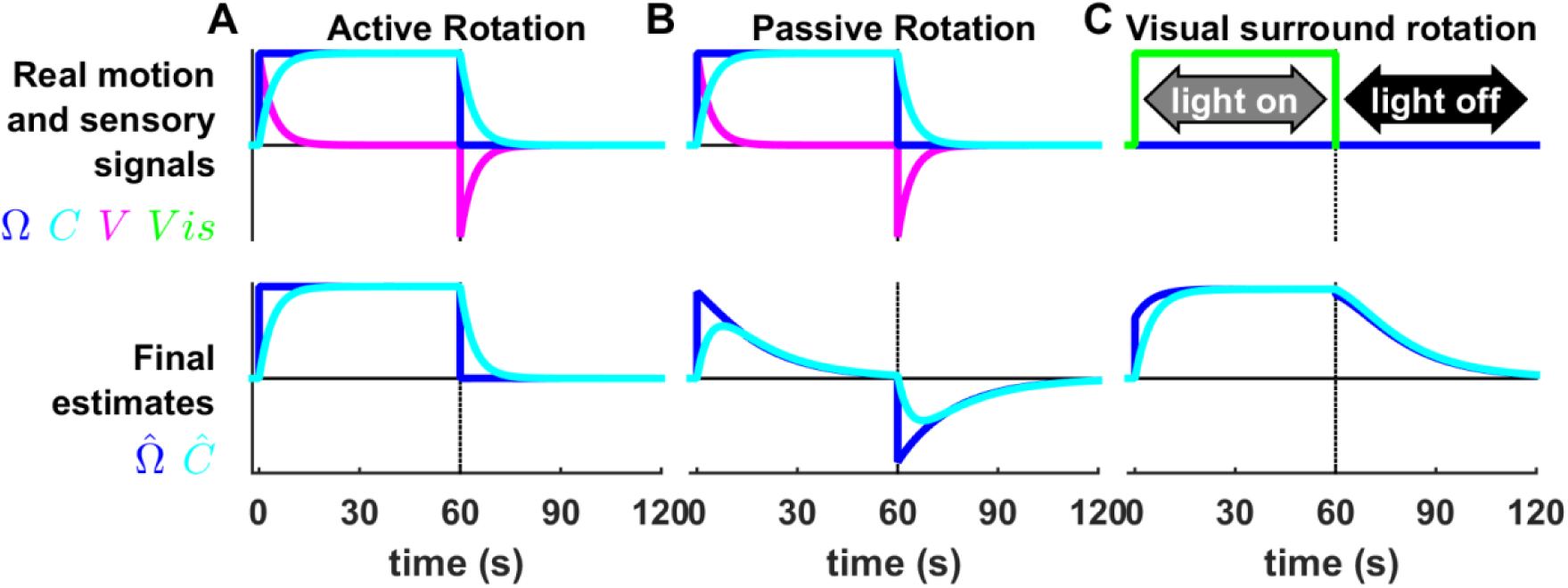
Processing of rotation information during long-duration motion. (A,B) Simulation of constant velocity rotation lasting 60s, followed by 60s where the head doesn’t rotate (top row, *Ω*, blue). The semicircular canal signal (top row, V, magenta) decreases exponentially (time constant=4s) during the rotation (*V* = *Ω* − *C*; *C* represented by cyan lines). The deceleration at t=60s induces a canal after-effect. During active rotation (A), the final estimates of rotation velocity (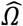 lower row, blue) and canal dynamics (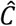 lower row, cyan) are identical to their real respective values (A, top panel). In contrast, during passive rotation (B), the final estimate of rotation differs from the stimulus: 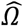 decreases exponentially towards zero, although the decrease is slower (time constant = 16.5s) than that of the canals. This prolonged estimate arises from the contribution of the internal state variable 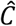 which rises at the beginning of rotation as in (A) but reaches a maximum and decreases towards zero. (C) Simulation of optokinetic stimulation, i.e., a visual stimulus rotating at constant velocity (*Vis*, top row, green) while the head is immobile. The rotation estimate 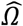 rises immediately to 70% of stimulus velocity at the beginning of the stimulation, and then increases exponentially to 96% of stimulus velocity. 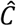 rises exponentially to the stimulation velocity but doesn’t exhibit an immediate increase at the beginning of the stimulation. Both 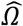 and 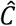 persist at the end of the rotation.

The simulated results in B and C match the perceptual (Bertolini et al. 2011) and reflex (Raphan et al. 1979) responses to these stimuli. Previous work (Raphan et al. 1979; Laurens and Angelaki 2011) used an intermediate variable called ‘velocity storage’ to model low frequency responses to vestibular and visual stimulation. The state variable 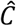 in the Kalman filter model is identical to the velocity storage. An optimal model based on particle filtering also produced identical results (Laurens and Droulez 2007, 2008).

The Kalman filter also predicts that rotation perception should last indefinitely during active constant-velocity rotation (A), unlike passive rotation (B). Note however that sustained rotation perception during active rotation is also accomplished by the velocity storage 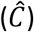, which likely saturates at velocities of ∼20-30°/s in humans (Laurens et al. 2011b). Therefore, actively rotating at substantially higher velocities may lead to disorientation and vertigo (as commonly experienced by children and waltz dancers). Note that all Kalman filter simulations do not include this saturation.

**Figure 4 Supplement 2:**
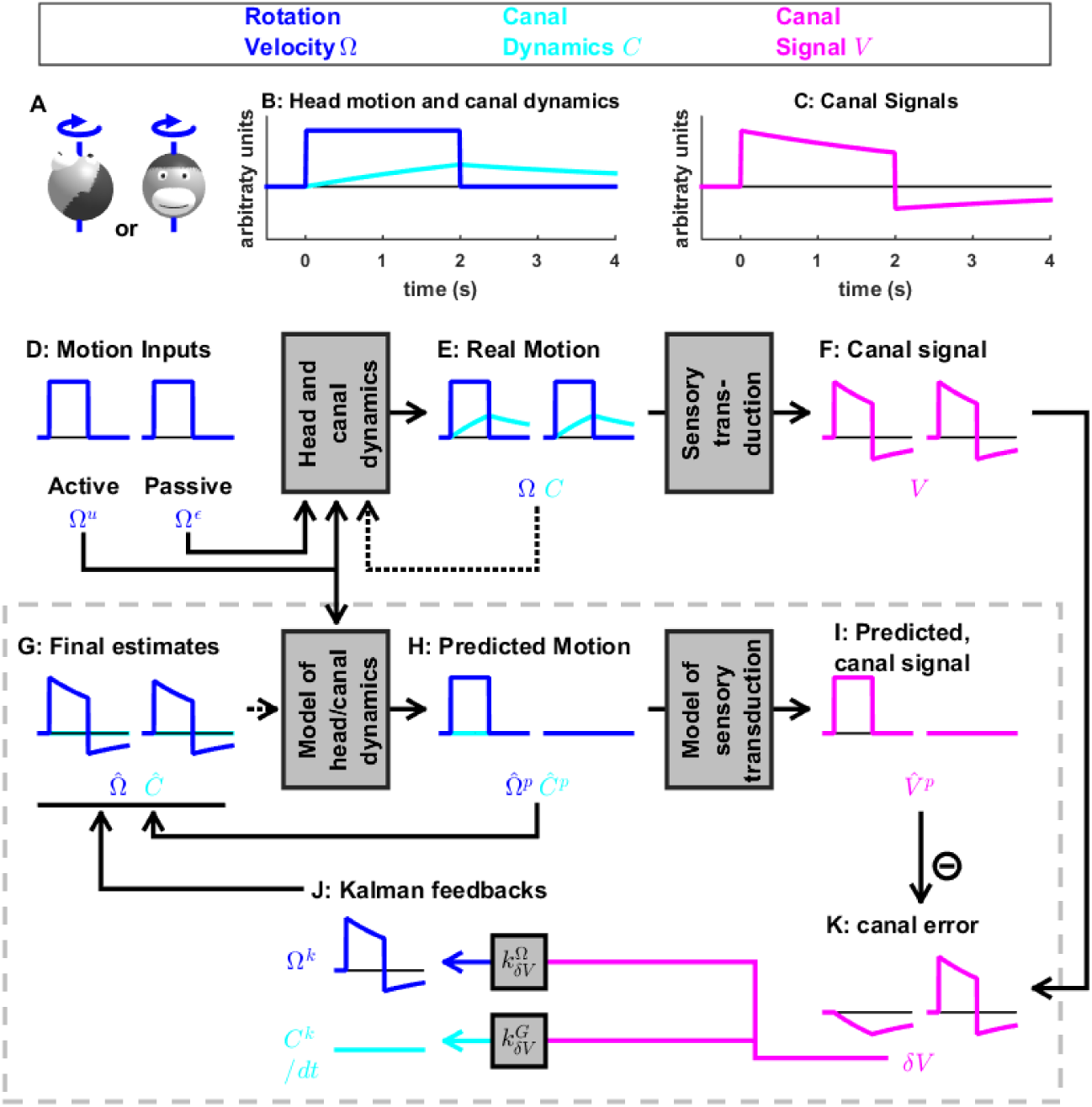
Same simulation as in Fig. 4, where the internal model of canals dynamic is not used. Note error in the final estimate of rotation (G) during both active and passive movements.

**Figure 4 Supplement 3:**
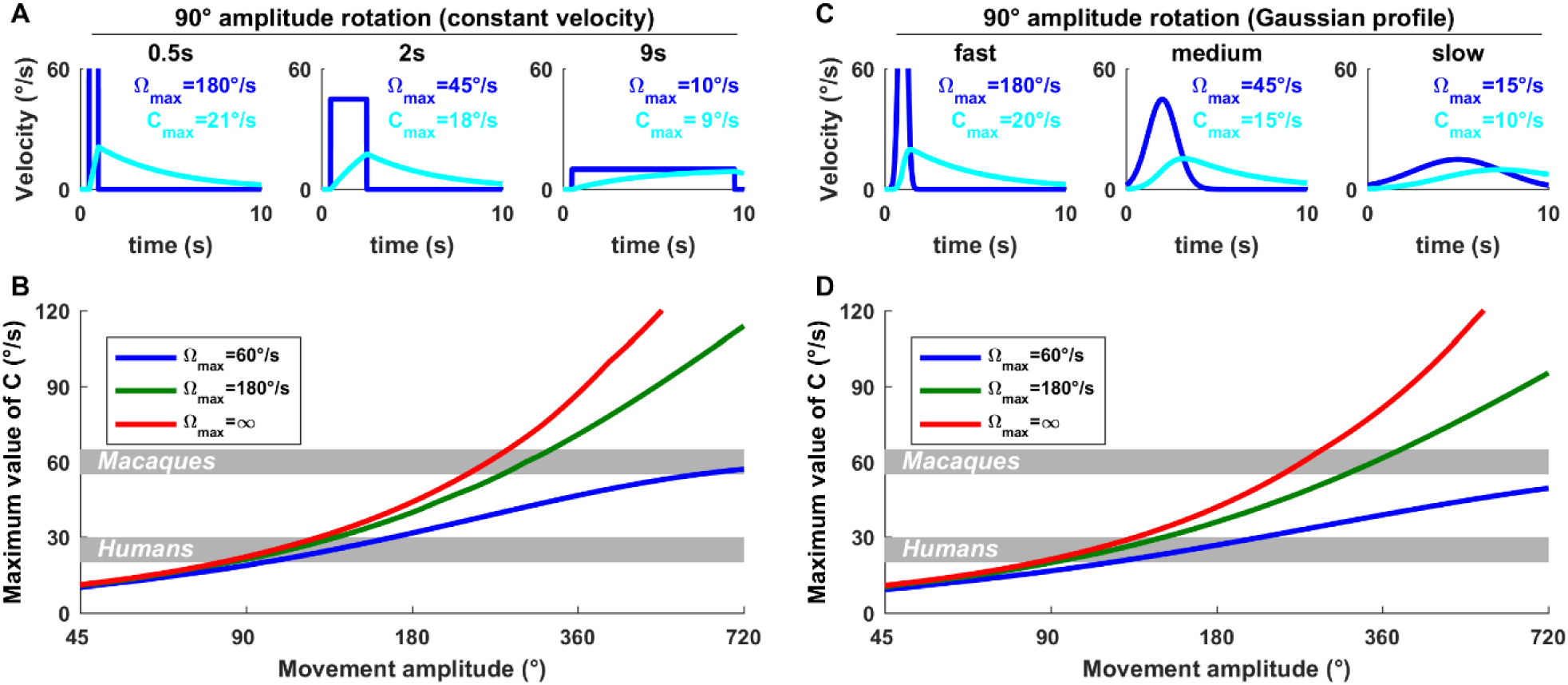
Quantitative analysis of the importance of an internal model of the canals for ecological movements. The internal variable *C*, which represents an internal model of the canals, has low-pass dynamics. Nevertheless, its importance is highest for high velocities. To illustrate this, here we simulate *C* during either (A,B) constant velocity or (C,D) Gaussian-velocity rotations of various amplitudes and durations. (A,C) Rotations with an amplitude of 90° and duration of 0.5, 2 and 9s (i.e. rotations at 180, 45 and 10°/s, respectively). We find that the peak value of *C* is 21°/s during a fast movement (0.5s) whereas it is only 9°/s during a slow movement (9s). Therefore, for a given movement amplitude, the internal model contribution is larger at larger velocities. (B,D) Maximum value of *C* as a function of movement amplitude for slow (60°/s) and rapid (180°/s, and infinite velocity) movements. During fast movements, *C* may exceed 20°/s during 90° rotation and 40°/s during 180° rotation. We note that the velocity storage likely saturates at ∼20-30°/s in humans (Laurens et al. 2011b) and ∼ 60°/s in macaques (Waespe et al. 1983) (shown in B and D as gray bands). (this saturation is not included in the Kalman filter). For fast rotations (180°/s), the saturation would be reached for rotations of 130° in humans (assuming a saturation velocity of 30°/s) and 290° in macaques. These simulations demonstrate that the role of the velocity storage in compensating for canal dynamics is not restricted to long-duration rotations but encompasses natural movements.

**Figure 5 Supplement 1:**
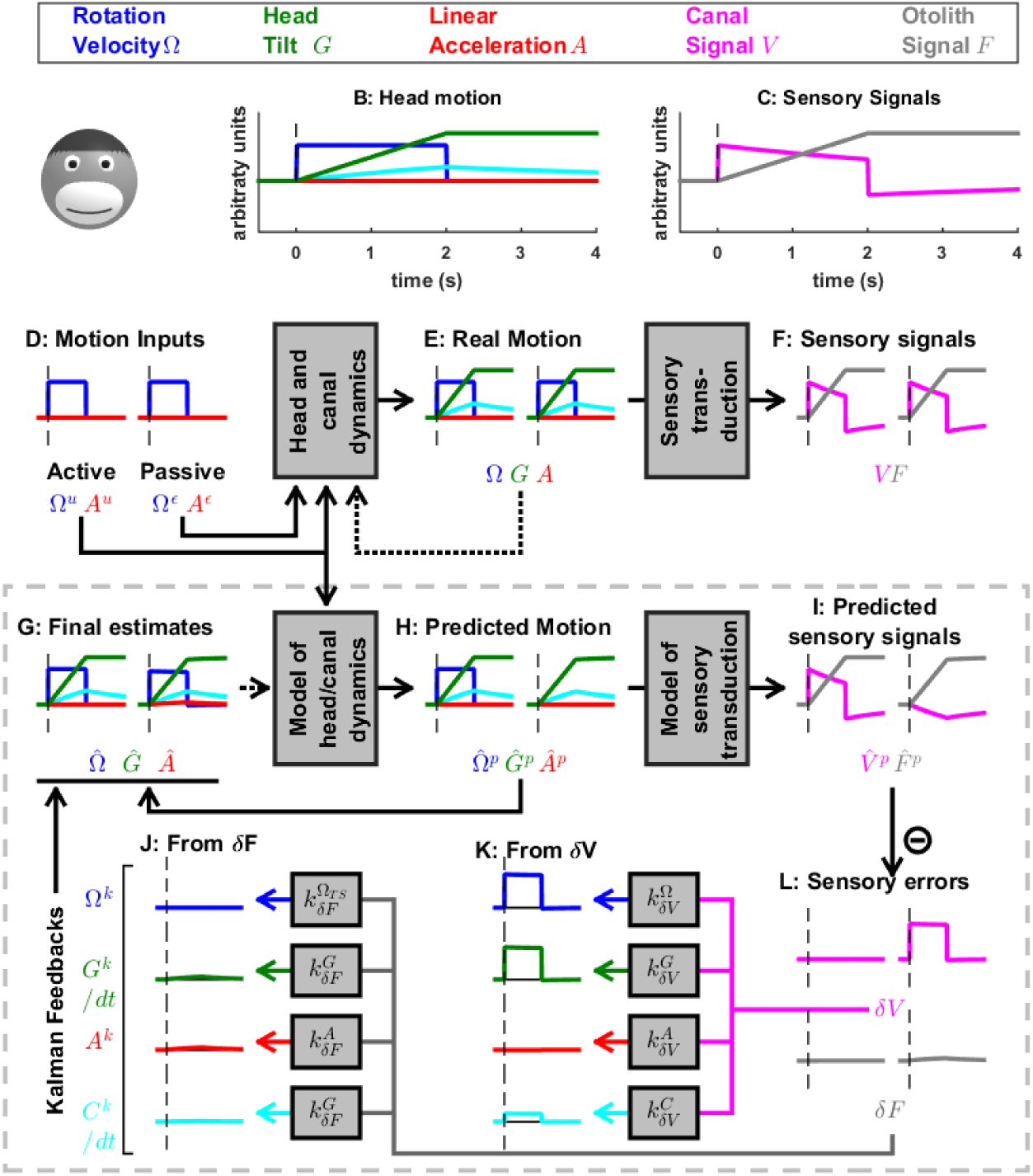
Simulation of medium duration (2s) head tilt movement. Panels as in Fig. 5; the canal dynamics state variable, *C*, is shown in cyan. Although canal responses exhibit significant dynamics (magenta in panel C), rotation perception remains accurate (panel G, blue traces) during both active and passive tilt (as in Fig. 4). The final estimate of tilt (panel G, green traces) is also accurate during active and passive tilt. As a consequence, the otolith error (panel L, gray traces) and the corresponding Kalman feedback (J) are negligible. Note that the feedbacks to *G*^*k*^ and *C*^*k*^ are scaled by a factor 1/*δt* (because of the integration; see Suppl. Methods, ‘Kalman feedback gains’).

**Figure 6 Supplement 1:**
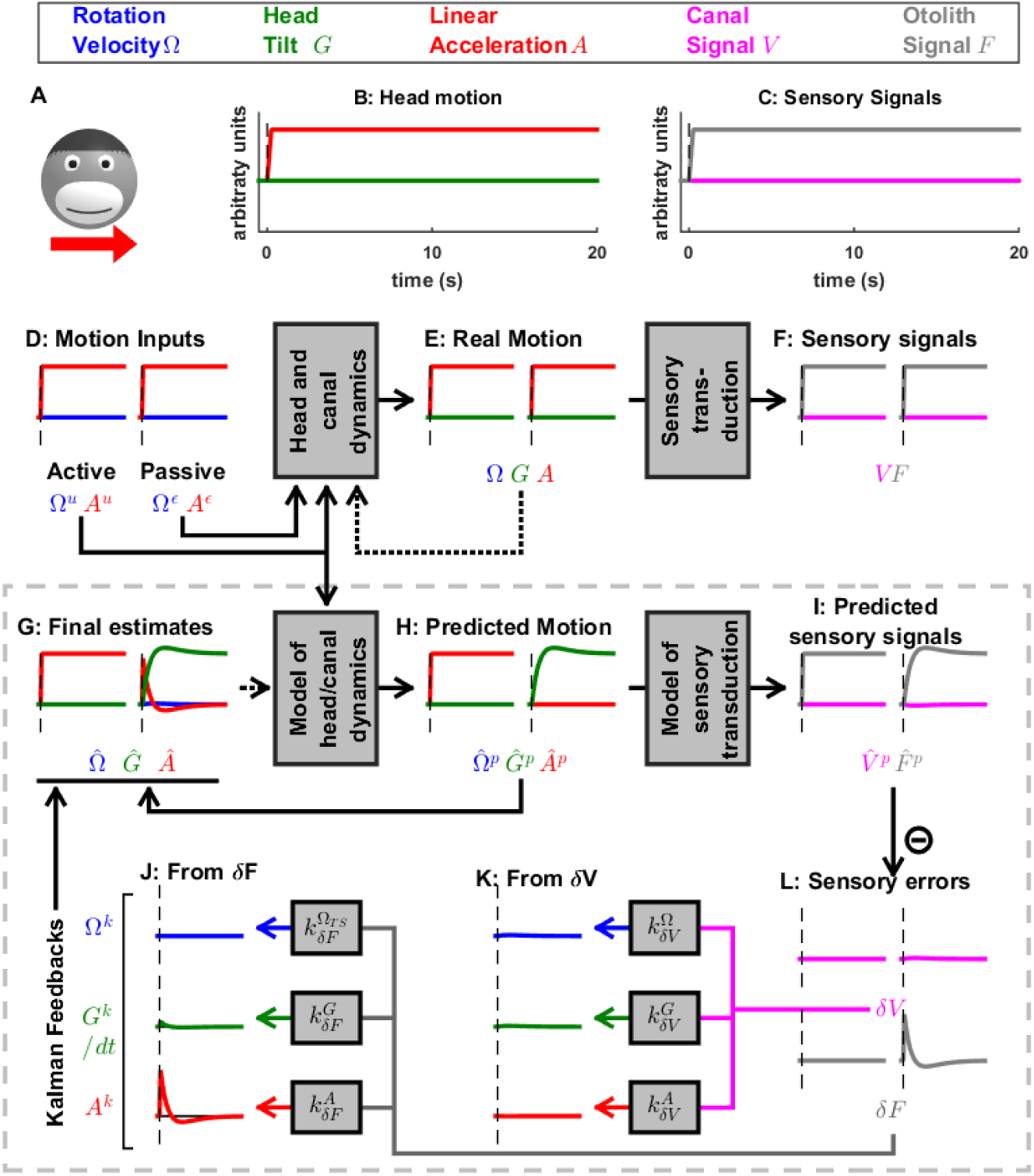
Long duration translation, demonstrating the time course of the somatogravic effect. Note that 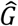 slightly overshoots *F* transiently: this is the indirect consequence of an additional feedback from *δF* to 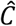 (not shown here for simplicity; but detailed in Laurens and Angelaki, 2011). In this simulation, we observe that the tilt estimate 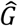 develops until its magnitude matches the otolith signal *F*. At this point, the corresponding predicted otolith signal 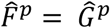 matches the real signal *F*. Therefore, the otolith error, *δF*, as well as the net acceleration estimate, 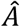, vanish. This feedback, which allows the otolith organs to create a rotation signal, has only a low magnitude and a slight effect on the dynamics of 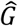 in this simulation. In general, however, it allows the otolith system to detect head rotation relative to gravity or the lack thereof and has a profound effect on the low-frequency dynamics of rotation perception (Fig. 6 Suppl. 3; see also Angelaki and Hess 1995a,b; Laurens et al. 2010; Laurens and Angelaki 2011).

**Figure 6 Supplement 2:**
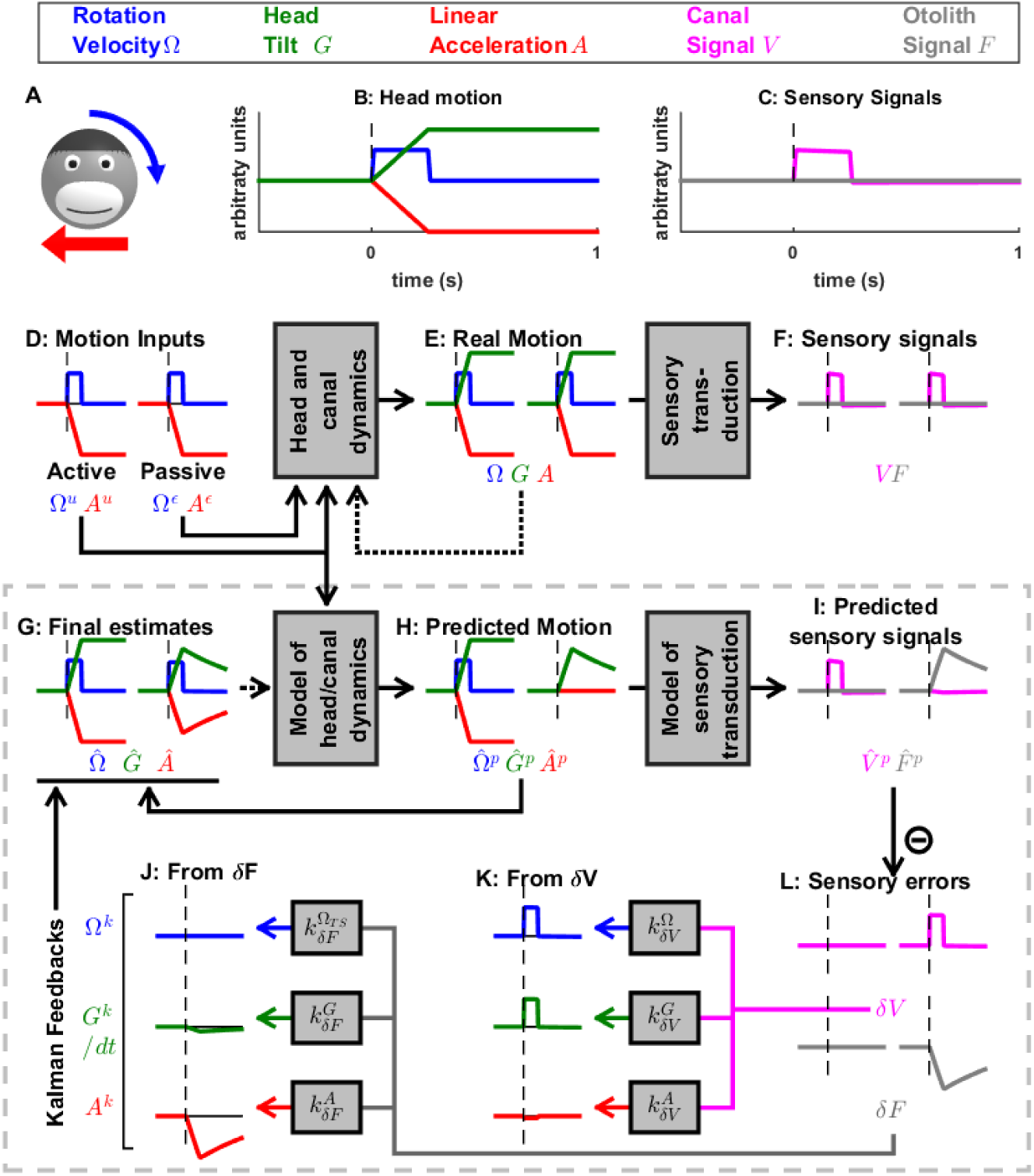
Simulation of simultaneous tilt and translation. Head tilt (as in Fig. 5) and translation (as in Fig. 6 but in the opposite direction) are performed simultaneously. As a consequence, the otolith signals (that are identical in Fig. 5C and Fig. 6C) now cancel each other (panel C, gray trace). This protocol can be seen as an activation of the semicircular canals that indicates a tilt movement, but in the absence of the corresponding tilt-induced activation of the otolith organs. As expected, and in agreement with behavioral (Angelaki et al. 1999; Merfeld et al. 1999) and neuronal findings (Angelaki et al. 2004; Shaikh et al. 2005; Yakusheva et al. 2007, 2008, 2013, Laurens et al. 2013a,b), the results of a tilt-translation simulation are the sum of a tilt simulation and a translation simulation (the latter being reversed). Interestingly, this simulation illustrates how activation of the semicircular canals without a corresponding activation of the otoliths leads to an otolith error (panel L, gray trace signaling *δF*). This simulates how translation-selective neurons in the vestibular nuclei and cerebellum modulate during tilt-translation in experimental studies using this stimulus (Angelaki et al. 2004; Shaikh et al. 2005; Yakusheva et al. 2007, 2008, 2013, Laurens et al. 2013a,b).

**Figure 6 Supplement 3:**
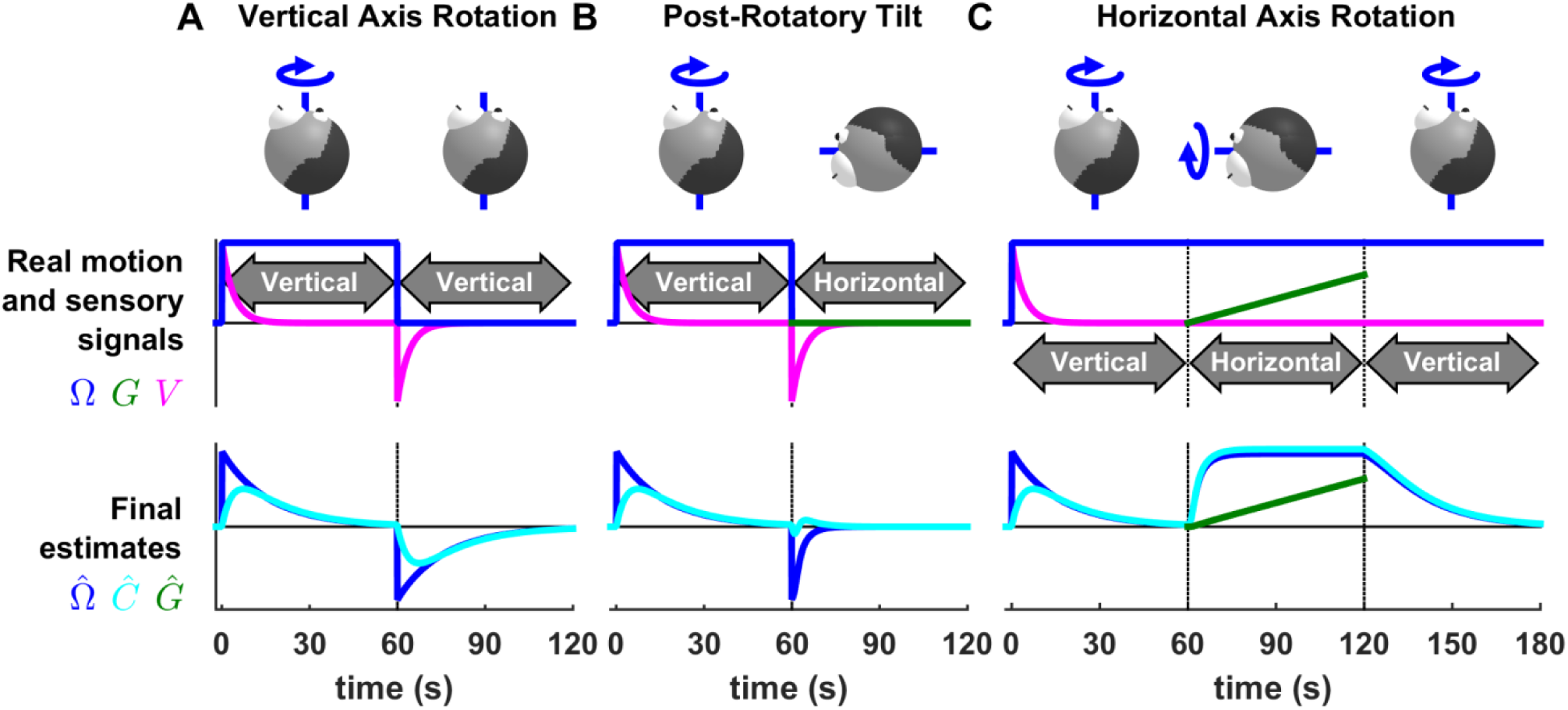
Otolith influence on rotation estimate during passive rotations. These simulations show that signals from the otoliths (that sense indirectly whether or not the head rotates relative to gravity) can also influence the rotation estimate 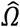 at low frequencies (this property has been extensively considered by Laurens and Angelaki, 2011). (A) Long-duration EVAR (“Vertical” refers to the orientation of the rotation axis), similar to Fig. 4 Suppl. 2B. The post-rotatory response (from t=60s to t=120s) is identical (with a reversed sign) to the initial rotatory response (from t = 0 to t = 60s). (B) Post-rotatory tilt (Angelaki and Hess 1995a, Laurens et al. 2010), where the head is rapidly reoriented at t=60s such that the rotation axis (i.e. around which the head rotated from t=0 to t=60s) becomes horizontal. The post-rotatory response is largely suppressed (time constant = 2.7s versus 16s in A). (C) Earth-horizontal Axis Rotation (Angelaki and Hess 1995b, Laurens et al. 2010), where the head is first rotated around a vertical axis until the canal signal *V* (and rotation estimates 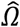 and 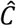) subside. Next (t=60s) the rotation axis is reoriented. The reorientation does not activate the canals (magenta). Yet, a rotation estimate 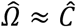 develops, rising exponentially with a short time constant (3.1s) close to the time constant of the post-rotatory response (as in Laurens et al. 2010). The rotation estimates 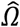 and 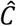 decrease exponentially when the rotation axis is reoriented back to vertical at t = 120s. These simulations illustrate the influence of the velocity feedback from the otoliths to 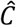 (detailed can be found in Laurens and Angelaki, 2011).

**Figure 7 Supplement 1:**
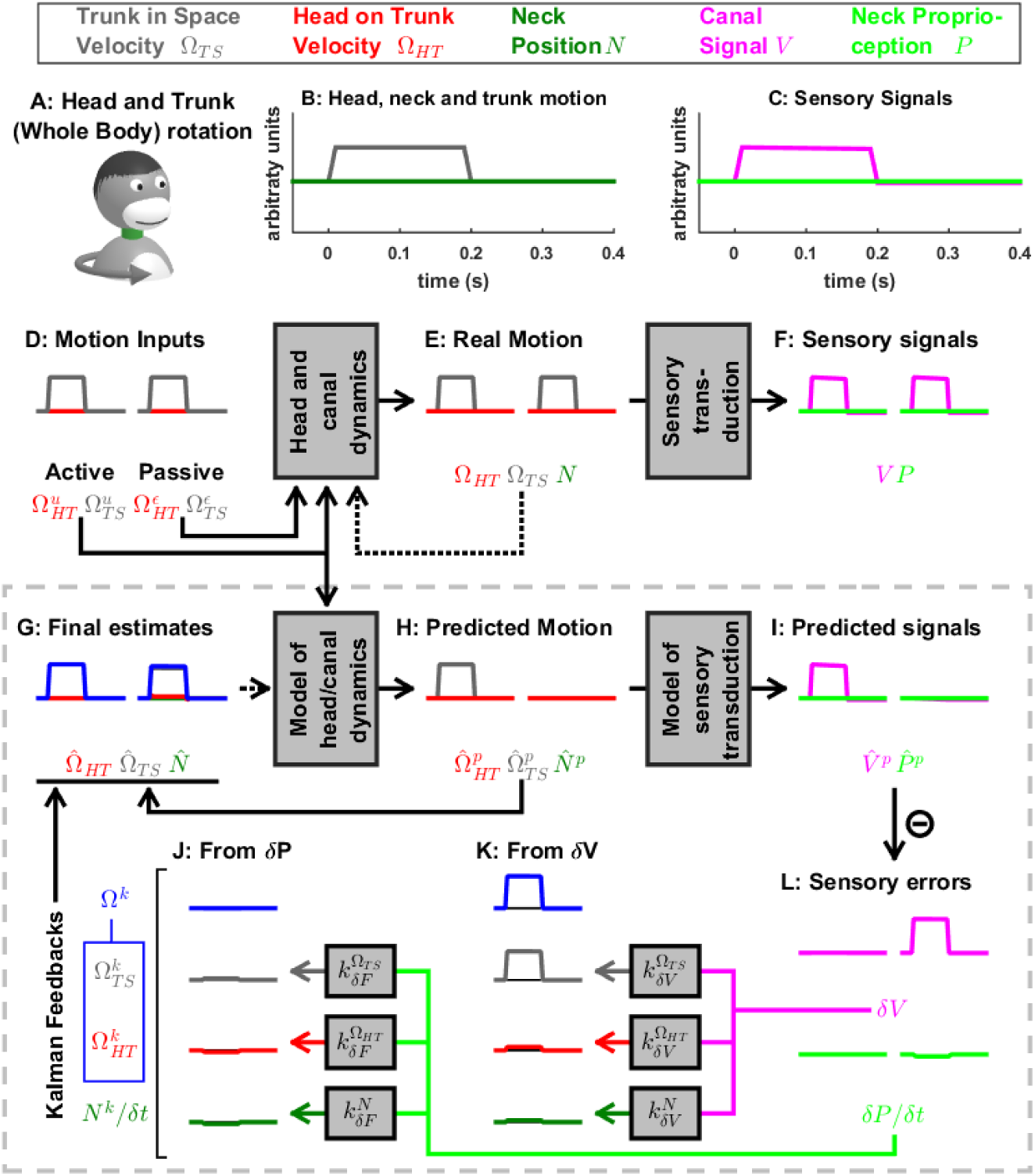
Active and passive head and trunk rotation. (A) Illustration of the stimulus. (B) Motion variables. (C) Sensory signals. (D-J) Simulated variables during active (left panels) and passive motion (right panels). Continuous arrows represent the flow of information during one time step, and broken arrows the transfer of information from one time step to the next. (K,J) Kalman feedback, shown during passive motion only (it is always zero during active movements in the absence of any perturbation and noise). For the rest of mathematical notations, see Table 1. In this simulation, the head/trunk rotates for 0.2s (thus, canal dynamics are excluded for simplicity). During active rotation, the predicted motion and sensory signals match the real motion and sensory signals, and sensory errors are null. During passive rotation, sensory errors are transformed into feedback signals (J,K) to 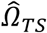 (gray), 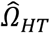 (red) and 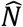 (green; note that the feedback *N*^*k*^ is scaled by 1/*δt*, see Suppl. Methods, ‘Feedback signals during neck movement’). The feedback about head velocity in space *Ω*^*k*^ (blue) is not directly computed by the model, but defined as the sum of 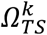 and 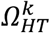. The canal error *δV* (panel L, magenta), which is similar to Fig. 3,4, is converted into a feedback about trunk in space velocity (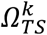, panel K, gray) and a corresponding feedback about head in space velocity *Ω*^*k*^, whereas other feedbacks originating from *δV* have a limited magnitude (see Table 3). The proprioceptive error *δP* (panel L, green; note that *δP* is scaled by 1/*δt*, see Suppl. Methods, ‘Feedback signals during neck movement’) is close to zero. In conclusion, as expected, an activation of the semicircular canals (when proprioceptive information doesn’t indicate a movement of the neck) induces a perception of whole head and trunk rotation, where the head and the trunk move together.

**Figure 7 Supplement 2:**
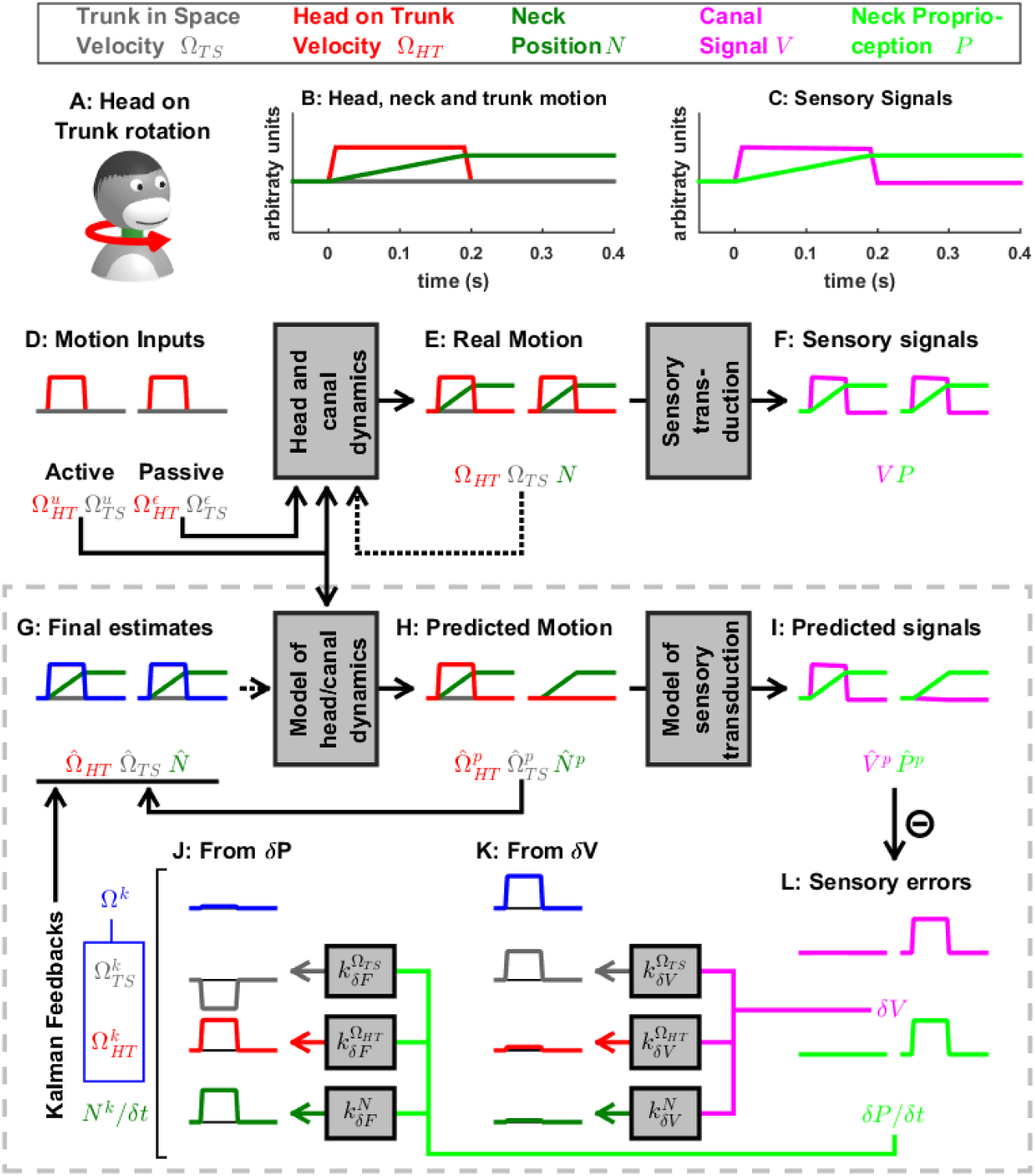
Active and passive head on trunk rotation. (A) Illustration of the stimulus. (B) Motion variables: the head rotates on the trunk (*Ω*_*HT*_, red) for 0.2s, and the neck position *N* increases during this period. (C) The sensory signals encode head velocity (canal signal, *V*, magenta) and neck position (proprioceptive signal, *P*, green). (D-J) Simulated variables during active (left panels) and passive motion (right panels). Continuous arrows represent the flow of information during one time step, and broken arrows the transfer of information from one time step to the next. (K,J) Kalman feedback, shown during passive motion only (it is always zero during active movements in the absence of any perturbation and noise).

Passive motion induces the same canal error *δV* as in Fig. 7 Suppl. 1. Furthermore, during passive motion, there is a small but significant mismatch between the predicted and actual proprioceptive signals (see Suppl. Methods, ‘Feedback signals during neck movements’) resulting in a a proprioceptive error *δP*. Note that, although proprioceptive signals *P* encode neck position, *δP* encodes the velocity of the movement (panel L; green). As in Fig. 7 Suppl. 1, the error *δV* is converted into a feedback that encodes rotation of the trunk in space (panel K, 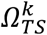, gray). However, the error *δP* is converted into an opposite feedback (panel J, 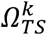, gray). Therefore, the total trunk rotation feedback and the final estimate of trunk velocity 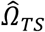 (panel G, gray) are zero. Furthermore, *δP* is converted into feedback 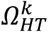 that encodes the velocity of the head relative to the trunk (panel J, red).

**Figure 7 Supplement 3:**
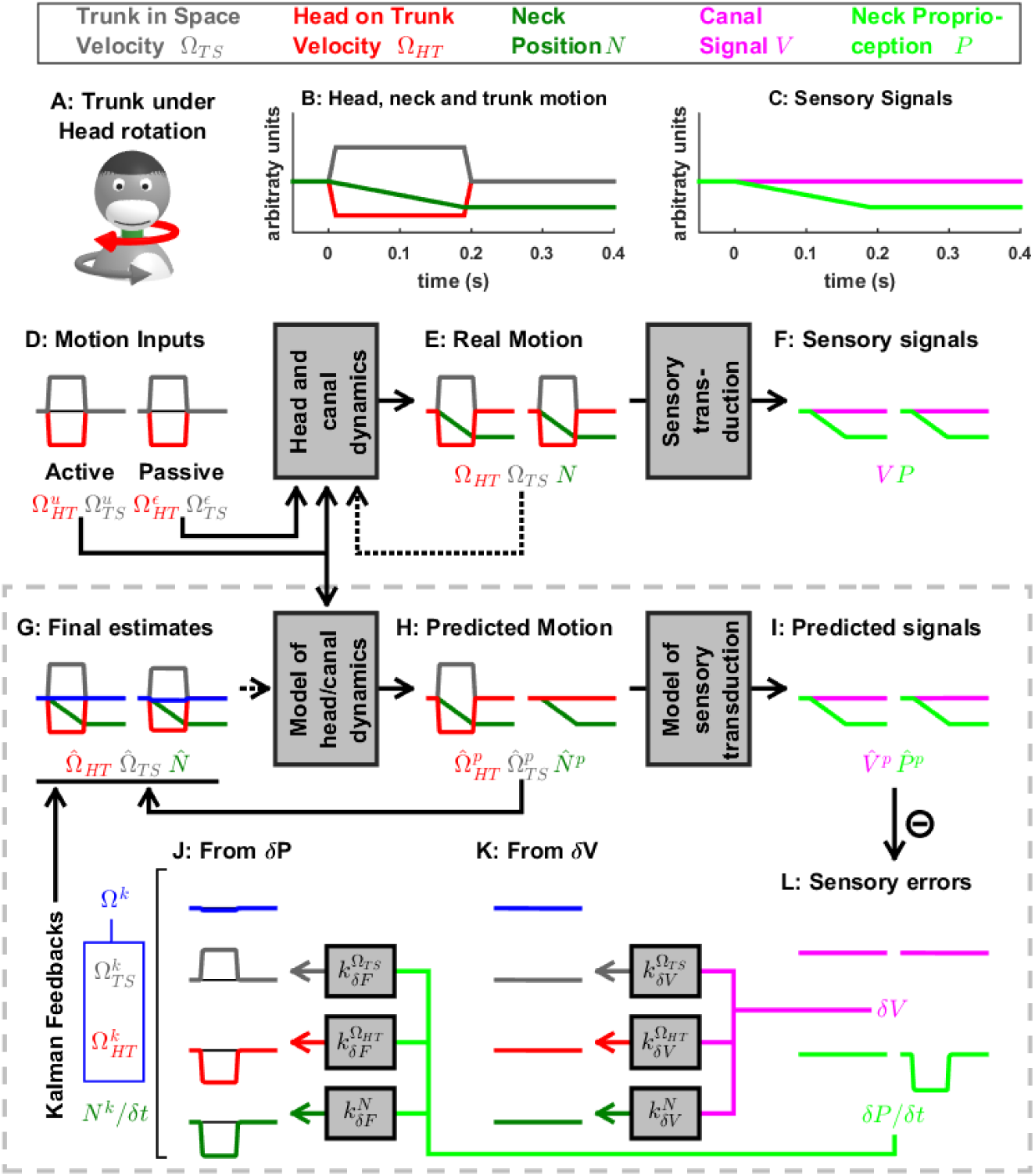
Active and passive rotation of the trunk while the head is stationary. This paradigm illustrates information processing in response to neck rotation in the absence of semicircular canal stimulation. Signals and parameters as in Fig. 7 Suppl. 1 and 2.

**Figure 7 Supplement 4:**
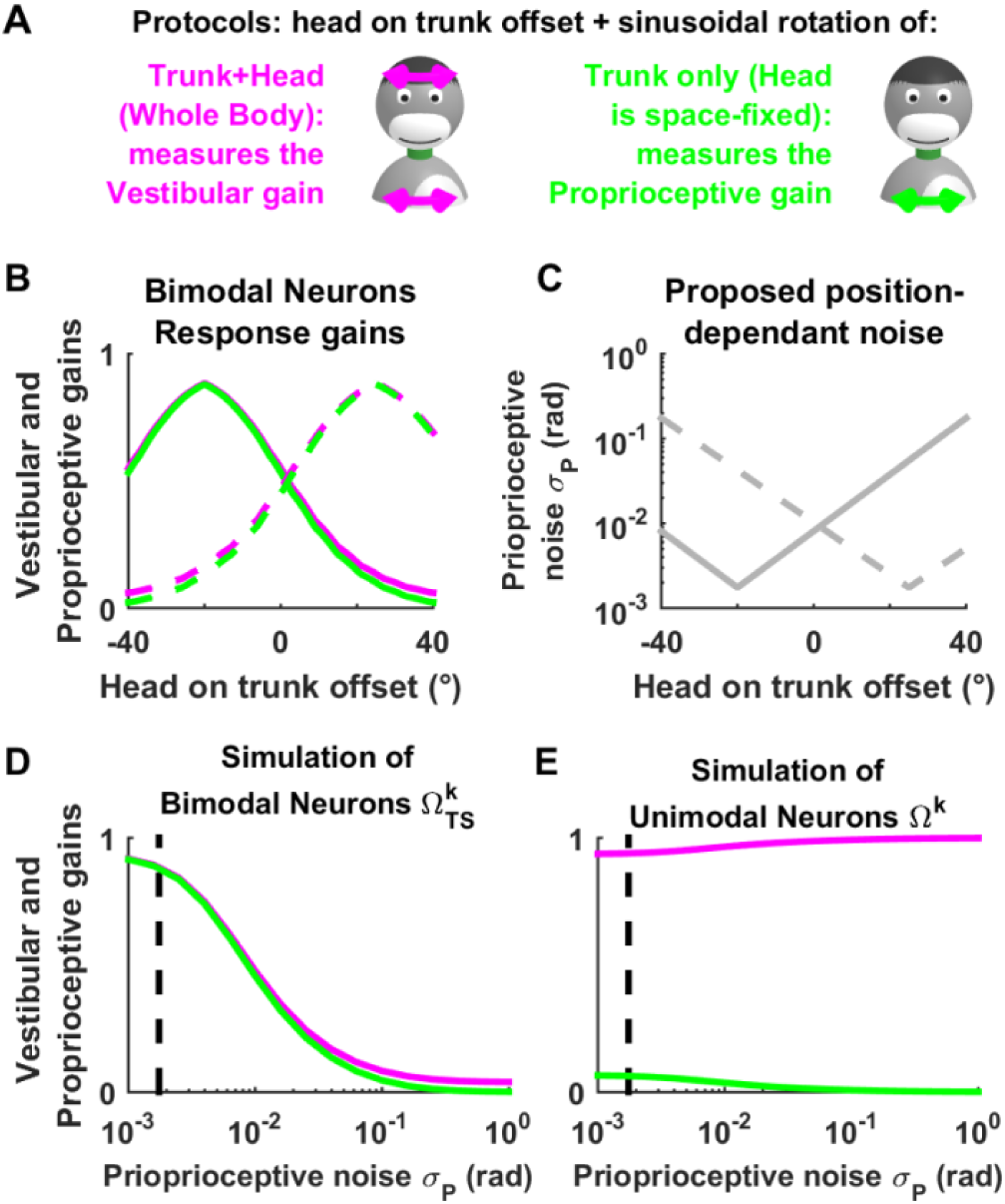
Simulated tuning of unimodal and bimodal neuron as a function of neck position offset. (A) Experimental manipulation used by (Brooks and Cullen 2009) to demonstrate that the response of bimodal neurons to sinusoidal motion varies if a head rotation offset is superimposed to the sinusoidal stimulus. (B) Schematic responses of two bimodal neurons (one in solid lines and the other in broken lines). Individual neurons exhibit hill-shaped tuning curves as a function of head rotation offset, and the location of the peak varies from neuron to neuron. Remarkably, neurons exhibit similar tuning curves in response to canal stimulation (rotation of the head and trunk; magenta curves), and to neck proprioceptor stimulation (rotation of the trunk only, green). The fact that these tuning curves are matched allows bimodal neurons to encode trunk velocity during combined trunk and head motion, regardless of head position offset (Brooks and Cullen 2009). However, the reason why bimodal neuron responses are modulated as a function of head offset is unknown.

We propose that head rotation offset extends or contracts individual neck muscles and affects the signal to noise ratio of their afferent proprioceptive signals. Assuming that individual neurons receive proprioceptive afferents from distinct pools of muscles, the signal to noise ratio of proprioceptive signals would form a distinct curve in each neuron (C: the solid and broken curves correspond to the solid and broken curves in B). (D) Simulations of bimodal neurons (i.e. the trunk velocity feedback 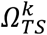) during passive trunk and head rotation (magenta) and trunk only rotation (green), with increasing amount of proprioceptive noise *σ*_*P*_ (the vertical broken lines in (A,B) indicate the value *σ*_*P*_ used in other figures). We find that increasing *σ*_*P*_ leads to a concomitant decrease of the response gains in both conditions. The responses curves in (B) can be obtained by combining panels (C) and (D).

(E) Furthermore, we find that the simulated response of unimodal neurons (assumed to encode the trunk velocity feedback *Ω*^*k*^) is independent of proprioceptive noise, in agreement with experimental results (Brooks and Cullen 2009) that demonstrate that that gain of unimodal neurons is not sensitive to static neck position.

**Figure 7 Supplement 5:**
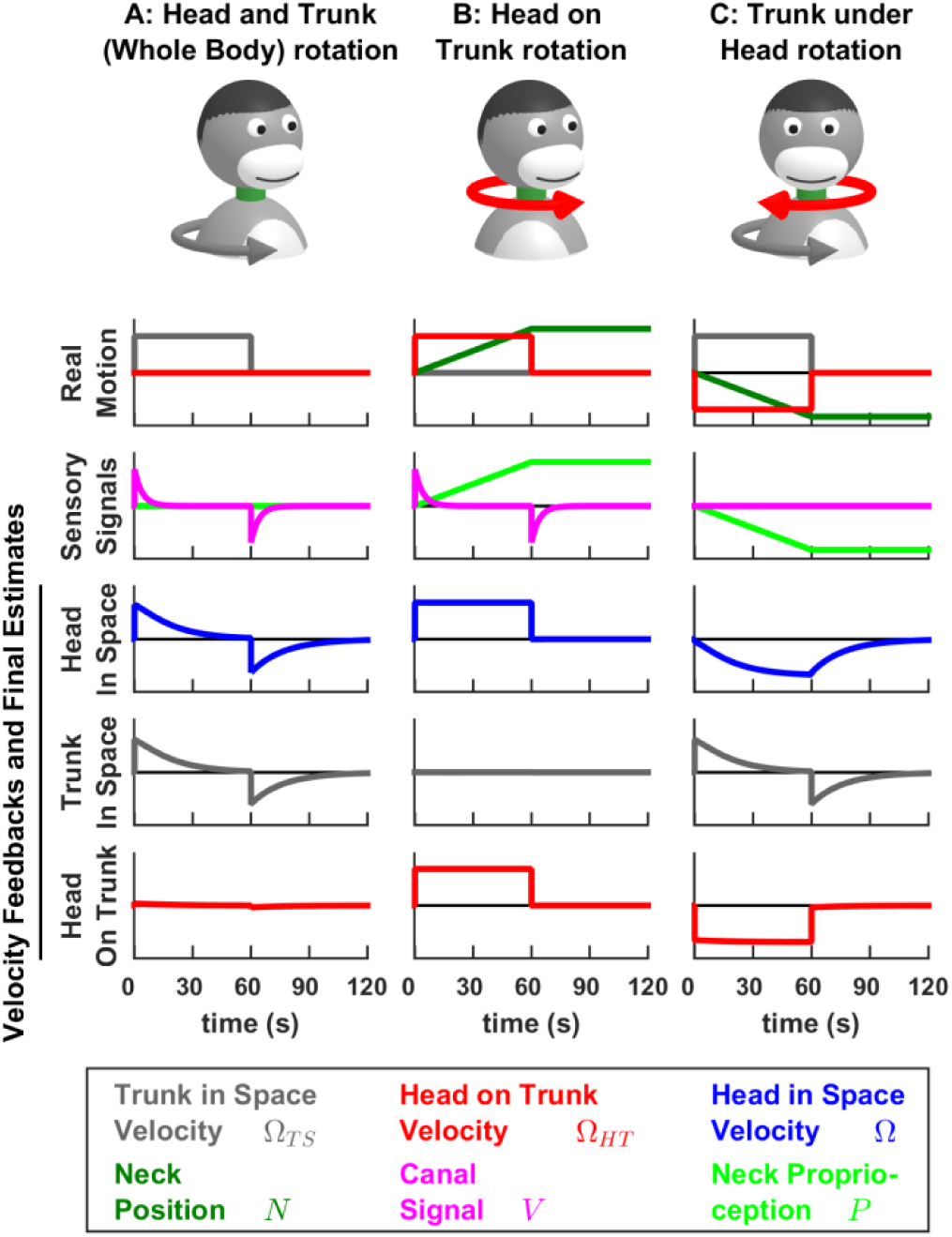
Long-duration passive head and trunk movements. The dynamics of head and trunk rotation perception during combined passive head and trunk movements was studied by (Mergner et al. 1991). Here we reproduce the results of this study by simulating the same movements as in Fig. 7 with a longer rotation duration (60s). The feedback signals about head in space velocity (third row, blue), trunk in space velocity (fourth row, gray) and head on trunk velocity (fifth row, red) are shown. During passive motion, these feedback are identical to the final motion estimates, and therefore, the curves in rows 3 to 5 also represent the perception of motion during these stimuli.

A combined rotation of the head and the trunk (A) activates the canals (magenta curve). Following the initial acceleration at t=0, canal signals fade away with a time constant of 4s. A canal aftereffect occurs following the deceleration at t=60s. The estimates of head in space and trunk in space velocity are identical to each other, and exhibit similar dynamics as the canal signal, although with a longer time constant of 16.5s (for the same reasons as those described in Fig. 4 Suppl. 1B).

When rotating the head while the trunk is stationnary (B), the simulated estimates of head in space (third row, blue) and head on trunk (fifth row, red) velocity persist indefinitely, as illustrated experimentally by Mergner et al. (1991). The difference in the time course of the signals in panels A and B is due to the additional presence of the neck proprioceptive signal.

Strinkingly, during a long duration rotation (C), the estimate of trunk in space velocity (fourth row, gray) decreases with an identical time constant as in (A), although the underlying sensory signals, that originate from the canals in A and from neck proprioception in C, have fundamentally different dynamics. Futhermore, the perception of trunk rotation is replaced by an illusion of head rotation in space (third row, blue) in an opposite direction. These model predictions exactly match experimental findings by Mergner et al. (1991).

**Figure 7 Supplement 6:**
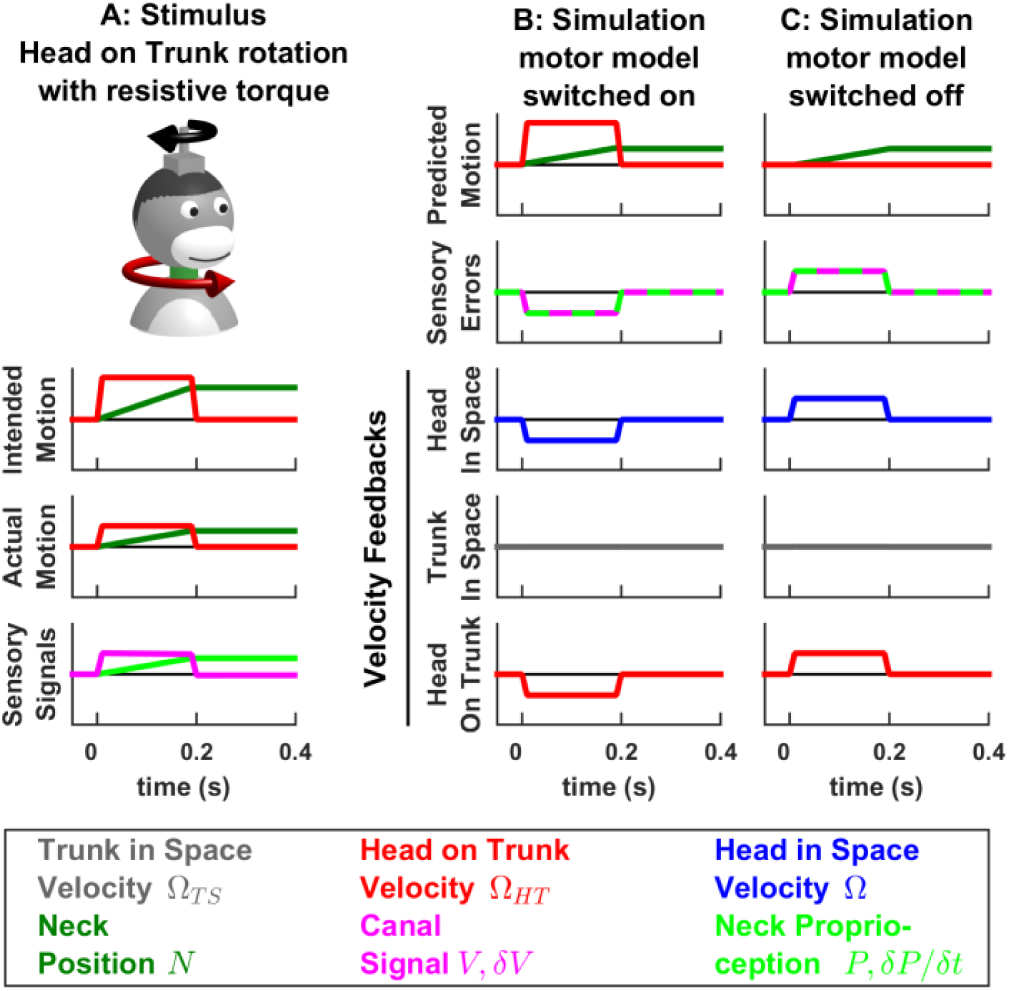
Impact of perturbing motor activity during active movement. Experimental studies (Roy and Cullen, 2001; 2004; Carriot et al. 2013; Brooks et al., 2015; Brooks and Cullen, 2015; Cullen and Brooks, 2015) have demonstrated that central neurons encode active and passive motion (i.e. head velocity or linear acceleration) indiscriminately when active movements are blocked or perturbed by unexpected torque, as simulated here. (A) Simulated experimental condition, where the animal attempts to perform a head movement (while the trunk remains immobile). Due to an unexpected resistive torque, the actual movement amplitude and velocity are only half of the intended motion, as indicated by neck position (green) and head velocity (red) traces (compare “Intended Motion” and “Actual Motion” panels). The resulting sensory signals (canals, magenta, and neck proprioceptors, green) encode the actual (reduced) velocity and amplitude of the movement. (B) Kalman filter simulation using the same model than in Fig. 7. Because the actual head velocity (A, “Actual Motion”) is slower than the predicted velocity (B, “Predicted Motion”), a negative canal error *δV* occurs (the predicted neck position and proprioceptive error are discussed below). As a result, the velocity feedback about head in space (blue) is negative. (C) Alternative simulation, where the internal model of the motor plant is switched off (the matrix M in Fig. 1 is set to zero). In this situation, the motion is perceived as if it was entirely passive (exactly as in Fig. 7 Suppl. 2, with only half the amplitude) and therefore the feedback pathways encode a positive head in space velocity signal. Thus, the simulation in B predicts that central neurons, that encode feedback signals about head velocity in space, are inhibited in this experimental condition. In contrast, the simulation in C predicts that these neurons are activated (and encode net head velocity with the same gain as in Fig. 7 Suppl. 2). Experiments (Brooks et al. 2015) yield results consistent with (C) but contrary to (B). This indicates that a mechanism, which is additional to the Kalman filter algorithm (shown Fig. 1, 9 and used in panel B), switches the motor model off when active movements are perturbed.

Note that, in (B) and (C), the predicted neck position is close to the actual position and the proprioceptive error *δP*/*δt* encodes a velocity error (close to *δV*). This occurs for the same reason as during passive neck movements, as described in Supplementary Methods, “Feedback signals during neck movement”.

**Figure 9 Supplement 1:**
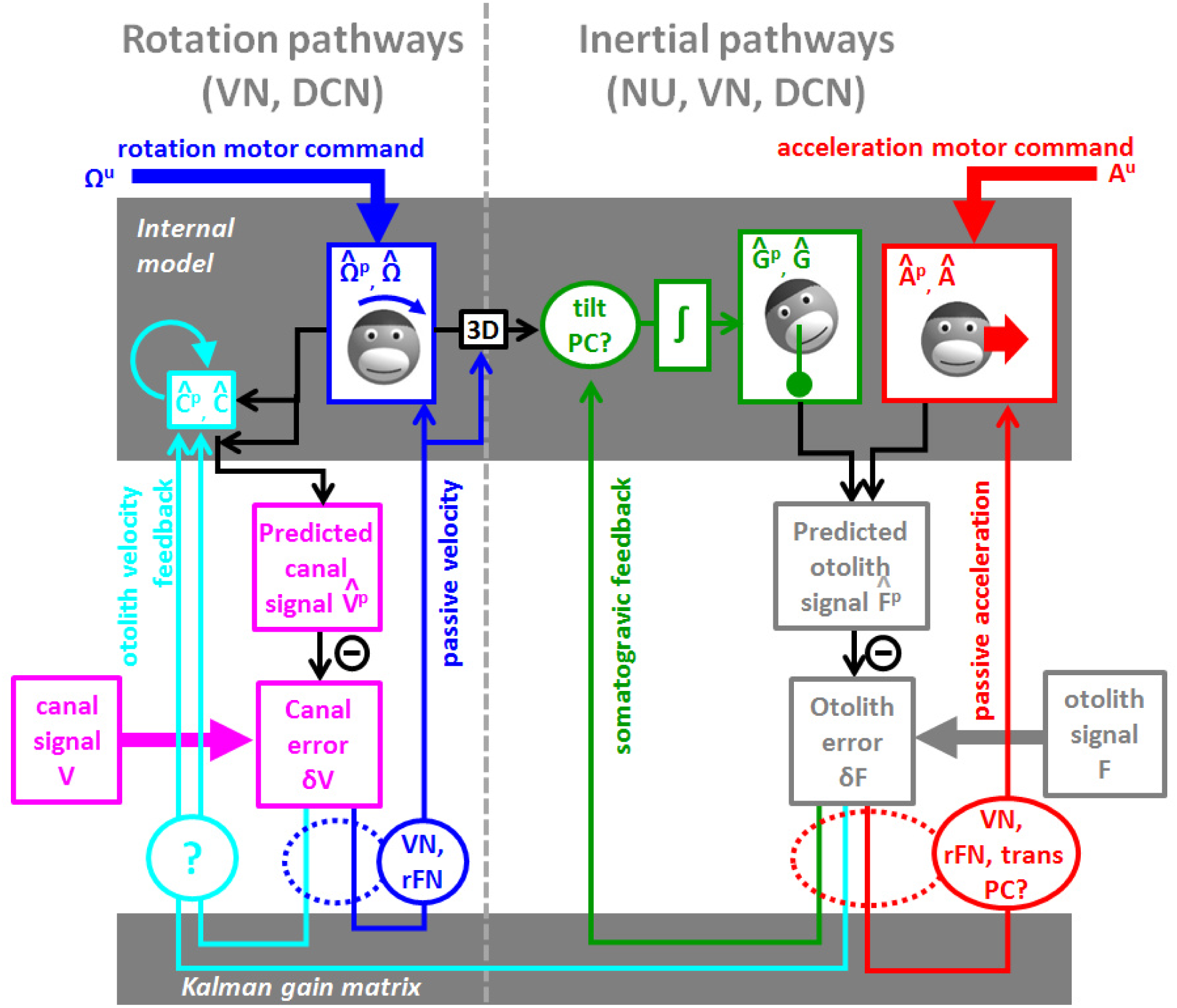
Alternative diagram, where the Kalman feedback encoding passive tilt velocity is removed. Instead, a final rotation estimate 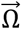 is computed and transformed into a tilt velocity signal.

**Figure 9 Supplement 2:**
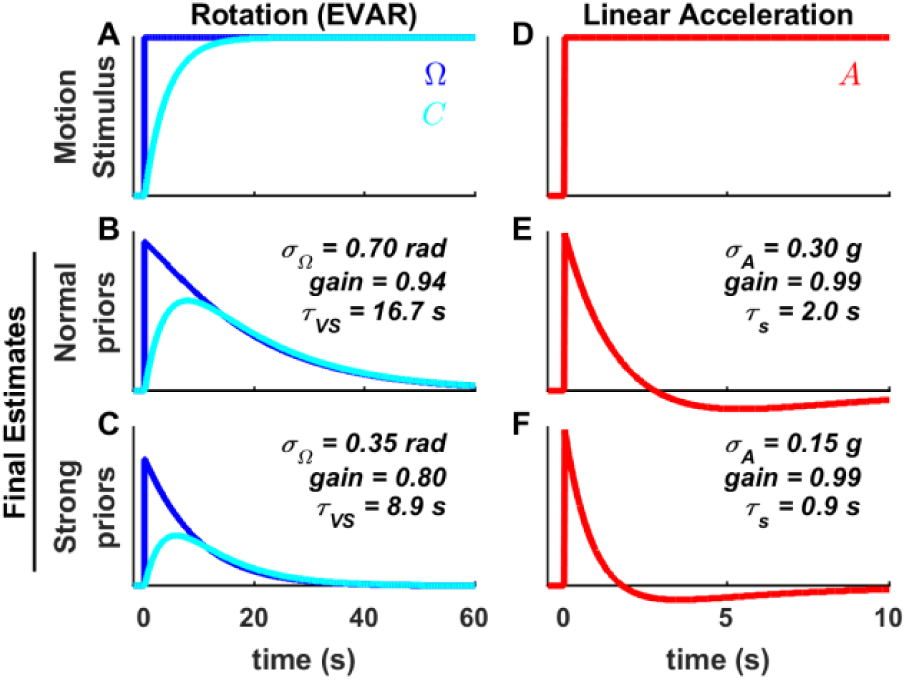
Influence of Bayesian priors on the dynamics of motion estimates. We simulated a passive long-duration rotation (A-C, similar as Fig. 4 Suppl. 2A) and a passive constant linear acceleration (D-E, similar as Fig 6 Suppl. 1). Simulations were performed with the same set of parameters as in other figures (B,E), or with lower values of *σ*_Ω_. and *σ*_A_ resulting in narrower distributions of *Ω*^*ε*^ (C) and *A*^*ε*^ (F). These distributions represent Bayesian priors that drive dynamic motion estimate towards zero. As a consequence, narrowing the priors decreased the gain and time constant of the rotation estimate (C; time constant *τ*_VS_) and of the acceleration estimate (D; time constant *τ*_*s*_).

## Supplementary Methods

Here we describe the Kalman model in more detail. We present the model of head motion and vestibular information processing, first as a set of linear equations (‘Model of head motion and vestibular sensors’), and then in matrix form (‘Model of head motion in matrix form’). Next we present the Kalman filter algorithm, in the form of matrix computations (‘Kalman filter algorithm’) and then as a series of equations (‘Kalman filter algorithm developed’).

Next, we derive a series of properties of the internal model computations (‘Velocity Storage during EVAR’; ‘Passive Tilt’,’ Kalman feedback gains’,’ Time constant of the somatogravic effect’, ‘Model of motor commands’).

We then present some variations of the Kalman model (‘Visual rotation signals’, ‘Model of head and neck rotations’, ‘Feedback signals during neck movement’, ‘Three-dimensional Kalman filter’).

### Model of head motion and vestibular sensors

The model of head motion in (Fig. 2) can be described by the following equations (see Table 1 for a list of mathematical variables):

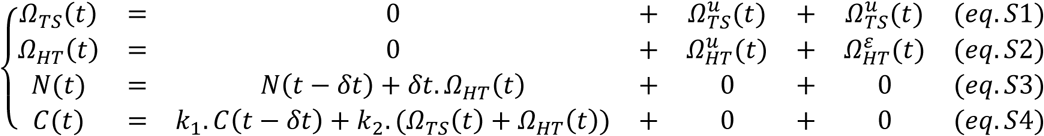

Here, *eq*.1 states that head velocity *Ω*(*t*) is the sum of self-generated rotation *Ω*^*u*^(*t*) and an externally generated rotation *Ω*^*ε*^(*t*). Therefore, in the absence of motor commands, *Ω*(*t*) is expected to be zero on average, independently from all previous events.

*Eq*.2 describes the first-order low-pass dynamics of the canals:

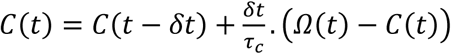

which yields:

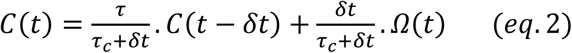

with 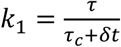 and 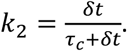.

*Eq*.3 integrates rotation *Ω* into tilt *G*. The variable *s* acts as a switch: it is set to 1 during tilt and to 0 during EVAR (in which case *G* remains equal to zero, independently of *Ω*).

Finally, *Eq*.4 that describes linear acceleration, resembles *Eq*.1 in form and properties.

The system of these equations is rewritten as follows in order to eliminate *Ω*(*t*) from the right-hand side (which is needed so that it may fit into the form of *eq*.7 below):

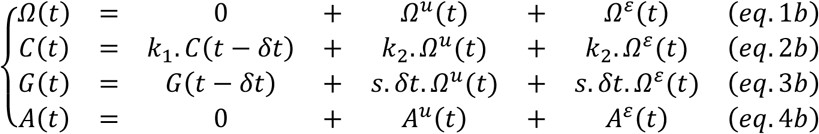

The model sensory transduction is:

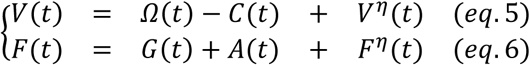

*Eq*.5 indicates that the semicircular canals encode rotation velocity, minus the dynamic component *C*; and *Eq*.6 indicates that the otolith organs encode the sum of tilt and acceleration.

### Model of head motion in matrix form

The system of equations (1*b* − 4*b*) can be rewritten in matrix form:

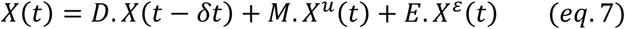

with:

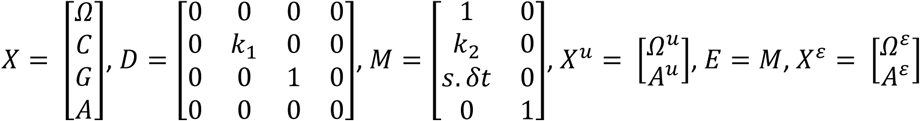

Similarly, the model of sensory transduction (*eq*.5 − 6) is rewritten as:

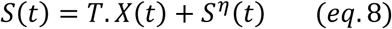

With:

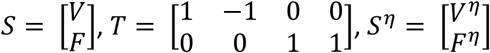

Given the standard deviations of *Ω*^*ε*^, *A*^*ε*^, *V*^*η*^ and *F*^*η*^ (*σ*_*Ω*_, *σ*_*A*_, *σ*_*V*_ and *σ*_*F*_), the covariance matrices of *X*^*ε*^ and *S*^*η*^ (that are needed to perform Kalman filter computations) are respectively:

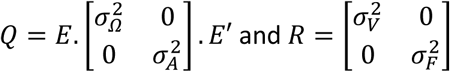

### Kalman filter algorithm

The Kalman filter algorithm (Kalman 1960) computes optimal state estimates in any model that follows the structure of (*eqn*. 7,8) (Fig. 1). The optimal estimate 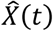 is computed by the following steps (Fig. 1B):

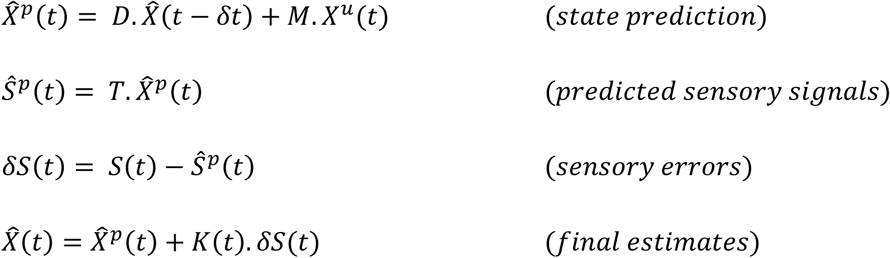

The Kalman gain matrix *K*(*t*) is computed as:

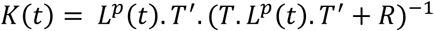

where *L*^*p*^(*t*) = *D*. *L*(*t − δt*). *D*′ + *Q* and *L*(*t*) = (*Id − K*(*t*). *T*). *L*^*p*^(*t*) are the covariance of the predicted and updated estimates, *Q* and *R* are the covariance matrices of *E*. *X*^*ε*^ and *S*^*η*^, and *Id* is an identity matrix. These equations are not shown in Fig. 1.

The initial conditions of 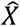 are set according to the initial head position in the simulated motion, and the initial value of *L* is *L* = *Q*.

### Kalman filter algorithm developed

The inference is performed by applying the Kalman filter algorithm on *Eqs*. 7 - 8. The corresponding computations can be expanded in the following equations:

*State predictions:*

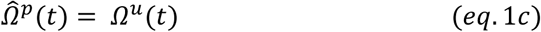

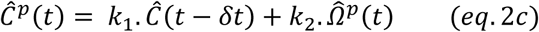

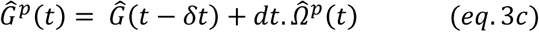

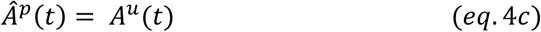

*Sensory predictions:*

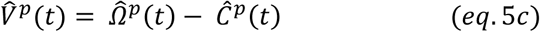

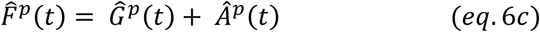

*Sensory errors:*

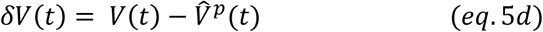

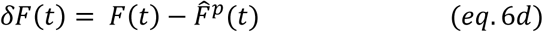

*Final estimates:*

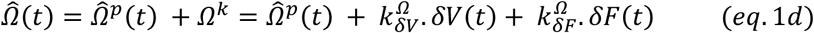

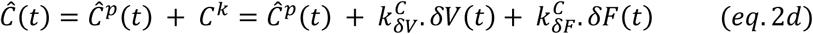

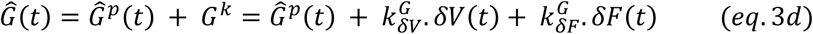

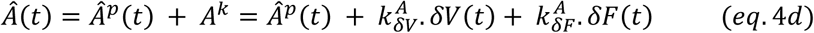

These equations form the basis of the model (in Fig. 9, 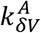 and 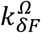 are assumed to be zero, see ‘Kalman feedback gains’ and Table 2).

### Velocity Storage during EVAR

Here we analyze the Kalman filter equations to derive the dynamics of rotation perception during passive EVAR and compare it to existing models. During passive EVAR (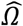^*p*^ = *Ω*^*u*^ = 0 and *δF* = 0), the dynamics of the rotation estimate depends of 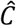, which is governed by (*eq*. 2*c*, *d*):

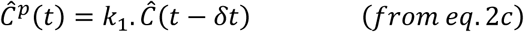

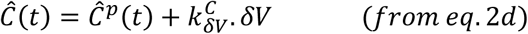

With:

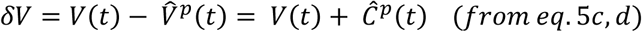

Based on these equations, 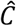 follows a first-order differential equation:

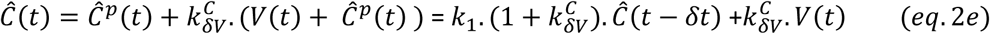

This equation is characteristic of a leaky integrator, that integrates *V* with a gain 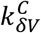, and has a time constant *τ*_*VS*_ which is computed by solving:

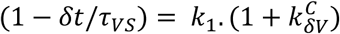

Based on the values of Table 2, we compute *τ*_*VS*_=16.5s (in agreement with the simulations in Fig. 4 Suppl. 2).

The final rotation estimate is the sum of 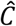 and the canal signal:

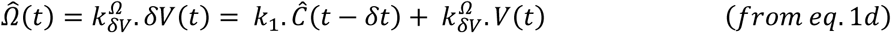

These equations reproduce the standard model of (Raphan et al. 1979). Note that the gains, 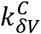 = 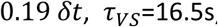, and 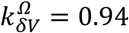 are similar to the values in (Raphan et al. 1979) and to model fits to experimental data in (Laurens et al. 2011b). The dynamics of 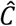 (*eq*. 2*e*) can be observed in simulations, i.e. in Fig. 4 Suppl 2B where the leaky integrator is charged by vestibular signals *V* at t = 0 to 10s and t = 60 to 70s; and subsequently discharges with a time constant *τ*_*VS*_=16.5s. The discharge of the integrator is also observed in Fig. 4 Suppl 2C when t>60s Fig. 6 Suppl 3C when t>120s.

### Passive tilt

Here we provide additional mathematical analyses about motion estimation during passive tilt. During passive tilt 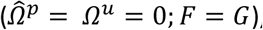, the internal estimate 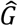 follows:

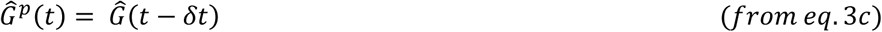

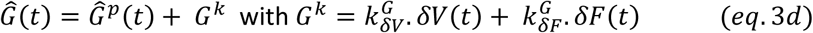

First, we note that (*eq*. 3*c*, *d*) combine into 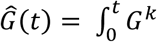. In other words, the tilt estimate during passive tilt is computed by integrating feedback signals *G*^*k*^.

Also, to a first approximation, the gain 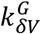 is close to *δt*, the canal error *δV* encodes *D* and *δF* is approximately null. In this case, *G*^*k*^ ≈ *δt*. *Ω* and 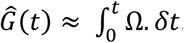. Therefore, during passive tilt (Fig. 5, Fig. 5 Suppl. 1), the internal model (*eq*. 3*c*) integrates tilt velocity signals that originate from the canals and are conveyed by feedback pathways.

Note, however, that the Kalman gain 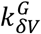 is slightly lower than *δt* (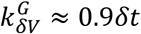; Table 2). Yet, the final tilt estimate remains accurate due to an additional feedback originating from *δF* which can be analyzed as follows. Because 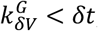, the tilt estimate 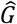 lags behind *G*, resulting in a small otolith error 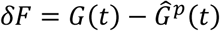 that contributes to the feedback signal (via the term 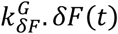 in *eq*. 3*d*). The value of *δF* stabilizes to a steady-state where 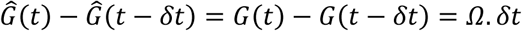. Based on *eq*. 3*d*, we obtain:

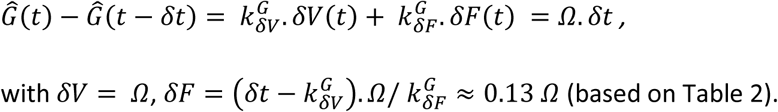

Thus, a feedback signal originating from the otolith error complements the canal error. This effect is nevertheless too small to be appreciated in Fig. 5.

### Kalman feedback gains

Here we provide additional information about Kalman gains and we justify that some feedback signals are considered negligible.

First, we note that some values of the Kalman gain matrix (those involved in a temporal integration), include the parameter *δt* (Table 2). This is readily explained by the following example. Consider, for instance, the gain of the vestibular feedback to the tilt estimate 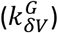. During passive tilt, the tilt estimate 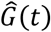 is tracked by the Kalman filter according to:

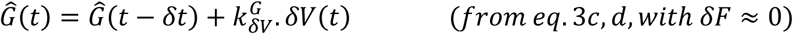

Since 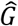 is computed by integrating canal signals (*δV*) over time, we would expect the equation above to be approximately equal to the following:

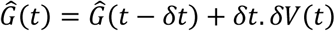

Therefore, we expect that 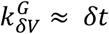. When simulations are performed with *δt* = 0.01s, we find indeed that 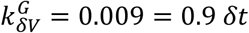. Furthermore, if simulations are performed again, but with *t* = 0.1s, we find 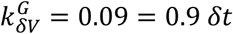. In other words, the Kalman gain computed by the filter is scaled as a function of *δt* in order to perform the operation of temporal integration (albeit with a gain of 0.9). For this reason, we write 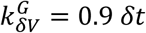 in Table 2, which is more informative than 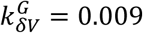. Similarly, other Kalman gain values corresponding to 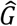 and 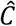 (which is also computed by temporal integration of Kalman feedbacks) are shown as a function of *δt* in Table 2.

Furthermore, the values of *C*^*k*^ and *G*^*k*^ are divided by *δt* in all figures, for the same reason. If, for example, *δV* = 1, then the corresponding value and *G*^*k*^ would be 0.009 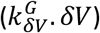. This value is correct (since 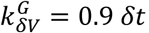) but would cause *G*^*k*^ to appear disproportionately small. In order to compensate for this, we plot *G*^*k*^/*dt* in the figures. The feedback *C*^*k*^ is scaled in a similar manner. In contrast, neither *Ω*^*k*^ nor *A*^*k*^ are scaled.

Note that the feedback gain 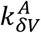 (from the canal error *δV* to 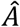) is equal to 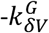 (Table 2). This compensates for a part of the error *δF* during tilt (see previous section), which generates an erroneous acceleration feedback *A*^*k*^. This component has a negligible magnitude and is not discussed in the text or included in the model of Fig. 9.

Note also that the Kalman filter gain 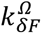 is practically equal to zero (Table 2). In practice, this means that the otoliths affect rotation perception only through variable 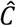. Accordingly, otolith-generated rotation signals (e.g. Fig. 6 Suppl. 3C, from t = 60s to t = 120s) exhibit low-pass dynamics.

Because 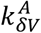 and 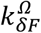 are practically null and have no measurable effect on behavioral or neuronal responses, the corresponding feedback pathways are excluded from Fig. 9.

### Time constant of the somatogravic effect

Here we analyze the dynamics of the somatogravic effect. During passive linear acceleration, the otolith error *δF* determines the feedback *G*^*k*^ that aligns 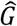 with *F* and therefore minimizes the feedback. This process can be modeled as a low-pass filter based on the following equations:

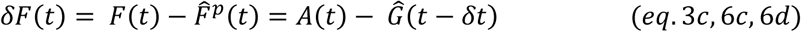

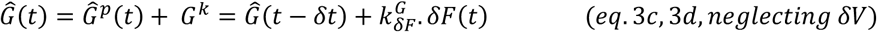

Leading to:

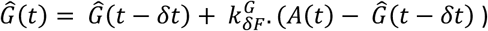

This equation illustrates that 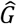 is a low-pass filter that converges towards *A* with a time constant *τ*_*s*_ = 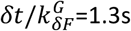 (Table 2).

Note that the feedback from *δF* to 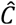 adds, indirectly, a second component to the differential equation above, leading 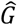 to transiently overshoot *A* in Fig. 6 Suppl. 1. Nonetheless, describing the somatogravic effect as a first-order low-pass filter is accurate enough for practical purposes.

### Model of motor commands

In this model, we have assumed that motor commands simply encode head rotation velocity and linear acceleration. However, more complex models could be designed where motor commands affect other movement parameters. These models would be encoded by changing the matrices *M* and *X*^*u*^.

However, neither *M* nor *X*^*u*^ appear in the Kalman filter algorithm outside of (*eq*. 7). Therefore, the only role of the motor command is to contribute to the prediction (*eq*. 7). Importantly, motor commands *X*^*u*^ don’t carry any noise (motor noise is a part of *X*^*ε*^); and neither *M* nor *X*^*u*^ participate in the computing of the Kalman gain matrix *K*. Therefore, changing the model of motor commands would have no effect on Kalman feedbacks.

Furthermore, in all simulations presented in this study, we have observed that motor commands were transformed into 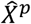 that encoded active movement accurately. We reason that, if the model of motor commands was changed, it would still lead to accurate predictions of the self-generated motion component, as long as the motor inputs are unbiased and are sufficient to compute all motion variables, either directly or indirectly through the internal model. Under these hypotheses, simulation results would remain unchanged. This justifies our approach of using the simplest possible model of motor commands.

### Visual rotation signals

In Fig. 4 Suppl. 2C, a visual sensory signal *Vis* was added to the model as in (Laurens 2006, Laurens and Droulez 2008) by simply assuming that it encodes rotation velocity:

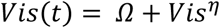

Where *Vis*^*η*^ is a Gaussian noise with standard deviation *σ*_*Vis*_ = 0.12 rad/s (Laurens and Droulez 2008). This signal is incorporated into the matrices of the sensory model as follows:

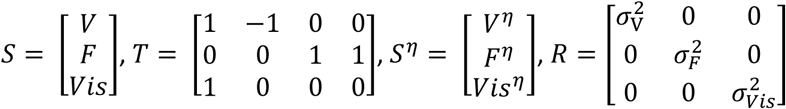

The model of head motion and the matrix equations of the Kalman filter remain unchanged.

### Model of head and neck rotations

We created a variant of the Kalman filter, where trunk velocity in space and head velocity relative to the trunk are two independent variables *Ω*_*TS*_ and *Ω*_*HT*_. We assumed that head position relative to the trunk is sensed by neck proprioceptors. To model this, we added an additional variable *N* (for ‘neck’) that encodes the position of the head relative to the trunk: *N* = *∫ Ω*_*HT*_. *dt*. We also added a sensory modality *P* that represents neck proprioception.

Total head velocity (which is not an explicit variable in the model but may be computed as *Ω* = *Ω*_*TS*_ + *Ω*_*HT*_) is sensed through the semicircular canals, which were modeled as previously.

The model of head and trunk motion is based on the following equations:

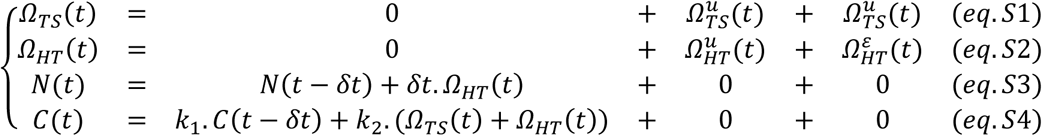

Note that (*eq*. *S*1) and (*eq*. *S*2), are analogous to (*eq*. 1) in the main model and imply that *Ω*_*TS*_ and *Ω*_*HT*_ are the sum of self-generated 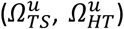 and unpredictable components 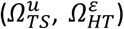. E*q*. *S*3 encodes *N* = *∫ Ω*_*HT*_. *dt*. The canal model (*eq*. *S*4) is identical as in (*eq*. 2), the input being the velocity of the head in space, i.e. *Ω*_*TS*_(*t*) + *Ω*_*HT*_(*t*).

The sensory model includes the canal signal *V* and a neck proprioceptive signal *P* that encodes neck position:

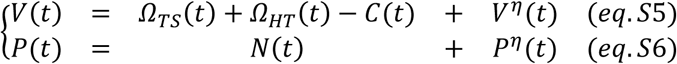

Note that (*eq*. *S*5) is identical to (*eq*. 5) in the main model, and that *P* is subject to sensory noise *P*^*𝜂*^.

Similar to the main model, (*eq*. *S*1 *− S*6) are written in matrix form:

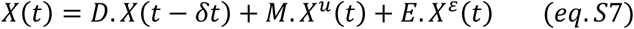

with:

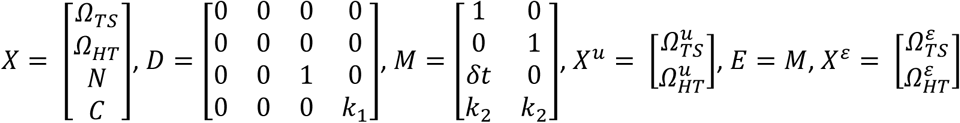

Similarly, the model of sensory transduction (*eq*. *S*5 *− S*6) is rewritten as:

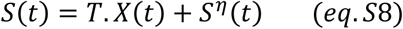

with:

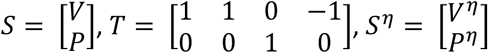

Given the standard deviations of 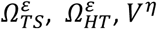 and 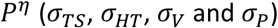, the covariance matrices of *X*^*ε*^ and *S*^*𝜂*^ are respectively:

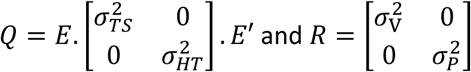

Simulations were performed using the Kalman filter algorithm, as in the main model.

### Feedback signals during neck movement

In Fig. 7 Suppl. 2,3, we note that passive neck motion induces a proprioceptive feedback *δP* that encodes neck velocity, although proprioceptive signals *P* are assumed to encode neck position. This dynamic transformation is explained by considering that, during passive motion:

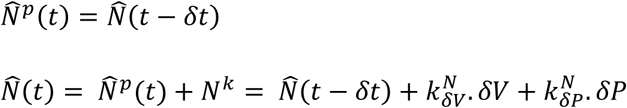

Because 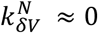 (Table 3), neck position is updated exclusively by its own proprioceptive feedback *δP*. Furthermore, the equation above is transformed into:

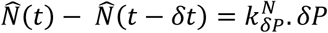

In a steady-state, 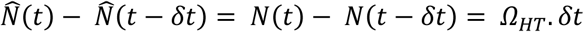, leading to:

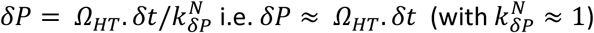

Thus, similar to the reasons already pointed out in section “Kalman feedback gains”, the feedback signal *δP* is scaled by 1/*δt* in Fig. 7 Suppl. 1-3. Also, because the gain 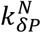 (which is close to 1, see Table 3) doesn’t scale with *δt*, the feedback 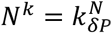. *𝛿P* is also scaled by 1/*δt* in Fig. 7 Suppl. 1-3.

Importantly, the equations above indicate that neck proprioception error should encode neck velocity even when the proprioceptive signals are assumed to encode neck position.

Next, we note that, during passive neck rotation, the estimate of head velocity relative to the trunk is determined by 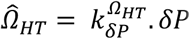. Since *δP ≈ Ω*_*HT*_. *δt*, we expect that 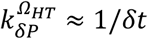. Accordingly, simulations performed with *δt* = 0.01*s* yield 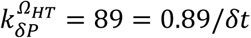. Furthermore, performing simulations with *δt* = 0.1*s* leads to 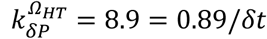. We find that 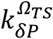 is also dependent of *δt*. Therefore, the corresponding Kalman gains scale with *δt* in Table 3.

Note that the considerations above can equally explain the amplitude and dynamics of the predicted neck position and of the neck proprioceptive error in Fig. 7 Suppl. 6. The simulation in Fig. 7 Suppl. 6C is identical (with half the amplitude) to a passive rotation of the head relative to the trunk (Fig. 7 Suppl. 2), where *δP* = *δV* > 0. The simulation in Fig. 7 Suppl. 6C can be explained mathematically by noticing that it is equivalent to an active head motion (where *δP* = 0) superimposed to a passive rotation of the head with a gain of 0.5 and in the opposite direction, resulting in *δP* = *δV* < 0.

### Three-dimensional Kalman filter

For the sake of simplicity, we have restricted our model to one dimension in this study. However, generalizing the model to three-dimensions may be useful for further studies and is relatively easily accomplished, by (1) replacing one-dimensional variables by three-dimensional vectors and (2) locally linearizing a non-linearity that arises from a vectorial cross-product (eq. 9 below), as shown in this section.

*Principle:* Generalization of the Kalman filter to three dimensions requires replacing each motion and sensory parameter with a 3D vector. For instance, *Ω* is replaced by *Ω*_*x*_, *Ω*_*y*_ and *Ω*_*z*_ that encode the three-dimensional rotation vector in a head-fixed reference frame (*x*, *y*, *z*). Sensory variables are also replaced by three variables, i.e. (*V*_*x*_, *V*_*y*_, *V*_*z*_) and (*F*_*x*_, *F*_*y*_, *F*_*z*_) that encode afferent signals from the canals and otoliths in three dimensions.

With one exception, all variables along one axis (e.g. *Ω*_*x*_, *C*_*x*_, *G*_*x*_, *A*_*x*_, *V*_*x*_, *F*_*x*_ along the *x* axis) are governed by the same set of equations (*eqn* 1 − 6) as the main model. Therefore, the full 3D model can be thought of as three independent Kalman filters operating along the *x*, *y* and *z* dimensions. The only exception is the three-dimensional computation of tilt, which follows the equation:

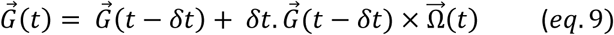

Where 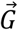 and 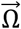 are vectorial representations of (*G*_*x*_, *G*_*y*_, *G*_*z*_) and (*Ω*_*x*_, *Ω*_*y*_, *Ω*_*z*_), and × represents a vectorial cross-product. In matrix form,

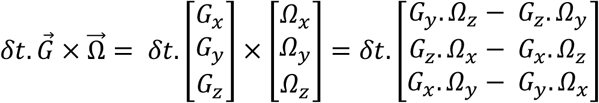

This non-linearity is implemented by placing the terms *δt* and (*G*_*x*_, *G*_*y*_, *G*_*z*_) in the matrices *M* and *E* that integrate rotation inputs (*Ω*^*u*^ and *Ω*^*E*^) into tilt, as shown below.

*Implementation:* The implementation of the 3D Kalman filter is best explained by demonstrating how the matrices of the 1D filter are replaced by scaled-up matrices.

First, we replace all motion variables and inputs by triplets of variables along *x*, *y* and *z*:

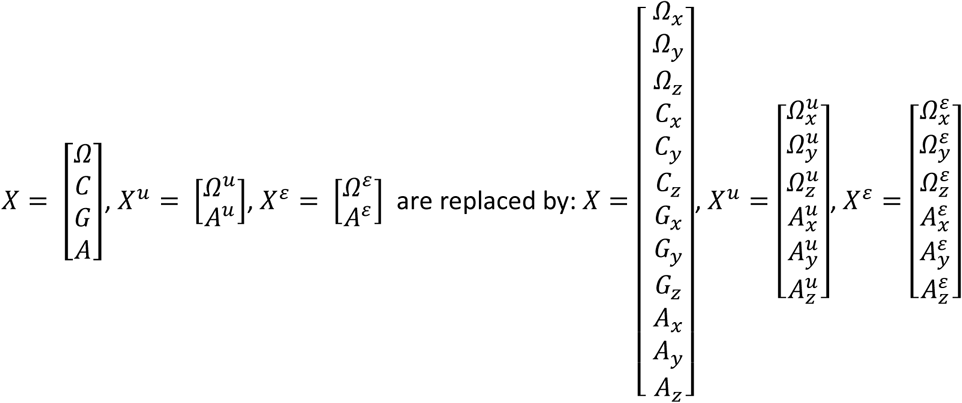

Next, we scale the matrix *D* up; each non-zero element in the 1D version being repeated twice in the 3D version:

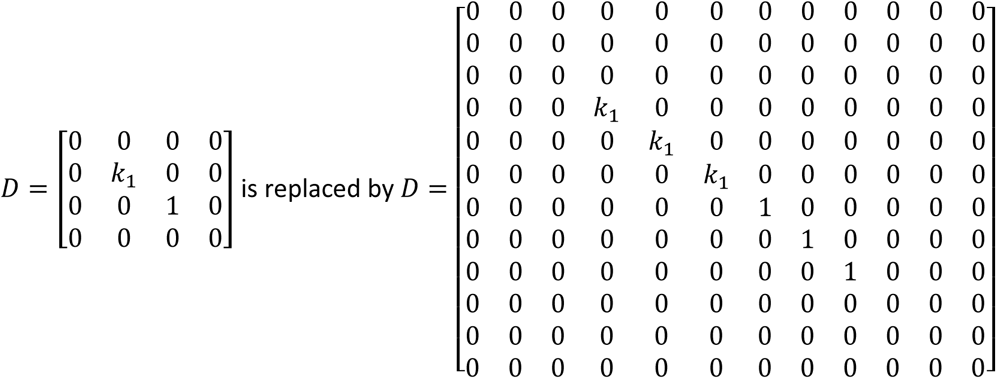

We build *M* in a similar manner. Furthermore, the element *s*. *δt* (that encodes the integration of *Ω*^*u*^ into *G*) is replaced by a set of terms that encode the cross-product in (*eq*. 9). As previously, *E* = *M*. Note that *M* and *E* must be recomputed at each iteration since *G*_*x*_, *G*_*y*_ and *G*_*z*_ change continuously.

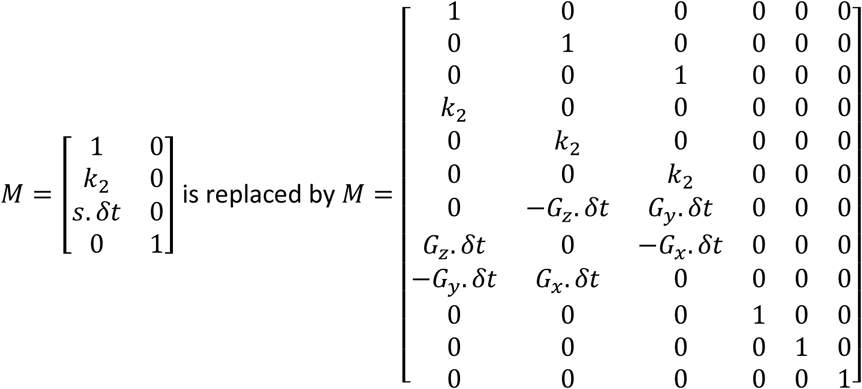

Similarly, we build the sensor model as follows:

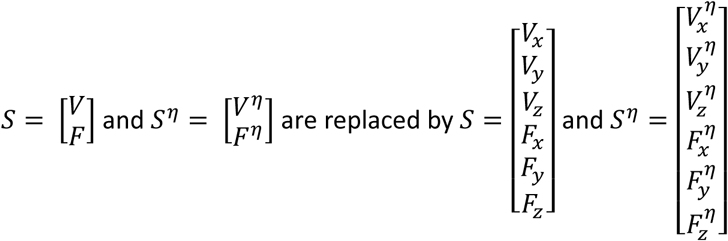

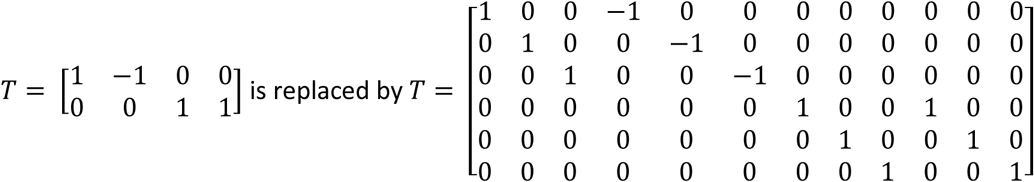

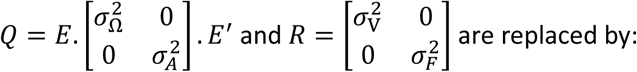

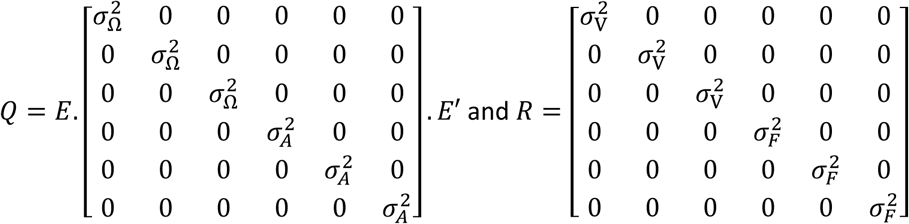

## References

Alpini, D., Botta, M., Mattei, V., & Tornese, D. (2009). Figure ice skating induces vestibulo-ocular adaptation specific to required athletic skills. Sport Sciences for Health, 5(3), 129–134.

Angelaki, D. E., & Hess, B. J. (1995a). Lesion of the nodulus and ventral uvula abolish steady-state off-vertical axis otolith response. Journal of Neurophysiology, 73(4), 1716–1720.

Angelaki, D. E., & Hess, B. J. (1995b). Inertial representation of angular motion in the vestibular system of rhesus monkeys. II. Otolith-controlled transformation that depends on an intact cerebellar nodulus. Journal of Neurophysiology, 73(5), 1729–1751.

Angelaki, D. E., & Hess, B. J. (1996). Three-dimensional organization of otolith-ocular reflexes in rhesus monkeys. II. Inertial detection of angular velocity. Journal of neurophysiology, 75(6), 2425–2440.

Angelaki, D. E., Shaikh, A. G., Green, A. M., & Dickman, J. D. (2004). Neurons compute internal models of the physical laws of motion. Nature, 430(6999), 560–564.

Angelaki, D. E., & Cullen, K. E. (2008). Vestibular system: the many facets of a multimodal sense. Annu. Rev. Neurosci., 31, 125–150.

Brandt, T., Büchele, W., & Arnold, F. (1977). Arthrokinetic nystagmus and ego-motion sensation. Experimental Brain Research, 30(2-3), 331–338.

Bertolini, G., Ramat, S., Laurens, J., Bockisch, C. J., Marti, S., Straumann, D., & Palla, A. (2011). Velocity storage contribution to vestibular self-motion perception in healthy human subjects. Journal of neurophysiology, 105(1), 209–223.

Black, F. O., Shupert, C. L., Horak, F. B., & Nashner, L. M. (1988). Abnormal postural control associated with peripheral vestibular disorders. Progress in brain research, 76, 263–275.

Borah, J., Young, L. R., & Curry, R. E. (1988). Optimal estimator model for human spatial orientation. Annals of the New York Academy of Sciences, 545(1), 51–73.

Brooks, J. X., & Cullen, K. E. (2009). Multimodal integration in rostral fastigial nucleus provides an estimate of trunk movement. Journal of Neuroscience, 29(34), 10499–10511.

Brooks, J. X., & Cullen, K. E. (2013). The primate cerebellum selectively encodes unexpected self-motion. Current Biology, 23(11), 947–955.

Brooks, J. X., & Cullen, K. E. (2014). Early vestibular processing does not discriminate active from passive self-motion if there is a discrepancy between predicted and actual proprioceptive feedback. Journal of neurophysiology, 111(12), 2465–2478.

Brooks, J. X., Carriot, J., & Cullen, K. E. (2015). Learning to expect the unexpected: rapid updating in primate cerebellum during voluntary self-motion. Nature neuroscience, 18(9), 1310–1317.

Carriot, J., Brooks, J. X., & Cullen, K. E. (2013). Multimodal integration of self-motion cues in the vestibular system: active versus passive translations. Journal of neuroscience, 33(50), 19555–19566.

Crawford, J. (1964). Living without a balancing mechanism. The British journal of ophthalmology, 48(7), 357.

Cullen, K. E., & Minor, L. B. (2002). Semicircular canal afferents similarly encode active and passive head-on-trunk rotations: implications for the role of vestibular efference. J Neurosci, 22(11), RC226.

Cullen, K. E. (2012). The vestibular system: multimodal integration and encoding of self-motion for motor control. Trends in neurosciences, 35(3), 185–196.

Cullen, K. E., Brooks, J. X., Jamali, M., Carriot, J., & Massot, C. (2011). Internal models of self-motion: computations that suppress vestibular reafference in early vestibular processing. Experimental brain research, 210(3-4), 377–388.

Dale, A. & Cullen, K.E. (2016) Sensory coding in the vestibular thalamus discriminates passive from active self-motion. Program No. 181.10/JJJ22, Neuroscience Meeting Planner. Society for Neuroscience Meeting, San Diego.

Fusi, S., Miller, E. K., & Rigotti, M. (2016). Why neurons mix: high dimensionality for higher cognition. Current opinion in neurobiology, 37, 66–74.

Gdowski, G. T., & McCrea, R. A. (1999). Integration of vestibular and head movement signals in the vestibular nuclei during whole-trunk rotation. Journal of neurophysiology, 82(1), 436–449.

Gdowski, G. T., Boyle, R., & McCrea, R. A. (2000). Sensory processing in the vestibular nuclei during active head movements. Archives italiennes de biologie, 138(1), 15–28.

Graybiel, A. (1952). Oculogravic illusion. AMA archives of ophthalmology, 48(5), 605–615.

Herdman, S. (1996) Vestibular Rehabilitation. In Baloh, R. W., & Halmagyi, G. M. (Eds.), Disorders of the vestibular system. Oxford University Press, USA, 583–597.

Herdman, S. J., Blatt, P., Schubert, M. C., & Tusa, R. J. (2000). Falls in patients with vestibular deficits. Otology & Neurotology, 21(6), 847–851.

Hess, B. J. M., & Angelaki, D. E. (1993). Angular velocity detection by head movements orthogonal to the plane of rotation. Experimental brain research, 95(1), 77–83.

Horstmann, G. A., & Dietz, V. (1988). The contribution of vestibular input to the stabilization of human posture: a new experimental approach. Neuroscience letters, 95(1), 179–184.

Jamali, M., Sadeghi, S. G., & Cullen, K. E. (2009). Response of vestibular nerve afferents innervating utricle and saccule during passive and active translations. Journal of neurophysiology, 101(1), 141–149.

Kalman, R. E. (1960). A new approach to linear filtering and prediction problems. Journal of basic Engineering, 82(1), 35–45.

Karmali, F., & Merfeld, D. M. (2012). A distributed, dynamic, parallel computational model: the role of noise in velocity storage. Journal of Neurophysiology, 108(2), 390–405.

Kennedy, A., Wayne, G., Kaifosh, P., Alviña, K., Abbott, L. F., & Sawtell, N. B. (2014). A temporal basis for predicting the sensory consequences of motor commands in an electric fish. Nature neuroscience, 17(3), 416–422.

Keshner, E. A., Allum, J. H. J., & Pfaltz, C. R. (1987). Postural coactivation and adaptation in the sway stabilizing responses of normals and patients with bilateral vestibular deficit. Experimental Brain Research, 69(1), 77–92.

Laurens, J. (2006). Modélisation Bayésienne des interactions visuo-vestibulaires (Doctoral dissertation, Paris 6).

Laurens, J., & Droulez, J. (2007). Bayesian processing of vestibular information. Biological cybernetics, 96(4), 389–404.

Laurens, J., & Droulez, J. (2008). Bayesian modelling of visuo-vestibular interactions. In Probabilistic reasoning and decision making in sensory-motor systems (pp. 279–300). Springer Berlin Heidelberg.

Laurens, J., Straumann, D., & Hess, B. J. (2010). Processing of angular motion and gravity information through an internal model. Journal of neurophysiology, 104(3), 1370–1381.

Laurens, J., Strauman, D., & Hess, B. J. (2011a). Spinning versus wobbling: how the brain solves a geometry problem. Journal of Neuroscience, 31(22), 8093–8101.

Laurens, J., Valko, Y., & Straumann, D. (2011b). Experimental parameter estimation of a visuo-vestibular interaction model in humans. Journal of Vestibular Research, 21(5), 251–266.

Laurens, J., & Angelaki, D. E. (2011). The functional significance of velocity storage and its dependence on gravity. Experimental brain research, 210(3-4), 407–422.

Laurens, J., Meng, H., & Angelaki, D. E. (2013a). Computation of linear acceleration through an internal model in the macaque cerebellum. Nature neuroscience, 16(11), 1701–1708.

Laurens, J., Meng, H., & Angelaki, D. E. (2013b). Neural representation of orientation relative to gravity in the macaque cerebellum. Neuron, 80(6), 1508–1518.

Laurens, J., & Angelaki, D. E. (2015). How the Vestibulocerebellum Builds an Internal Model of Self-motion. The Neuronal Codes of the Cerebellum, 97.

Laurens, J., Liu, S., Yu, X. J., Chan, R., Dickman, D., DeAngelis, G. C., & Angelaki, D. E. (2017). Transformation of spatiotemporal dynamics in the macaque vestibular system from otolith afferents to cortex. eLife, 6, e20787.

Lee, R. X., Huang, J. J., Huang, C., Tsai, M. L., & Yen, C. T. (2015). Plasticity of cerebellar Purkinje cells in behavioral training of trunk balance control. Frontiers in systems neuroscience, 9.

Lim, K., Karmali, F., Nicoucar, K., & Merfeld, D. M. (2017). Perceptual precision of passive trunk tilt is consistent with statistically optimal cue integration. Journal of Neurophysiology, jn-00073.

Mackrous, I., Carriot, J., Jamali, M., Brooks, J, & Cullen, K.E. (2016) Selective encoding of unexpected head tilt by the central neurons takes into account the cerebellar computation output. Program No. 718.04/QQ2, Neuroscience Meeting Planner. Society for Neuroscience Meeting, San Diego.

MacNeilage, P. R., Ganesan, N., & Angelaki, D. E. (2008). Computational approaches to spatial orientation: from transfer functions to dynamic Bayesian inference. Journal of neurophysiology, 100(6), 2981–2996.

Marlinski, V., & McCrea, R. A. (2008). Coding of self-motion signals in ventro-posterior thalamus neurons in the alert squirrel monkey. Experimental brain research, 189(4), 463.

Marlinski, V., & McCrea, R. A. (2009). Self-motion signals in vestibular nuclei neurons projecting to the thalamus in the alert squirrel monkey. Journal of neurophysiology, 101(4), 1730–1741.

Mayne, R. (1974). A systems concept of the vestibular organs. In Vestibular system part 2: psychophysics, applied aspects and general interpretations (pp. 493–580). Springer Berlin Heidelberg.

McCrea, R. A., Gdowski, G. T., Boyle, R., & Belton, T. (1999). Firing behavior of vestibular neurons during active and passive head movements: vestibulo-spinal and other non-eye-movement related neurons. Journal of neurophysiology, 82(1), 416–428.

McCrea, R. A., & Luan, H. (2003). Signal processing of semicircular canal and otolith signals in the vestibular nuclei during passive and active head movements. Annals of the New York Academy of Sciences, 1004(1), 169–182.

Meng, H., May, P. J., Dickman, J. D., & Angelaki, D. E. (2007). Vestibular signals in primate thalamus: properties and origins. Journal of Neuroscience, 27(50), 13590–13602.

Meng, H., & Angelaki, D. E. (2010). Responses of ventral posterior thalamus neurons to three-dimensional vestibular and optic flow stimulation. Journal of neurophysiology, 103(2), 817–826.

Merfeld, D. M. (1995). Modeling the vestibulo-ocular reflex of the squirrel monkey during eccentric rotation and roll tilt. Experimental Brain Research, 106(1), 123–134.

Merfeld, D. M., Zupan, L., & Peterka, R. J. (1999). Humans use internal models to estimate gravity and linear acceleration. Nature, 398(6728), 615–618.

Mergner, T., Siebold, C., Schweigart, G., & Becker, W. (1991). Human perception of horizontal trunk and head rotation in space during vestibular and neck stimulation. Experimental Brain Research, 85(2), 389–404.

Oman, C. M. (1982). A heuristic mathematical model for the dynamics of sensory conflict and motion sickness. Acta Oto-Laryngologica, 94(sup392), 4–44.

Paige, G. D., & Seidman, S. H. (1999). Characteristics of the VOR in response to linear acceleration. Annals of the New York Academy of Sciences, 871(1), 123–135.

Raphan, T., Matsuo, V., & Cohen, B. (1979). Velocity storage in the vestibulo-ocular reflex arc (VOR). Experimental Brain Research, 35(2), 229–248.

Reisine, H., & Raphan, T. (1992). Neural basis for eye velocity generation in the vestibular nuclei of alert monkeys during off-vertical axis rotation. Experimental brain research, 92(2), 209–226.

Requarth, T., & Sawtell, N. B. (2011). Neural mechanisms for filtering self-generated sensory signals in cerebellum-like circuits. Current opinion in neurobiology, 21(4), 602–608.

Rigotti, M., Barak, O., Warden, M. R., Wang, X. J., Daw, N. D., Miller, E. K., & Fusi, S. (2013). The importance of mixed selectivity in complex cognitive tasks. Nature, 497(7451), 585–590.

Riley, N. H. (2010). Neuromuscular adaptations during perturbations in individuals with and without bilateral vestibular loss. (Doctoral dissertation, University of Iowa)

Robinson, D. A. (1977). Vestibular and optokinetic symbiosis: an example of explaining by modelling. Control of gaze by brain stem neurons. Elsevier, Amsterdam, 49–58.

Roy, J. E., & Cullen, K. E. (2001). Selective processing of vestibular reafference during self-generated head motion. Journal of Neuroscience, 21(6), 2131–2142.

Roy, J. E., & Cullen, K. E. (2004). Dissociating self-generated from passively applied head motion: neural mechanisms in the vestibular nuclei. Journal of Neuroscience, 24(9), 2102–2111.

Sadeghi, Soroush G., Lloyd B. Minor, and Kathleen E. Cullen. “Response of vestibular-nerve afferents to active and passive rotations under normal conditions and after unilateral labyrinthectomy.” Journal of neurophysiology 97.2 (2007): 1503–1514.

Shaikh, A. G., Ghasia, F. F., Dickman, J. D., & Angelaki, D. E. (2005). Properties of cerebellar fastigial neurons during translation, rotation, and eye movements. Journal of neurophysiology, 93(2), 853–863.

Shadmehr, R., Smith, M. A., & Krakauer, J. W. (2010). Error correction, sensory prediction, and adaptation in motor control. Annual review of neuroscience, 33, 89–108.

Tanguy, S., Quarck, G., Etard, O., Gauthier, A., & Denise, P. (2008). Vestibulo-ocular reflex and motion sickness in figure skaters. European journal of applied physiology, 104(6), 1031.

Tseng, Y. W., Diedrichsen, J., Krakauer, J. W., Shadmehr, R., & Bastian, A. J. (2007). Sensory prediction errors drive cerebellum-dependent adaptation of reaching. Journal of neurophysiology, 98(1), 54–62.

Waespe, W., & Henn, V. (1977). Neuronal activity in the vestibular nuclei of the alert monkey during vestibular and optokinetic stimulation. Experimental Brain Research, 27(5), 523–538.

Waespe, W., Cohen, B., & Raphan, T. (1983). Role of the flocculus and paraflocculus in optokinetic nystagmus and visual-vestibular interactions: effects of lesions. Experimental Brain Research, 50(1), 9–33.

Wearne, S., Raphan, T., & Cohen, B. (1998). Control of spatial orientation of the angular vestibuloocular reflex by the nodulus and uvula. Journal of neurophysiology, 79(5), 2690–2715.

Wolpert, D. M., Ghahramani, Z., & Jordan, M. I. (1995). An internal model for sensorimotor integration. Science, 269(5232), 1880.

Yakusheva, T. A., Shaikh, A. G., Green, A. M., Blazquez, P. M., Dickman, J. D., & Angelaki, D. E. (2007). Purkinje cells in posterior cerebellar vermis encode motion in an inertial reference frame. Neuron, 54(6), 973–985.

Yakusheva, T., Blazquez, P. M., & Angelaki, D. E. (2008). Frequency-selective coding of translation and tilt in macaque cerebellar nodulus and uvula. Journal of Neuroscience, 28(40), 9997–10009.

Yakusheva, T. A., Blazquez, P. M., Chen, A., & Angelaki, D. E. (2013). Spatiotemporal properties of optic flow and vestibular tuning in the cerebellar nodulus and uvula. Journal of Neuroscience, 33(38), 15145–15160.

Zupan, L. H., Merfeld, D. M., & Darlot, C. (2002). Using sensory weighting to model the influence of canal, otolith and visual cues on spatial orientation and eye movements. Biological cybernetics, 86(3), 209–230.

